# Florigen-producing cells express FPF1-LIKE PROTEIN 1 that accelerates flowering and stem growth in long days with sunlight red/far-red ratio in *Arabidopsis*

**DOI:** 10.1101/2024.04.26.591289

**Authors:** Hiroshi Takagi, Nayoung Lee, Andrew K. Hempton, Savita Purushwani, Michitaka Notaguchi, Kota Yamauchi, Kazumasa Shirai, Yaichi Kawakatsu, Susumu Uehara, William G. Albers, Benjamin L. R. Downing, Shogo Ito, Takamasa Suzuki, Takakazu Matsuura, Izumi C. Mori, Nobutaka Mitsuda, Daisuke Kurihara, Tomonao Matsushita, Young Hun Song, Yoshikatsu Sato, Mika Nomoto, Yasuomi Tada, Kousuke Hanada, Josh T. Cuperus, Christine Queitsch, Takato Imaizumi

**Affiliations:** Department of Biology, University of Washington, Seattle, Washington, 98195-1800, USA; Center for Gene Research, Nagoya University, Nagoya, 464-8602, Japan; Plant Genomics and Breeding Institute, Seoul National University, Seoul, 08826, Korea; Bioscience and Biotechnology Center, Nagoya University, Nagoya, 464-8601, Japan; Department of Botany, Graduate School of Science, Kyoto University, Kyoto, 606-8502, Japan; Department of Bioscience and Bioinformatics, Kyushu Institute of Technology, Iizuka, 820-8502, Japan; Department of Biological Chemistry, College of Bioscience and Biotechnology, Chubu University, Kasugai, 487-8501, Japan; Institute of Plant Science and Resources, Okayama University, Kurashiki, 710-0046, Japan; Bioproduction Research Institute, National Institute of Advanced Industrial Science and Technology (AIST), Tsukuba, 305-8566, Japan; Institute of Transformative Bio-Molecules (WPI-ITbM), Nagoya University, Nagoya, 464-8601, Japan; Institute for Advanced Research (IAR), Nagoya University, Nagoya, 464-8601, Japan; Department of Agricultural Biotechnology, Seoul National University, Seoul, 08826, Korea; Division of Biological Science, Graduate School of Science, Nagoya University, Nagoya, 464-8602, Japan; Department of Genome Sciences, University of Washington, Seattle, Washington, 98195-5065, USA; Brotman Baty Institute for Precision Medicine, University of Washington, Seattle, Washington, 98195-8047, USA

**Keywords:** photoperiodic flowering, florigen, stem growth, natural long days, FT, CO, FPF1 *Arabidopsis*, gibberellins, auxin

## Abstract

Seasonal changes in spring induce flowering by expressing the florigen, FLOWERING LOCUS T (FT), in *Arabidopsis*. *FT* is expressed in unique phloem companion cells with unknown characteristics. The question of which genes are co-expressed with *FT* and whether they have roles in flowering remains elusive. Through tissue-specific translatome analysis, we discovered that under long-day conditions with the natural sunlight red/far-red ratio, the *FT*-producing cells express a gene encoding FPF1-LIKE PROTEIN 1 (FLP1). The master *FT* regulator, CONSTANS (CO), controls *FLP1* expression, suggesting *FLP1*’s involvement in the photoperiod pathway. FLP1 promotes early flowering independently of *FT,* is active in the shoot apical meristem, and induces the expression of *SEPALLATA 3* (*SEP3*), a key E-class homeotic gene. Unlike FT, FLP1 facilitates inflorescence stem elongation. Our cumulative evidence indicates that FLP1 may act as a mobile signal. Thus, FLP1 orchestrates floral initiation together with FT and promotes inflorescence stem elongation during reproductive transitions.

## Introduction

The precise regulation of flowering timing is essential for plants’ success in reproduction and survival in natural environments. Seasonal environmental factors, including light and temperature, are integrated as signals to coordinate the timing of flowering predominately through the function of the florigen gene, *FT*.^1,2^ FT protein acts as a systemic flowering signal widely conserved in flowering plants.^1,2^ Under the standard laboratory long-day (LD) conditions, *FT* exhibits a single evening peak in *Arabidopsis*. Therefore, *FT* regulating mechanisms have been discussed based on the expression levels of *FT* in the evening. However, our recent study in outdoor LD conditions (where natural sunlight is the sole light source) revealed the presence of a bimodal expression pattern of *FT* with morning and evening peaks.^3,4^ The morning *FT* expression is often the major peak in plants grown outside because the evening peak is attenuated by the temperature differences throughout the day.^3^ The emergence of the *FT* morning peak is attributed to the abundant far-red light present in the sunlight, which is low in laboratory light sources. The red/far-red light (R/FR) ratio in natural light conditions is approximately 1.0, whereas it is typically greater than 2.0 in laboratory conditions with artificial light sources. This sunlight level R/FR ratio induces *FT* expression in the morning through the stabilization of CONSTANS (CO), the master regulator of *FT*, and phytochrome A-mediated FR high-irradiance response.^3,4^

Following local FT protein synthesis in the leaf phloem companion cells, FT is transported to the shoot apical meristem (SAM).^5^ FT physically interacts with FD, the SAM-specific bZIP transcription factor (TF), with the assistance of a 14-3-3 protein.^6,7^ This florigen activation complex (FAC) induces floral integrator genes, such as *LEAFY* (*LFY*) and *APETALA1* (*AP1*), to initiate inflorescent development.^8^ In the ABCE floral organ development model, the combinatorial actions of floral homeotic genes specify identities of sepals, petals, stamens, and carpels. *LFY* and A-class gene *AP1* start expressing during the early stage of inflorescence meristem formation and boost each other’s expression through positive feedback loop regulation.^9,10^ AP1 subsequently upregulates the major E-class gene *SEPALLATA 3* (*SEP3*) through both direct binding on the *SEP3* promoter and downregulation of repressive floral genes such as *SHORT VEGETATIVE PHASE* (*SVP*).^9,11^ SEP3 is a crucial player in plant flowering because it not only triggers the expression of B- and C-class genes but is also involved in the formation of all four-type floral organs through the physical interaction with A-, B-, and C-class homeotic proteins.^11,12^ The induction of *LFY* and *AP1* mediated by FT–FD interaction is currently considered the critical opening event in floral organ formation. However, while AP1 induces *SEP3*, SEP3 also directly triggers the expression of other homeotic genes, including *AP1,*^13^ indicating the positive feedback interaction between *AP1* and *SEP3*. Consistently, like in the case of *AP1*, constitutive expression of *SEP3* is sufficient to induce the early flowering phenotype.^14^

FLOWERING PROMOTING FACTOR 1 (FPF1), another factor implicated in *Arabidopsis* floral transition, is a small 12.6-kDa protein with no known functional domain, expressed primarily in inflorescence and floral meristems.^15,16^ Although the loss-of-function mutant phenotype has not yet been characterized, constitutive expression of *FPF1* confers early flowering phenotypes.^15,16^ In contrast to the well-studied FT-dependent flowering pathway described above, the genetic pathway underlying the FPF1-dependent flowering induction remains elusive. The previous genetic study showed that *FPF1* synergistically works with *LFY* and *AP1,* downstream components of FT, on flowering,^16^ suggesting that the mode of action of FPF1 is different from FT. However, it has not yet experimentally been shown whether *FPF1*’s flowering-promoting function is independent of *FT* in *Arabidopsis*. Moreover, FPF1 appears to integrate with gibberellin signaling. Severe reduction of endogenous gibberellin levels, either by the *ga2-1* mutation or treatment of paclobutrazol (an inhibitor of gibberellin biosynthesis), reverted the early flowering phenotype induced by *FPF1* overexpression,^15^ suggesting that FPF1 elicits flowering induction through the gibberellin signaling. In addition to the flowering acceleration, when FPF1 is ectopically over-expressed in seedlings, it promotes hypocotyl elongation through cell size expansion, indicating a distinct function from FT. Recent studies also revealed that rice (*Oryza sativa*) and tomato (*Solanum lycopersicum*) homologs of *FPF1* (known in rice as *ACCELERATOR OF INTERNODE ELONGATION 1*; *ACE1*) promote stem internode elongation but not flowering induction, suggesting that FPF1’s function in tissue elongation is widely conserved among plant species, but it does not always accompany with acceleration of flowering.^17,18^

Whereas the genetic pathway and interaction responsible for *FPF1*-dependent phenotypes are largely inconclusive, a recent study using *Brachypodium distachyon* revealed that FPF1 family proteins function as repressors of TFs through physical interaction.^19^ In contrast to *Arabidopsis FPF1*, two *FPF1* homologs of *Brachypodium, FPL1* and *FPL7*, delay flowering. These FPF1 homologs prevent FAC formation and attenuate the DNA binding activity of FD1, an FD homolog of *Brachypodium*, through protein-protein interactions.^19^ Thus, it is possible that flowering promotion by FPF1 may also be mediated by the direct interaction with specific TFs.

In *Arabidopsis*, *FT* gene expression is restricted to specific phloem companion cells located in the distal parts of a leaf.^20,21^ Despite decades of research, the question of how florigen-producing cells differ from the rest of the companion cells remains unresolved.^20,22^ These florigen-producing cells serve as the sites for integrating seasonal cues and transmitting signals to initiate and maintain floral transitions. Thus, understanding the unique characteristics of these cells is essential for unraveling regulatory strategies in plant development and for understanding signal integrations of seasonal information. Given the unique spatial distribution of *FT*-producing cells among phloem companion cells,^20^ we employed tissue/cell-specific translatome analysis, Translating Ribosome Affinity Purification (TRAP)-seq,^23^ to elucidate the unique characteristics of *FT*-producing cells. It revealed that *FT*-producing cells in the leaf vasculature may be developing phloem companion cells with active transport and that some known *FT*-regulating TFs are enriched in these cells. In addition, our translatome analysis demonstrated that, under LD conditions with natural R/FR, *FPF1-LIKE PROTEIN 1* (*FLP1*), a homologous gene of *FPF1*, is highly expressed in the *FT*-producing cells. We found that FLP1 not only accelerates flowering in an *FT*-independent manner but also promotes inflorescence stem elongation. FLP1 induces early flowering when ectopically expressed in the SAM and may function as a systemic signaling protein. Thus, our study uncovered that phloem companion cells specialized for seasonal sensing supply multiple small (and possibly mobile) proteins to orchestrate growth and development during reproductive transitions.

## Results

### Tissue-specific translatome analysis of florigen-producing cells

To investigate the characteristics of *FT*-producing cells, we performed tissue/cell-specific translatome (TRAP-seq) analysis, as translatome more closely reflects protein abundance than transcriptome.^24^ In the TRAP-seq analysis, epitope-tagged RIBOSOMAL PROTEIN L18 (RPL18) controlled by tissue/cell-specific promoters are co-immunoprecipitated with translating mRNAs attached to the tagged RPL18-containing ribosomes. To analyze which transcript is translated in *FT*-producing cells, we generated the *pFT:FLAG-GFP-RPL18* line. To express the *FLAG-GFP-RPL18* gene in the same way as *FT* is expressed, the *pFT:FLAG-GFP-RPL18* construct possesses not only the 5.7 kb upstream *FT* promoter region containing the entire upstream enhancer region “Block C” but also has the genomic sequences spanning from the *FT* gene to the 3’ Block E enhancer region inserted after the stop codon of *FLAG-GFP-RPL18* (Figures 1A and S1A).^22,25^ To simulate *FT* expression in natural light environments, plants were grown under LD conditions with the R/FR ratio adjusted to natural sunlight (R/FR = approximately 1.0, referred to as LD+FR) alongside conventional lab LD conditions (R/FR > 2.0). The *pFT:FLAG-GFP-RPL18* plants grown under LD+FR showed GFP signals in a part of cotyledon and true leaf vasculatures (Figures S1B and S1C), consistent with the spatial expression pattern of *FT*. Moreover, the levels of *FLAG-GFP-RPL18* transcripts showed uptrends at both Zeitgeber Time 4 (ZT4) and ZT16 under LD+FR conditions compared with those in LD conditions (Figure S1D), indicating our construct captures *FT* regulation controlled by R/FR ratio changes. To elucidate the translatome profiles unique to the *FT*-producing companion cells, we included previously established TRAP-seq lines targeting phloem companion cells (*pSUC2*), mesophyll cells (*pRBCS1A*), epidermal cells (*pCER5*), and whole tissues (*p35S*) as comparison (Figure 1A).^23^ Although the activity of the *35S* promoter is reportedly weaker in reproductive tissues such as bolted stems and inflorescences compared with other tissues, it is stably highly active in whole tissues of cotyledons, young true leaves, and roots at the vegetative stages,^26,27^ thus, we exploited the *p35S:FLAG-GFP-RPL18* line as a whole tissue control for our analysis. Due to the depressed expression levels of the *FLAG-GFP-RPL18* gene in the *pCER5:FLAG-GFP-RPL18* line obtained from the ABRC stockcenter, we used the *pCER5:FLAG-RPL18* line for the epidermal cell translatome. Previous TRAP RNA purification analyses utilized FLAG-RPL18 protein both with and without GFP, ^23,28^ indicating that the presence or absence of GFP with RPL18 does not affect the isolation of translating mRNA. Samples were collected at ZT4, a timepoint four hours after the onset of light, corresponding to the timing of the morning peak of *FT* in LD+FR.^4^

**Figure 1.**
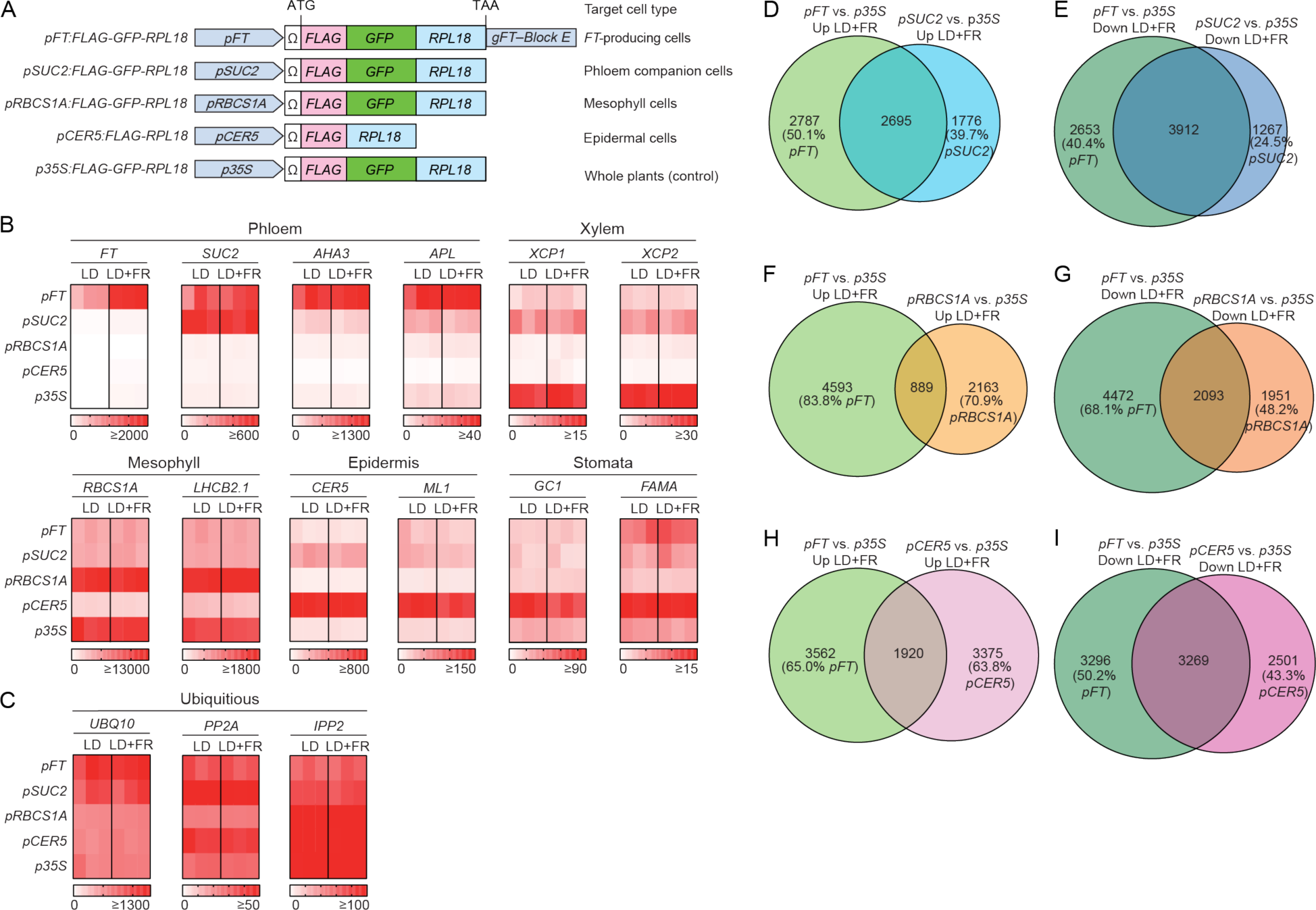
The translatome profiling reveals the *FT*-producing cells are similar to the *SUC2*-expressing phloem companion cells but express unique transcripts. (A) A schematic diagram of constructs equipped with tissue-specific promoters and *FLAG-GFP-RPL18* used in this study. It is reported that with or without the inclusion of *GFP* coding sequences in the *RPL18* construct does not affect the efficiency of TRAP. (B and C) Unique translational patterns of tissue/cell-specific marker transcripts in our TRAP-seq datasets. The tissue/cell-specific TRAP lines are indicated on the left, and the growth conditions are at the top. Plants were grown under LD and LD+FR conditions. Color gradation reflects RPKM values depicted below for each gene. The results from 3 biological replicates are shown. The translated transcript levels of tissue-specific marker genes and typical qRT-PCR reference genes are shown in (B) and (C), respectively. (D–I) Quantitative Venn diagrams showing overlaps between translatome datasets of *pFT:FLAG-GFP-RPL18* and other tissue-specific lines. Up-regulated genes (D, F, and H) and down-regulated genes (E, G, and I) in comparison with *p35S:FLAG-GFP-RPL18* under LD+FR conditions. The percentages of the translated genes in each unique portion of the diagrams are shown.

First, we assessed whether our tissue/cell-specific translatome datasets could enrich known marker gene transcripts. The normalized readcounts, Reads Per Kilobase per Million mapped reads (RPKM), of tissue/cell-specific genes driving the tagged *RPL18* gene (*FT*, *SUC2*, *RBCS1A*, and *CER5*) and other marker genes (*AHA3* and *APL* for phloem; *XCL1* and *XCL2* for xylem; *LHCB2.1* for mesophyll cells; *ML1* for epidermis; *GC1* and *FAMA* for stomata) were specifically increased in their corresponding lines, while typical reference genes in qRT-PCR analysis (*UBQ10*, *PP2A*, and *IPP2*) were ubiquitously found in all tissues (Figures 1B and C), confirming the unique tissue-specific enrichments of isolated mRNAs from each TRAP line. *FT* transcripts exhibited higher translation levels in LD+FR compared to LD (Figure 1B), reflecting the induction of the *FT* morning peak in LD+FR. Neither *pFT:FLAG-GFP-RPL18* nor *pSUC2:FLAG-GFP-RPL18* lines enriched xylem marker genes, *XCP1* and *XCP2*, indicating that our TRAP-seq analysis provided high-resolution datasets that separated different cell types in vasculature tissues. Moreover, between *pFT:FLAG-GFP-RPL18* and *pSUC2:FLAG-GFP-RPL18* datasets, some phloem markers, *AHA3* and *APL*, were further enriched in the *FT*-producing cells (Figure 1B), highlighting the unique characteristics of specific phloem tissues expressing *FT*. To gain a comprehensive understanding of the variations in translated mRNA levels among the examined tissues, we compared the RPKM values of transcripts detected in each line with those of the control *p35S:FLAG-GFP-RPL18* line under LD and LD+FR conditions. This analysis revealed that there were larger numbers of differentially expressed genes (DEGs) existing in *pFT*, *pSUC2*, and *pCER5*-driven lines than in *pRBCS1A*-driven lines (Figures S2A–H; Data S1), indicating the uniqueness of phloem and epidermal tissues compared with photosynthetic ground tissues. Next, to holistically analyze unique translatome profiles in *FT*-producing cells, we asked how many DEGs overlapped between *pFT* and other tissue-specific lines. Overall, the DEGs in *pFT:FLAG-GFP-RPL18* mostly overlapped with (but were not entirely contained within) the DEGs in the phloem companion cell-targeting *pSUC2:FLAG-GFP-RPL18* and overlapped least with those in the mesophyll cell-enriched *pRBCS1A:FLAG-GFP-RPL18* in both LD and LD+FR (Figures 1D–I and S2I–N). These findings indicate that *FT*-producing cells are closely related to *SUC2-*expressing phloem companion cells; however, they also have a unique identity.

To gain more insight into the unique characteristics of *FT*-producing cells, we next identified genes with tissue-specific enrichment patterns similar to *FT* using gene clustering analysis (Figure 2A and Data S2).^29^ For this analysis, we applied log_2_ fold-changes between four tissue/cell-specific *RPL18* lines and control *p35S:FLAG-GFP-RPL18* line grown under either LD or LD+FR conditions. These eight combinations (four different lines with two growth conditions) were used to separate genes exhibiting similar tissue/cell-specific enrichment patterns. We divided our translatome data into 12 groups in the cluster analysis (Figure 2A). *FT*, *SUC2*, *RBCS1A*, and *CER5* genes were grouped into clusters 3 (388 genes), 12 (368 genes), 2 (1,007 genes), and 8 (1,424 genes), respectively. Terms enrichment analysis with Metascape^30^ showed that clusters 2 and 8 were enriched in genes related to photosynthesis and wax/cutin biosynthesis, respectively, consistent with the functions associated with *RBCS1A* and *CER5* (Figures 2B and D). The *SUC2* gene in cluster 12 co-expressed with many *EXTENSIN* (*EXT*) genes (*EXT6*, *8*, *7*, *9*, *10*, *13*, *15*, *16* and *17*) encoding cell wall proteins (Figure 2F; Data S2).

**Figure 2.**
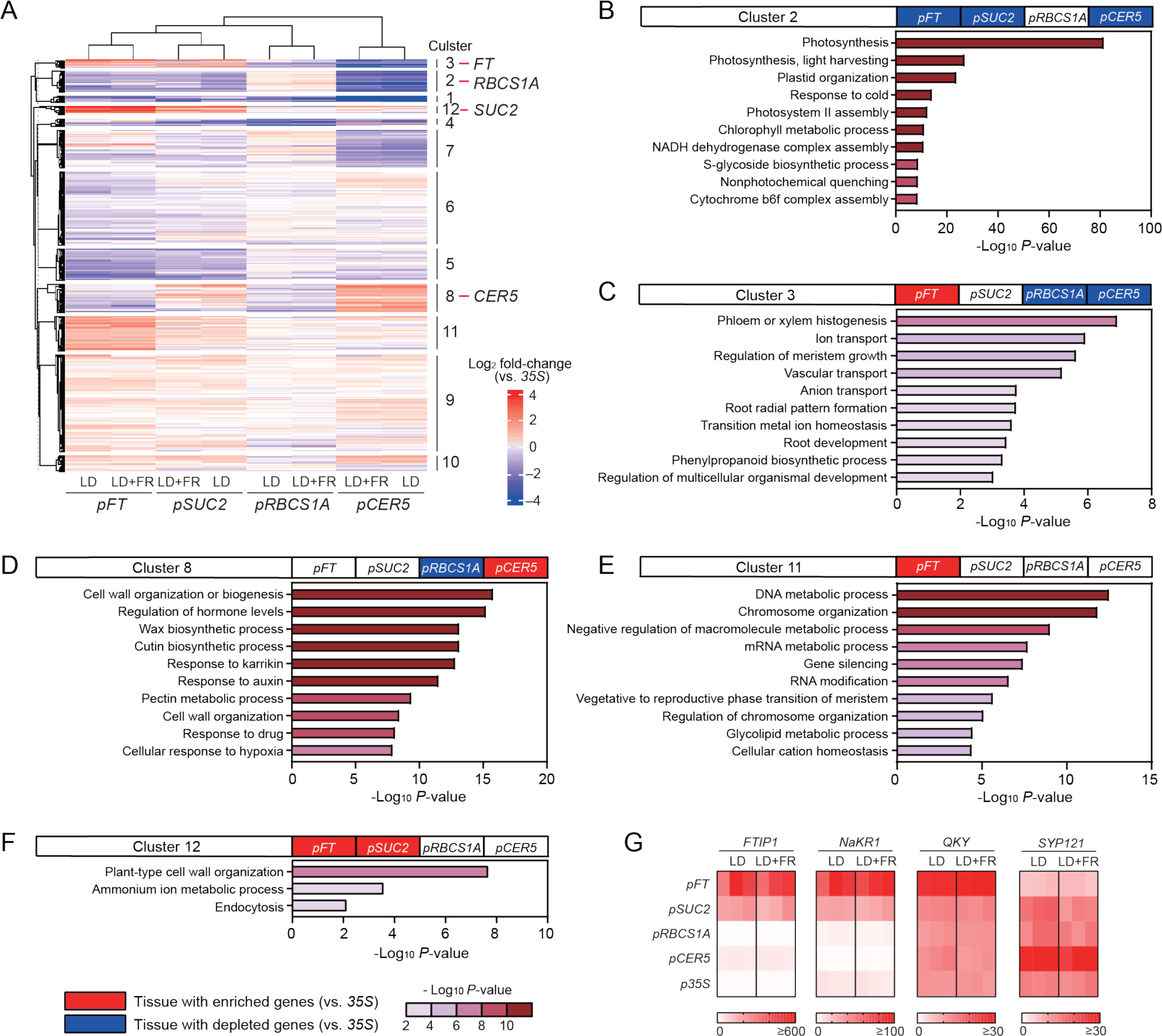
The FT-producing cells are active in transport and nucleic acid metabolism. (A) Heatmap clustering of the TRAP-seq data. The color indicates log_2_ fold-change compared to the *p35S:FLAG-GFP-RPL18* data. The tissue/cell-specific TRAP lines and the growth conditions are indicated at the bottom of the heatmap. Clusters including *FT* (Cluster 3, 388 genes), *RBCS1A* (Cluster 2, 1007 genes), *SUC2* (Cluster 12, 368 genes), and *CER5* (Cluster 8, 1424 genes) are indicated. (B–F) The top 10 enriched terms in clusters (2, 3, 8, 11, and 12) in the heatmap shown in (A). Enrichment analysis was conducted using Metascape. *pFT*, *pSUC2*, *pRBCS1A*, and *pCER5* on top indicate the transgenic lines for TRAP-seq analysis; *pFT:FLAG-RPL18, pSUC2:FLAG-GFP-RPL18, pRBCS1A:FLAG-GFP-RPL18*, and *pCER5:FLAG-RPL18*, respectively. Red and blue colors on the boxes indicating each TRAP-dataset (*pFT*, *pSUC2*, *pRBCS1A*, and *pCER5*) denote that more than 50% of genes are significantly up- or down-regulated (enriched or depleted) in the target tissues compared to the *p35S:FLAG-GFP-RPL18* dataset. The size and color of the bars indicate negative logarithm of the *P*-values. The cluster 12 genes were associated with only 3 statistically enriched terms. (G) Unique translational patterns of FT protein-transporting genes in our TRAP-seq datasets. The tissue/cell-specific TRAP lines are indicated on the left, and the growth conditions are at the top. Plants were grown under LD and LD+FR conditions. Color gradation reflects RPKM values depicted below for each gene. The results from 3 biological replicates are shown.

The emergence of *EXT* genes is evolutionally associated with plant vascularization.^31,32^ Thus, possible roles of EXTs in vascular development are under discussion, although their spatial expression patterns are still largely uncharacterized. Cluster 3, which contains *FT*, showed strong enrichment in terms related to phloem histogenesis, meristem growth regulation, and vascular/ion transport (Figure 2C), suggesting that *FT*-producing cells may be differentiating phloem companion cells with high transport activity. Among the transporters, FT transporter genes *NaKR1*, *FTIP1*, and *QKY* were highly translated in *FT*-producing companion cells (Figure 2G),^33–35^ suggesting that FT protein synthesis and transport are co-regulated. Cluster 11 also included genes highly translated in *pFT:FLAG-GFP-RPL18*. This cluster enriched genes related to DNA and mRNA metabolic processes, chromosome organization, gene silencing, etc. (Figure 2E), implying that gene transcription and translation are actively regulated in the *FT*-producing cells. These results imply that *FT*-producing cells may be differentiating phloem companion cells that are high in metabolic and transport activities within the phloem tissues.

In addition to identifying the molecular function linked to the *FT*-producing cells, our TRAP-seq datasets could be utilized to classify already known *FT* regulators from the point of tissue/cell-specific expression patterns. Although numerous *FT*-regulating transcription factors (TFs) have been identified through genetic studies,^3^ whether any of these TF transcripts are enriched in *FT*- producing cells is not known. To address this question, we compared the translational levels of known *FT*-regulating TFs across different tissues/cells using our TRAP-seq dataset to assess their potential tissue/cell-specific involvement in *FT* expression under LD+FR conditions (Figures S3 and S4). Although many of the known positive *FT* regulators showed similar translated mRNA levels across all tested tissues, *CO* and some CO-interacting regulators (*NF- YB2*, *NF-YC3*, *APL*, *AS1*, *VOZ1,* and *CIB4*) were particularly enriched in *FT*-producing cells (Figure S3). On the other hand, the levels of negative *FT* regulators varied across all tissue/cell types examined (Figure S4). Some negative *FT* regulators (*FLC, MAF2, CDF5, COL9, HHO4*, etc.) were particularly enriched in *FT*-producing cells, while specific TF families (*DELLA*, *AP2- like*, and *bHLH*) were enriched in other cell types such as epidermal cells (Figure S4). These results suggest that certain negative *FT* regulators may control *FT* levels in *FT*-producing cells, while other negative regulators may be involved in preventing *FT* expression in different cell types (such as mesophyll and epidermal cells) to restrict the spatial *FT* expression patterns.

### The *FT*-producing cells express *FLP1*

With tissue-specific translatome datasets, we also aimed to identify a novel regulator of morning *FT* induction and developmentally relevant genes co-expressed with *FT,* which might have been overlooked by common usage of the artificial light (R/FR >2.0) growth conditions. To explore this possibility, we analyzed translational changes in specific tissues/cells, particularly *FT*- producing cells, upon adjustment of the R/FR ratio (Data S3). As expected, adjusting the R/FR ratio significantly increased the translational levels of *FT* transcripts in all TRAP lines except the *pCER5:FLAG-RPL18* line (Figures 3A and S5). However, only a small number of genes were significantly upregulated by the R/FR adjustment across all examined tissues and cell types. In *pFT:FLAG-GFP-RPL18*, unlike FR-marker genes such as *PIL1* and *HFR1*, none of the known *FT*-regulating TFs were strongly enriched by the light treatment (Figure 3A and Data S3).

**Figure 3.**
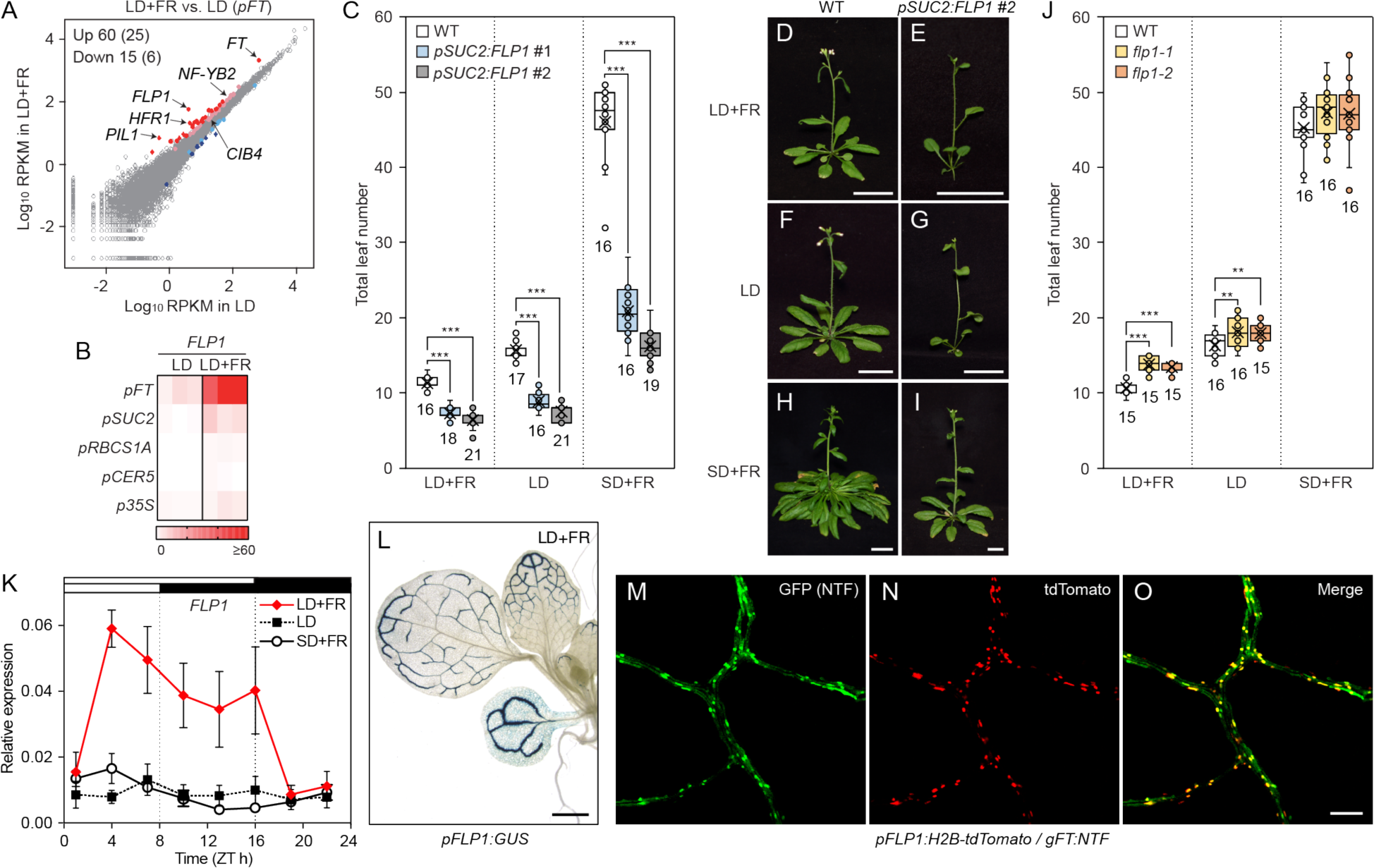
FLP1 is a photoperiodic flowering activator expressed in the FT-producing cells. (A) Comparison between translatome datasets of the *pFT:FLAG-GFP-RPL18* plants grown under LD and LD+FR conditions. X and Y axes are log_10_-transformed RPKM values in LD and LD+FR, respectively. Red/pink and blue/light blue color dots indicate significantly up-regulated and down-regulated genes in LD+FR (FDR < 0.05). Red and blue colors exhibit more than 2- fold difference, while pink and light blue are less than 2-fold difference. The number of the genes with FDR < 0.05 and, within the parenthesis, that of genes with FDR < 0.05 and more than 2-fold difference, are shown in the left corner. The positions of *FT*, *FLP1*, FR-marker genes (*PIL1* and *HFR*), and *FT* regulators (*CIB4* and *NY-FB2*) are indicated. (B) Tissue/cell-specific translational levels of *FLP1* mRNA in TRAP-seq. Letters on the left indicate the tissue/cell-specific TRAP lines used. The growth conditions are displayed on the top. The colors of squares indicate RPKM values in each biological replicate (*n* = 3). (C) Accelerated flowering of the *pSUC2:FLP1* overexpressors. The bottom and top lines of the box indicate the first and third quantiles, and the bottom and top lines of the whisker denote minimum and maximum values. Circles indicate inner and outlier points. The bar and the X mark inside the box are median and mean values, respectively. The numbers below the box indicate sample sizes. Asterisks denote significant differences from WT (\*\*\**P*<0.001, *t*-test). (D–I) The representative images of flowered WT (D, F, and H) and *pSUC2:FLP1* #2 (E, G, and I) plants grown under the LD+FR (D and E), LD (F and G), and SD+FR (H and I) conditions. Scale bar, 2 cm. (J) The flowering phenotype of the *flp1* mutants. The results were obtained and displayed the same way as the *pSUC2:FLP1* plant results shown in (C). Asterisks denote significant differences from WT (\*\**P*<0.01; \*\*\**P*<0.001, *t*-test). (K) Time course of *FLP1* gene expression in WT plants grown under LD+FR, LD, and SD+FR. The results represent the means ± SEM. (*n* = 3 biologically independent samples). White and black bars on top indicate time periods with light and dark. (L) *FLP1* promoter-driven GUS activity under the LD+FR conditions. There was no visible staining around the SAM and in the roots. Scale bar, 1 mm. (M–O) Overlapped distribution patterns of *FT-* and *FLP1*-promoter activities in leaf minor veins in LD+FR. *pFT:NTF* (M), *FLP1:H2B-tdTomato* (N), and merge of GFP and RFP channel images (O). Note that GFP signals from NTF were observed in nuclei but also sometimes in both nuclei and the cytosol, while RFP signals from H2B-tdTomato exclusively existed in the nuclei. Scale bar, 50 μm.

Instead, we found that the translated mRNA level of a gene named *FLP1* (At4g31380), a homolog of the meristem-expressing flowering activator *FPF1,*^15,16^ was upregulated predominantly in *FT*-producing cells under LD+FR conditions (Figures 3A, 3B and S5). *FLP1* encodes a 14-kDa protein without any known functional domains. Due to its substantial homology with FPF1 (63% identity and 77% similarity over the entire 124 amino acids), we hypothesized that *FLP1* may also play a role in regulating flowering. To investigate this hypothesis, given the phloem-specific expression of *FLP1*, we overexpressed *FLP1* using the *SUC2* promoter in wild-type (WT) plants possessing the *pFT:GUS* reporter (Figure S6).^21^ Remarkably, the *pSUC2:FLP1* lines displayed strong early flowering phenotypes under all growth conditions, including short-day conditions with sunlight R/FR (SD+FR), which usually do not induce *FT* expression (Figures 3C–I). We generated *flp1* mutants to assess the loss-of-function effect of *FLP1* (Figure S7). Consistent with the high expression of *FLP1* under LD+FR conditions (Figure 3B), the *flp1* mutants exhibited a late flowering phenotype in LD+FR (Figure 3J). These results indicate that FLP1 functions as a flowering activator.

Having established the genetic role of *FLP1* in flowering regulation, we then conducted a detailed analysis of *FLP1* expression patterns. By analyzing the diurnal time course of *FLP1* expression levels under LD+FR, LD, and SD+FR conditions, we found that *FLP1* was specifically induced under LD+FR conditions during the daytime (Figure 3K). This indicates that both the LD photoperiod and the appropriate R/FR ratio mimicking natural sunlight are required to induce the *FLP1* gene, similar to the morning induction of *FT*.

The *Arabidopsis* genome contains another uncharacterized *FLP1* homolog, *FLP2* (At5g10625). Unlike *FLP1*, our TRAP-seq datasets and qRT-PCR analysis demonstrated that neither *FLP2* nor *FPF1* showed induction by the R/FR light adjustment, although *FLP2* also exhibited the highest expression in *FT*-producing cells among tested tissue/cell types (Figure S8).

We next examined *FLP1* spatial expression patterns using the *FLP1* promoter-driven *GUS* (ß- Glucuronidase) gene to assess whether *FLP1* induction is tissue-specific (Figure 3L). The spatial expression pattern of *FLP1* in LD+FR resembled that of *FT*.^21^ To more precisely investigate the extent to which the spatial expression patterns of *FLP1* overlap with *FT*, we generated the *pFLP1:H2B-tdTomato/pFT:NTF* transgenic line (NTF: Nuclear Targeting Fusion protein, which contains GFP^36^). This dual reporter line demonstrated that *FLP1* spatial expression signals highly overlapped with *FT-*expressing companion cells in leaf minor veins (Figures 3M–O and S9).

Thus, we conclude that *FLP1* is spatiotemporally induced in a manner similar to morning *FT* under natural LD conditions to regulate flowering.

### *FLP1* promotes flowering in parallel with *FT*

Because *FLP1* is expressed in *FT*-producing cells and promotes flowering, we investigated the genetic relationship between *FLP1* and *FT*. To assess whether the early flowering phenotypes in *pSUC2:FLP1* lines were caused by an increase in *FT* expression, we measured *FT* expression levels in 2-week-old plants. *FT* expression was significantly higher in the *pSUC2:FLP1* lines under both LD+FR and LD conditions (Figure 4A). The *pFT:GUS* assay demonstrated that *FLP1* overexpression enhanced *FT* promoter activity specifically in cotyledons (Figures 4B–D).

**Figure 4.**
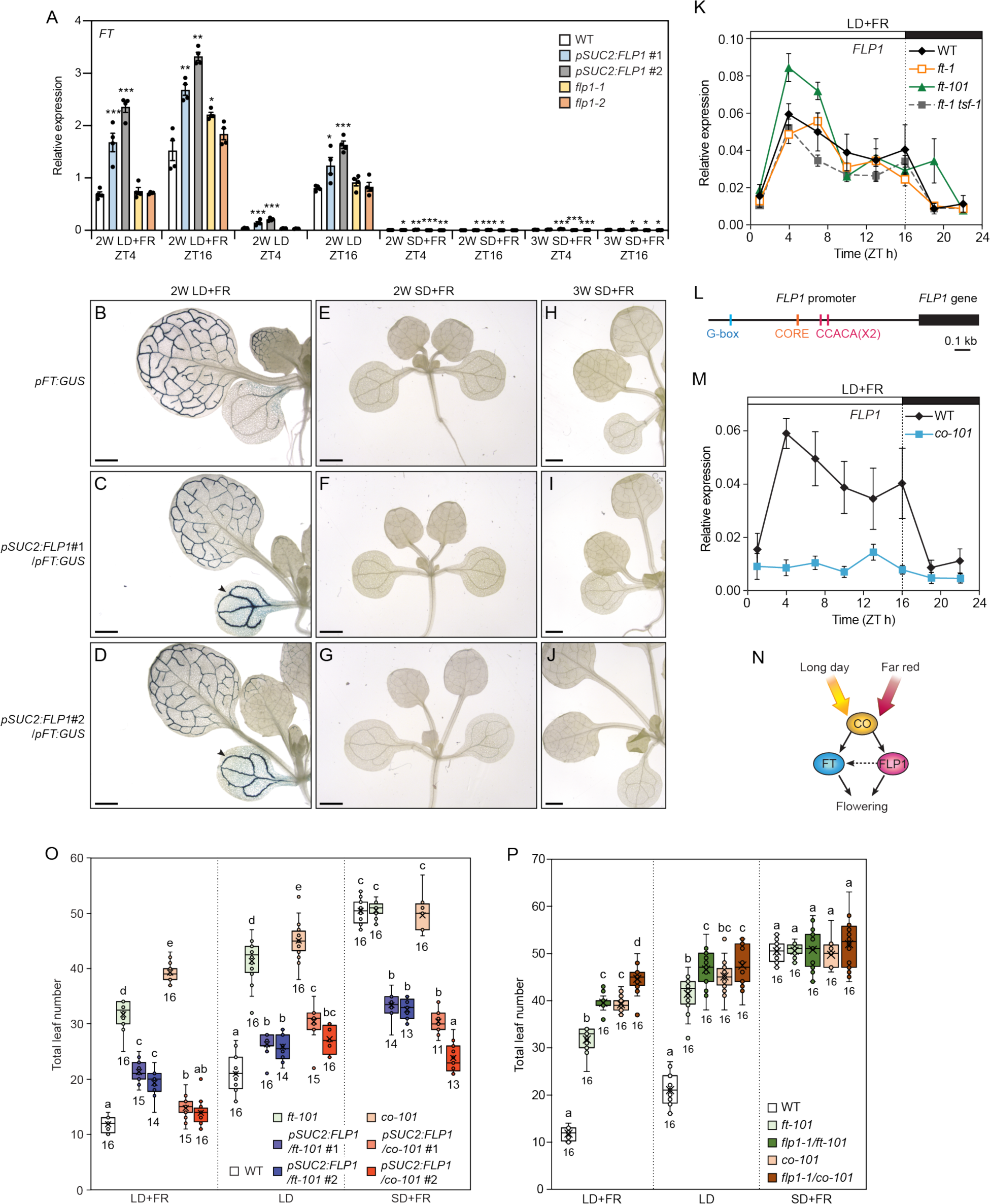
FLP1 induces flowering independently of FT in a CO-dependent manner. (A) *FT* expression in WT, *pSUC2:FLP1,* and *flp1* mutants. *FT* gene expression levels were analyzed in the plants grown under LD+FR, LD, and SD+FR for 2 or 3 weeks and harvested at ZT4 and ZT16. The results represent the means ± SEM. Each dot indicates a biological replicate (*n* = 4). Asterisks denote significant differences from WT (\**P*<0.05; \*\**P*<0.01; \*\*\**P*<0.001, *t*- test). (B–J) The effect of *FLP1* overexpression on the *FT* promoter activity. GUS staining assay was conducted using *pFT:GUS* lines with WT background (B, E, and H), *pSUC2:FLP1* #1 (C, F, and I), and *pSUC2:FLP1* #2 (D, G, and J) grown under LD+FR conditions for 2 weeks (B–D), SD+FR for 2 (E–G) and 3 weeks (H–J). Arrowheads indicate increased *FT* promoter activity repeatedly observed in cotyledons of the *pSUC2:FLP1* lines grown in LD+FR. Scale bars, 1 mm. (K) Time course of *FLP1* expression in WT, *ft-1*, *ft-101,* and *ft-1 tsf-1* double mutant plants grown in LD+FR. White and black bars on top indicate time with light and dark. The results represent the means ± SEM (*n* = 3 biologically independent samples). (L) The positions of CO-responsive element (CORE), CCACA motif, and G-box *cis*-element sequences on the 1.8 kb-upstream of *FLP1*, which was used as the *FLP1* promoter sequences in our experiments. (M) Time course of *FLP1* expression in WT and *co-101* mutant plants grown in LD+FR. The results represent the means ± SEM (*n* = 3 biologically independent samples). (N) A Model of the CO-dependent regulation of FLP1 and FT. CO regulates the expression of both *FLP1* and *FT* in photoperiod- and far-red light-dependent manners. FT and FLP1 promote floral transition in parallel. FLP1 may increase *FT* expression under LD potentially through the unknown positive feedback pathway. (O) The effect of *FLP1* overexpression on flowering of *ft-101* and *co-101* mutants. The bottom and top lines of the box indicate the first and third quantiles, and the bottom and top lines of the whisker denote minimum and maximum values. Circles indicate inner and outlier points. The bar and the X mark inside the box indicate median and mean values, respectively. The numbers below the box indicate the sample sizes. Different letters indicate statistically significant differences (*P*<0.05, Tukey’s test). (P) The effect of the *flp1* mutation on flowering time of the *ft-101* and *co-101* mutants. Different letters indicate a statistically significant difference (*P*<0.05, Tukey’s test).

However, under SD+FR conditions, *FT* expression levels remained low in the *pSUC2:FLP1* lines, although flowering timing of SD+FR-grown *pSUC2:FLP1* is comparable with LD-grown WT plants (Figures 3C, 4A, and 4E–G). This low *FT* level persisted even in 3-week-old plants in SD+FR (Figures 4A and 4H–J). In addition, despite their late flowering phenotype of the *flp1* mutants in LD+FR, the *flp1* mutants did not exhibit decreased *FT* expression under LD+FR conditions (Figure 4A). Hence, *FT* expression change alone cannot fully explain the flowering acceleration caused by *FLP1*. To explore other potential factors contributing to *FLP1*-induced flowering acceleration, we examined the expression of *TWIN SISTER OF FT* (*TSF*), the closest homolog of *FT* with a similar function in floral induction (Figure S10).^37^ However, we observed no significant difference in *TSF* expression among the plant lines examined, suggesting that neither changes in *FT* nor *TSF* expression can fully account for the flowering acceleration mediated by FLP1.

Next, we explored the possibility that *FLP1* accelerates flowering as a downstream component of *FT* by quantifying *FLP1* expression in *ft* and *ft tsf* double mutants. *FLP1* expression levels in these mutants were comparable with those in WT plants (Figure 4K), indicating that *FT* (and *TSF*) are not upstream of *FLP1*. Given the similar spatial and temporal expression patterns of *FLP1* and *FT* under natural light conditions, we began to suspect that core modulators of *FT* expression might drive their similar patterns of expression. CO, a key transcriptional activator of *FT*, binds to CO-responsive elements [COREs; TGTG(N2–3)ATG] and CCACA motifs located proximal to the transcriptional start site of *FT.*^22^ Finding CORE and CCACA motifs in the *FLP1* promoter (Figure 4L) prompted us to analyze *FLP1* expression in the *co* mutant. The induction of *FLP1* was nearly abolished in this mutant under LD+FR conditions (Figure 4M), suggesting that CO may directly regulate *FLP1*. These results indicate that FLP1 is a component within the photoperiod pathway and CO likely regulates the transcription of *FT* and *FLP1*, which promote flowering in parallel, in the same companion cells under natural light conditions (Figure 4N).

Since *FLP1* appears downstream of CO in parallel with *FT*, we transformed *pSUC2:FLP1* into *ft-101* and *co-101* mutant lines to ask whether *FLP1* alone was sufficient to induce flowering independently of *FT*. Multiple *pSUC2:FLP1/ft-101* and *pSUC2:FLP1/co-101* lines flowered early compared to the *ft-101* and *co-101* parental lines in LD+FR, LD, and SD+FR (Figures 4O and S11), indicating that functional *CO* and *FT* are not essential for *FLP1*-dependent flowering induction once *FLP1* is expressed. We further analyzed the flowering time of *flp1-1 ft-101* and *flp1-1 co-101* double mutants (Figure 4P). The *flp1-1 ft-101* mutant flowered later than the *ft-101* mutant under LD+FR and LD conditions, indicating that *FT* and *FLP1* act additively to promote flowering. The *flp1-1 co-101* lines also showed later flowering than the *co-101* line in LD+FR, although the difference was smaller (Figure 4P). Since CO function was necessary for *FLP1* induction, the small but significant delay in flowering in *flp1-1 co-101* double mutant lines was unexpected, suggesting that even the marginal levels of *FLP1* mRNA present in the *co-101* mutant might contribute to the induction of flowering or that *FLP1* may also regulate flowering other than the photoperiodic pathway.^15^ These findings indicate that FLP1 may promote flowering through an *FT*-independent mechanism.

### *FLP1* promotes inflorescence stem growth

We speculated on the reasons behind plants equipping FLP1 as an independent flowering inducer parallel to FT. One possibility is that FLP1 acts as a booster of flowering to ensure developmental transition occurs at the optimal timing under natural light conditions. However, beyond its role in flowering, we also observed that *FLP1* overexpressors and *flp1* mutants exhibited significantly elongated and shortened hypocotyl and leaf length, respectively (Figures 5A–C), similar to previous studies on *FPF1* overexpression phenotypes.^15^ These results suggest that FLP1 possesses developmental functions distinct from FT.

**Figure 5.**
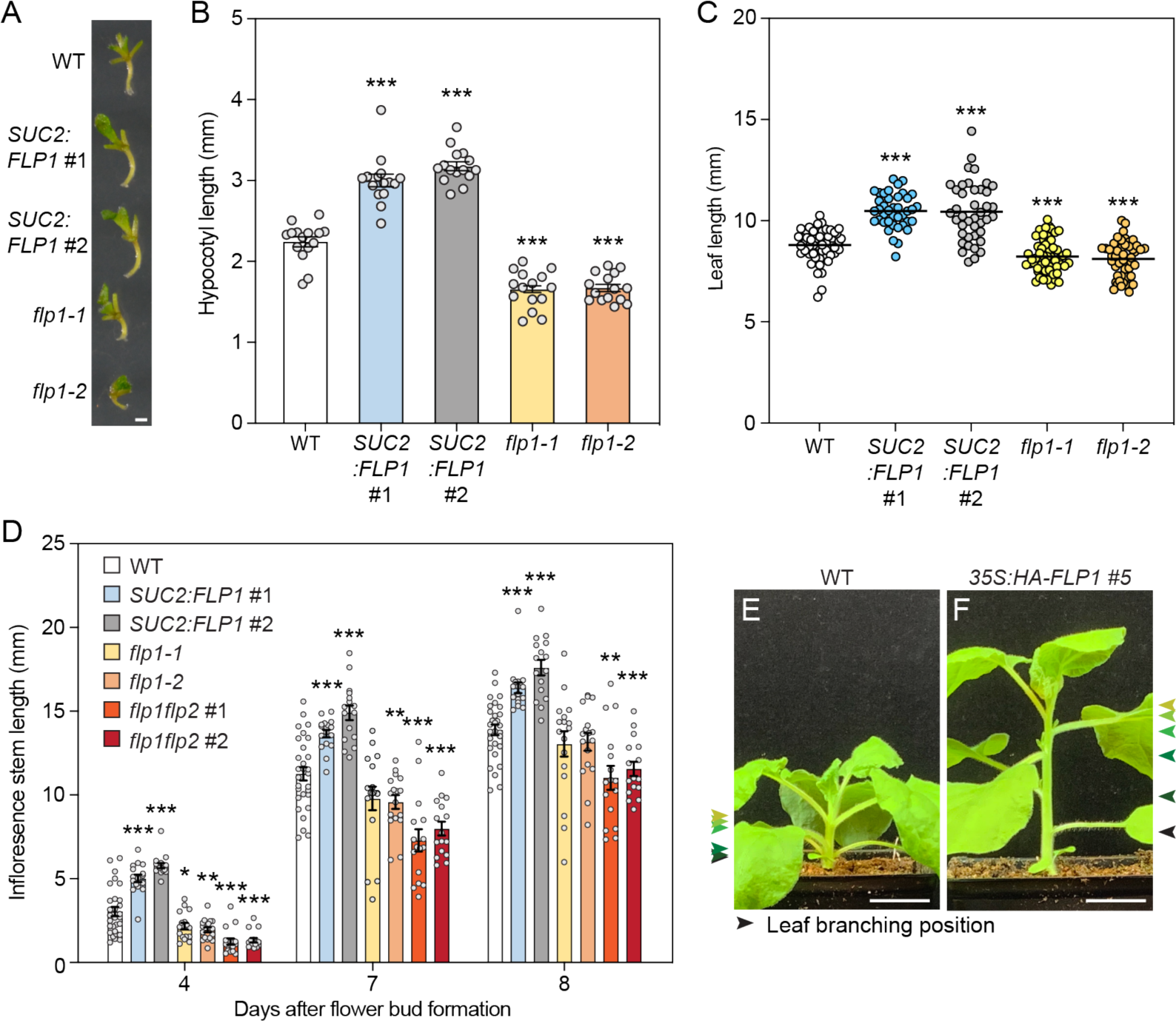
FLP1 promotes stem elongation. (A) Representative images of hypocotyls of 10-day-old WT, *pSUC2:FLP1* plants, and *flp1* mutants under LD+FR conditions. Scale bar, 1 mm. (B) Hypocotyl length of 10-day-old WT, *pSUC2:FLP1,* and *flp1* plants grown under LD+FR conditions. The results represent the means ± SEM of independent biological replicates. Each dot indicates a biological replicate (*n* = 15). (C) Length of 1^st^ and 2^nd^ true leaves (from the tip of the leaf blade to end of the petiole) of 14- day-old WT*, pSUC2:FLP1,* and *flp1* mutants grown under LD+FR conditions. Each dot indicates a biological replicate (*n* ≥ 40). Black bars indicate means. Asterisks denote significant differences from WT (\*\*\**P*<0.001, *t*-test). (D) The effects of overexpression of *FLP1* and mutations of *FLP1* and *FLP2* on inflorescence stem elongation. Stem length was measured 4, 7, and 8 days after the visible flower bud formation at the SAM. The results represent the means of independent biological replicates ± SEM. Each dot indicates a biological replicate (*n* = 16–29). Asterisks denote significant differences from WT (\**P*<0.05; \*\**P*<0.01; \*\*\**P*<0.001, *t*-test). (E and F) The effect of overexpression of *FLP1* on stem elongation of *Nicotiana benthamiana*. The arrowheads indicate the position of the leaf branching. The same colors of the arrowheads in (E) and (F) exhibit the same developmental sets of leaf branches in these plants. Plants shown in the figures are LD-grown 4-week-old plants. Scale bar, 2 cm.

Recent studies revealed that FPF1 homologs in rice (ACE1) and tomato promote stem elongation.^17,18^ These reports led us to hypothesize that FLP1 may also affect inflorescence stem growth. We analyzed the length of inflorescence stems over time following the floral transition. In this experiment, we individually set the day 0 of the stem measurement for each plant as the day when the first flower bud structure of the plant was visible at the SAM (before the recognizable stem elongation). Thus, flowering timing differences among individuals do not affect the results of timings of inflorescence stem elongation. Our stem measurement analysis revealed that *FLP1* overexpression significantly promoted inflorescence stem elongation (Figure 5D). The negative effect of the *flp1* mutations on stem elongation was moderate, suggesting the involvement of other redundant factors. Considering that *FLP2* was also expressed in *FT*- producing cells, albeit at lower levels, and that FLP1 and FLP2 share 65% amino acid identity and 78% amino acid similarity, we hypothesized that FLP1 and FLP2 may have overlapped functions. To investigate this, we generated the *flp1 flp2* double mutants by genome editing (Figure S12). The *flp1 flp2* mutants showed more severe stem elongation defects than the *flp1* single mutants (Figure 5D), suggesting functional redundancy existed between *FLP1* and *FLP2*. Furthermore, we demonstrated that *Arabidopsis FLP1* overexpression induced substantial stem elongation in *N. benthaminana* (Figures 5E, F, and S13). As *FPF1*/*FLP1* homologs exist in a wide variety of angiosperms (monocots and eudicots),^17^ their functional roles in stem elongation are likely conserved among diverse plant species. In addition, overexpression of *N. tabacum FPF1* in a few *Nicotiana* varieties induced early flowering with an elongated bolted stem.^38^ Taken together, our findings demonstrate that FLP1 regulates both flowering and stem elongation processes in *Arabidopsis*.

### *FLP1* induces a floral homeotic gene expression

To identify genes associated with the flowering and growth phenotypes of *FLP1* overexpressors and *flp1* mutants, we performed RNA-seq analysis by comparing 2-week-old WT with *pSUC2:FLP1* #2 under LD+FR and SD+FR conditions as well as *flp1-1* under LD+FR conditions sampled at ZT4 (Data S4–6). *pSUC2:FLP1* #2 showed upregulation of auxin- and gibberellin-responsive genes under LD+FR and SD+FR conditions, respectively (Figure S14), implying possible contributions of these hormones in flowering and growth phenotypes of *pSUC2:FLP1* lines. Previous studies also showed that overexpression of *FPF1* homologs in rice, *ROOT ARCHITECTURE ASSOCIATED 1* and *FPF1-LIKE 4*, enhanced auxin levels and signaling.^39,40^ Moreover, gibberellin is crucial for *FPF1*-dependent flowering and stem internode phenotypes.^15–17^ Therefore, we quantified the levels of auxin and gibberellin, together with other phytohormones also implicated in growth and cellular differentiation in *Arabidopsis* using WT, *pSUC2:FLP1 #2* and *flp1-1* lines (Figure S15). Plants significantly changed levels of many hormones between LD+FR and SD+FR conditions (Figure S15). However, *FLP1* gene expression levels did not affect indole-3-acetic acid (IAA) contents at least at the whole plant levels. Gibberellin A_4_ (GA_4_) was decreased in the *flp1-1* line, but no change in *pSUC2:FLP1* #2, suggesting that GA4 accumulation is not the primary cause of flowering acceleration and stem elongation. We also observed that there were no obvious changes in the levels of other phytohormones by the alternation of *FLP1* expression levels, suggesting that the developmental changes observed in *FLP1* overexpressors and *flp1* mutants are not primarily attributable to the alterations in phytohormone levels.

By analyzing the overlap of upregulated genes in *pSUC2:FLP1* under LD+FR and SD+FR conditions and downregulated genes in *flp1-1* under LD+FR conditions, we identified 14 genes as putatively acting downstream of FLP1 under these conditions (Figure 6A). Most of these genes encode nutrient transporters (that could be important for growth and development); however, we also found two TF genes, *SEP3* and *PHYTOCHROME INTERACTING FACTOR 8* (*PIF8*), among the 14 genes. PIF8 promotes hypocotyl elongation under FR light conditions,^41^ and could plausibly contribute to the hypocotyl phenotypes of *FLP1* overexpressors and *flp1* mutants. *SEP3* is a key floral homeotic E-class gene whose overexpression causes early flowering.^14^

**Figure 6.**
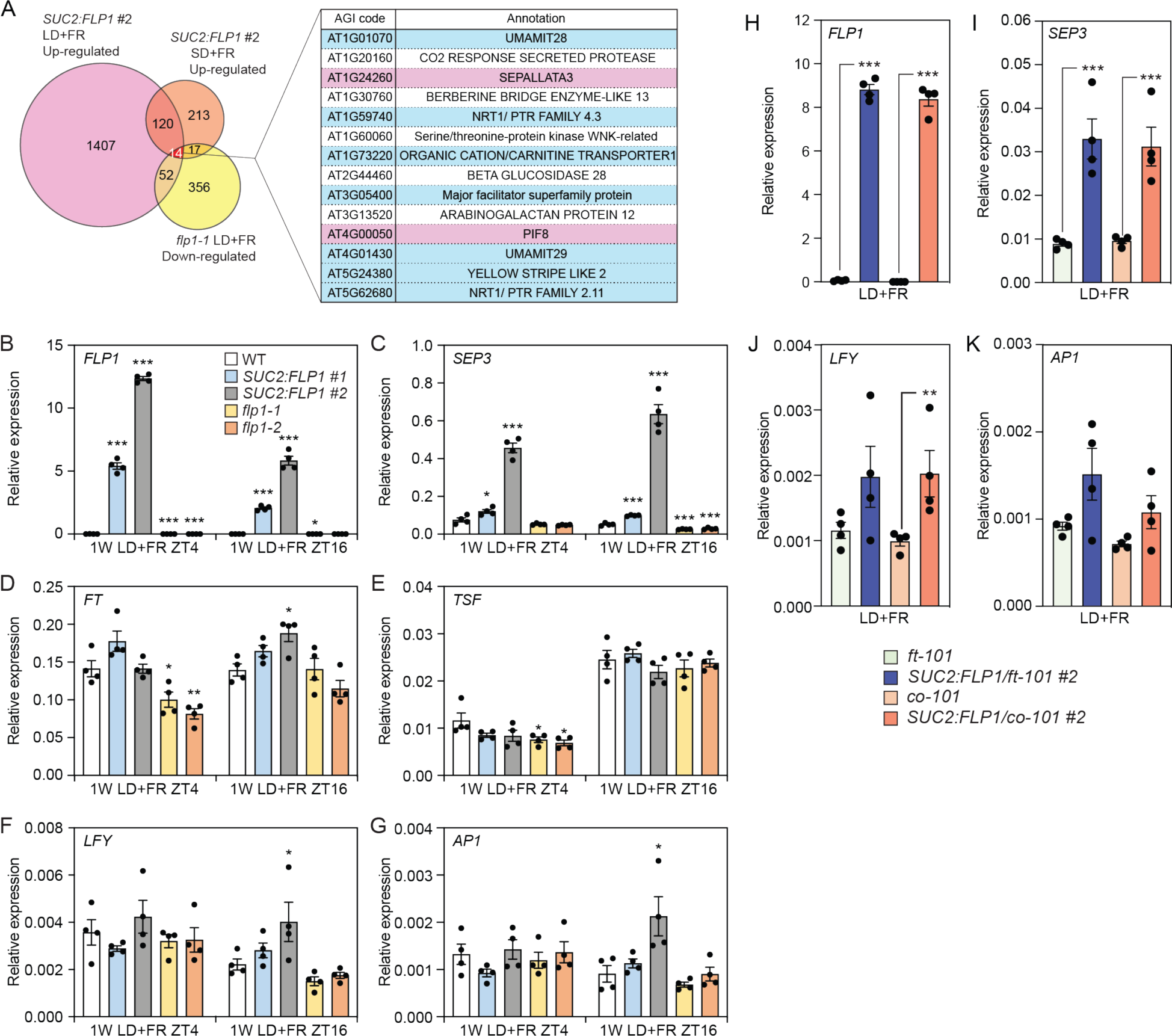
FLP1 induces the floral homeotic gene SEP3 in a FT-independent manner. (A) A Venn diagram consisting of up-regulated genes in 2-week-old *pSUC2:FLP1* #2 plants grown under LD+FR and SD+FR, and down-regulated genes in *flp1-1* mutant grown under LD+FR compared to the WT grown under the same conditions. The table on the right lists 14 genes that were overlapped by three conditions. Genes encoding transcription factors and nutrient transporters were highlighted in pink and blue, respectively. (B–G) The effect of *FLP1* levels on the gene expression at 1-week-old plants. Plants were grown under LD+FR conditions for 1 week and harvested at ZT4 and ZT16. The relative expression levels of *FLP1* (B), *FT* (D), *TSF* (E), and floral homeotic genes, *SEP3* (C), *LFY* (F), and *AP1* (G) were analyzed. The results represent the means ± SEM. Each dot indicates a biological replicate (*n* = 4). Asterisks denote significant differences from WT (\**P*<0.05; \*\*\**P*<0.001, *t*-test). (H–K) The effect of *FLP1* overexpression on the gene expression in the *ft-101* and *co-101* mutant backgrounds. The expression of *FLP1* (H) and floral homeotic genes, *SEP3* (I), *LFY* (J), and *AP1* (K) were analyzed. Plants were grown under LD+FR conditions for 2 weeks and harvested at ZT4. The results represent the means ± SEM. Each dot indicates a biological replicate (*n* = 4). Asterisks denote significant differences from the parental line (\**P*<0.05; \*\**P*<0.01; \*\*\**P*<0.001, *t*-test).

Consistent with our RNA-seq analysis, qRT-PCR analysis demonstrated that both *SEP3* and *PIF8* were upregulated in *pSUC2:FLP1* and downregulated in *flp1* mutants under LD+FR conditions (Figures S16A and C). However, *FT* was also upregulated in 2-week-old *pSUC2:FLP1* lines under LD+FR conditions (Figure 4A), which could potentially contribute to the observed increase in *SEP3* levels. To eliminate the possible influence of *FT* on *SEP3* induction, since *FT* expression increases with plant maturation,^42^ we examined gene expression in younger seedlings and tested whether *SEP3* induction started before or after *FT* induction by FLP1. In 1-week-old plants, *SEP3* was significantly upregulated in *pSUC2:FLP1* lines at both ZT4 and ZT16 according to *FLP1* expression levels, while neither *FT* nor *TSF* showed consistent upregulation by FLP1 (Figures 6B–E), indicating that FLP1 induced *SEP3* before *FT* induction.

Similarly, under SD+FR conditions, although the amplitude was lower in *pSUC2:FLP1* #1, both *pSUC2:FLP1* lines showed significant *SEP3* upregulation at ZT4 in 2-week-old and ZT16 in 3- week-old plants (Figure S16B). Since *FT* expression is extremely low under SD+FR conditions, these results support the notion that *FT* is not essential for FLP1-dependent *SEP3* induction. To genetically confirm this finding, we analyzed *SEP3* expression in *pSUC2:FLP1/ft-101* #2 and *pSUC2:FLP1/co-101* #2 lines. Compared with the parental lines, these transgenic lines showed significant upregulation of *SEP3* in both RNA-seq and qRT-PCR analyses (Figures 6H and I, Data S7). Using RNA-seq and qRT-PCR, we further addressed whether repressors of *SEP3,* such as *SVP*,^11^ might be suppressed by FLP1; however, we did not observe a clear correlation between the expression of *SEP3* and its repressors including *SVP* (Figure S17).

In the current model of the FT-dependent flowering pathway, FT forms a complex with the bZIP transcription factor FD that induces the expression of *LFY* and *AP1*, followed by the derepression and direct activation of *SEP3.*^8,9,11^ Because FLP1 does not rely on FT for its *SEP3* induction, we asked next whether FLP1 activates *SEP3* through the pathway mediating *LFY* and *AP1*. We therefore examined the expression of these FT downstream genes in our lines. At 1 week old, the *pSUC2:FLP1* lines did not consistently upregulate *LFY* and *AP1* (Figures 6F and G), in marked contrast to *SEP3* upregulation. Furthermore, in *pSUC2:FLP1* plants grown under SD+FR conditions for 2–3 weeks, the induction of *LFY* and *AP1* was weaker than that of *SEP3* (Figures S16F and H). In the 2-week-old *ft-101* and *co-101* backgrounds, significant upregulation of *AP1* expression by *pSUC2:FLP1* was not observed, while *SEP3* was clearly upregulated (Figures 6H–K). Taken together, our results suggest that FLP1 induces flowering through *SEP3* induction in an *LFY*- or *AP1*-independent manner, which differs from the FT-dependent mechanism that requires *LFY* and *AP1* induction prior to *SEP3*.

### FLP1 acts at the shoot apical meristem potentially through the interaction with TFs

To test whether FLP1 is active at the SAM in promoting flowering, we expressed *FLP1* using the SAM-specific *UNUSUAL FLORAL ORGANS* (*UFO*) promoter.^43^ Our *pUFO:FLP1* lines showed significant early flowering under LD and SD+FR conditions (Figure 7A), although it did not flower as early as *pSUC2:FLP1* lines likely due to the less FLP1 protein amount expressed in these lines. This result indicates that FLP1 can promote flowering at the SAM. Because FLP1 is a small protein lacking known functional domains, we questioned if FLP1 has TF interacting partners to modulate plant gene expression, similar to the FT–FD interaction. Employing a yeast two-hybrid *Arabidopsis* TF library comprising 1,736 TFs, we identified potential interactors with FLP1 (Data S8). To further narrow down functionally relevant interactors, we included FPF1 as a bait protein, considering the functional similarity between FLP1 and FPF1. We found that 162 TFs interacted with both FLP1 and FPF1 in yeast (Data S8). Interestingly, neither FLP1 nor FPF1 interacted with FD in our yeast two-hybrid assay, suggesting that the mode of action of FLP1 on flowering could be different from that of FT in *Arabidopsis*. Of these candidates, we selected 22 TFs based on their known functions in floral transitions and inflorescence stem elongation and validated their physical interaction with FLP1 using an *in vitro* protein-protein binding assay (Data S8 and Figures S18A–C).^44,45^ In this assay, 16 TFs exhibited more than 2- fold signals compared to the negative control (Figure S18A). Notably, these positive TF interactors included those expressed at the SAM and involved in regulating cell fate determination during the floral transition, such as WUSCHEL (WUS), DORNRÖSCHEN (DRN), DORNRÖSCHEN-LIKE (DNL), and PUCHI.^8,46^

**Figure 7.**
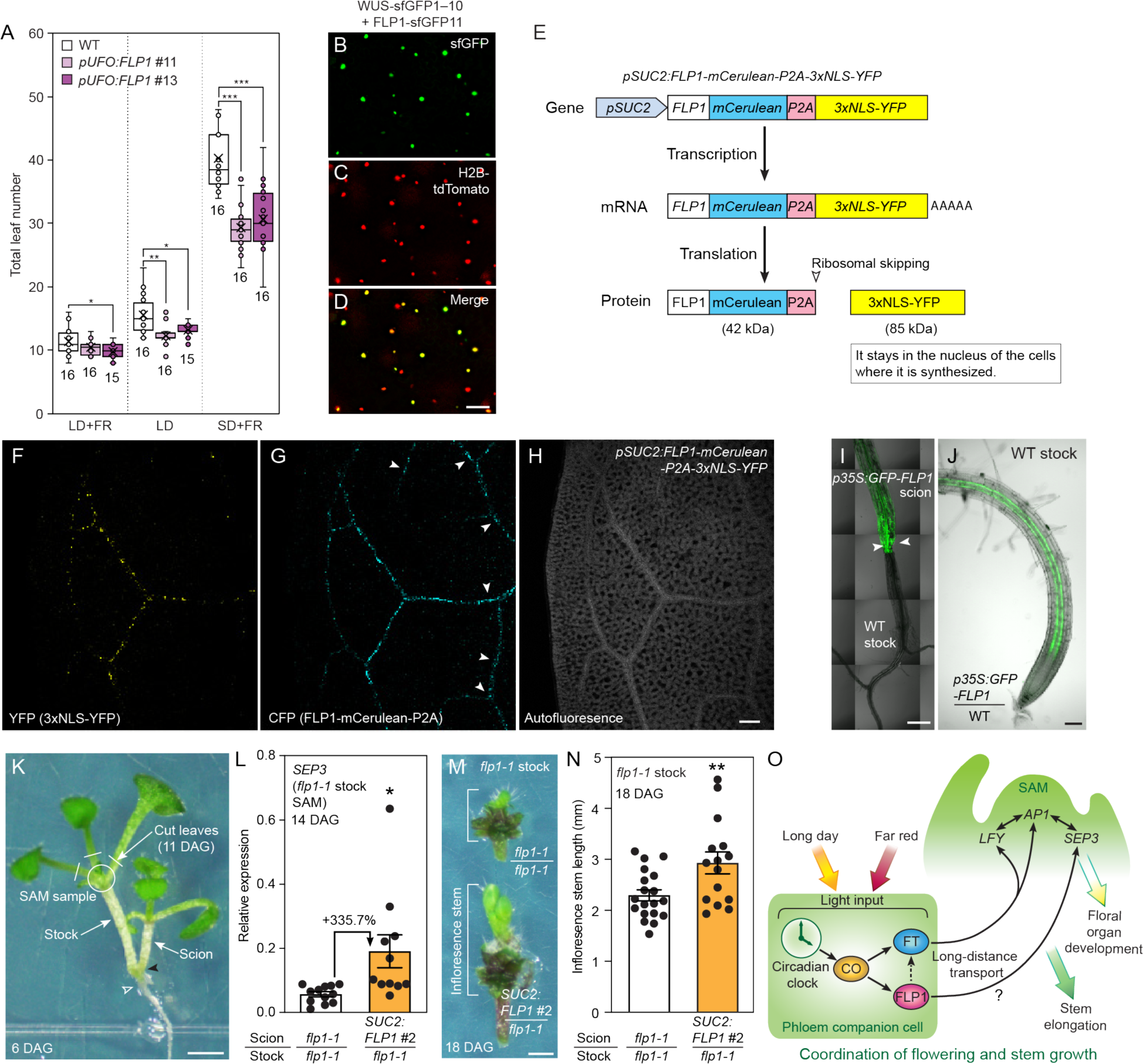
FLP1 may be a vascular-mobile protein that may function in the tissues distantly from leaves. (A) The flowering phenotype of the *pUFO:FLP1* lines grown under LD+FR, LD, and SD+FR. Asterisks denote significant differences from WT (\**P*<0.05; \*\**P*<0.01; \*\*\**P*<0.001, *t*-test). (B–D) BiFC assay for physical interaction between FLP1-sfGFP11 and WUS-sfGFP1–10 in tobacco epidermal cells shown in the reconstitution of GFP signals (B). Histone H2B-tdTomato (C) marks the positions of nuclei. The image (D) is the merged image of GFP and RFP channels. Scale bar, 100 µm. (E) A schematic diagram of the P2A system that enables to independently express two proteins, FLP1-mCerulean and 3xNLS-YFP proteins, using the single promoter-driven construct. (F–H) Confocal images of cleared true leaf of *pSUC2:FLP1-mCerulean-P2A-3xNLS-YFP* excited with 405 nm. The locations of 3xNLS-YFP (F) and FLP1-mCerulean-P2A (G) in a true leaf are shown. The image (H) shows the autofluorescence of the same sample. Arrowheads in (G) indicate mCerulean signals in the vasculatures where YFP signals in (F) are low. Scale bar, 100 µm. (I) Grafting junction (indicated by arrowheads) of *p35S:GFP-FLP1* scion and WT rootstock. The images of the GFP and DIC channels were merged. Scale bar, 500 µm. (J) Movement of the GFP-FLP1 protein to the WT rootstock from the *p35S:GFP-FLP1* scion. Scale bar, 100 µm. (K) The Y-shape grafted plant at 6 days after grafting (DAG). The circle indicates the location of the SAM of the stock used for gene expression analysis in (L). A black arrowhead and a white open arrowhead depict the positions of the graft junction and the bottom of the hypocotyl of the stock plant, respectively. Scale bar, 1 mm. (L) The effect of FLP1 derived from the scion on *SEP3* gene expression in the SAM of the *flp1-1* stock at 14 DAG. The results represent the means of independent biological replicates ± SEM. Each dot indicates a biological replicate derived from independent grafted plants (*n* = 11–12). The percentage signifies a relative difference compared to the control *flp1-1* scion. An asterisk denotes a significant difference (\**P*<0.05, *t*-test). (M) Representative images of the SAM containing tissues of the *flp1-1* stock grafted with either *pSUC2:FLP1* #2 scion or the *flp1-1* scion on 18 DAG. The image was used to measure the height of the inflorescence stem. Scale bar, 1 mm. (N) The length of the inflorescence stem of the *flp1-1* stock at 18 DAG. The results represent the means of independent biological replicates ± SEM. Each dot indicates a biological replicate (*n* = 15–19). Asterisks denote significant differences (\*\**P*<0.01, *t*-test). (O) A model of coordination of flowering and inflorescence stem growth by FLP1 and FT. CO regulates the expression of both *FLP1* and *FT* in photoperiod- and far-red light-dependent manners. FT is synthesized in specific leaf phloem companion cells in response to environmental stimuli and moved to the SAM to induce flowering by directly upregulating *LFY* and *AP1*. FLP1 is also synthesized in the same cells where FT is synthesized. It can move through the phloem, and possibly the movement of FLP1 may be important for promoting floral transition through *SEP3* induction. FLP1 also participates in the inflorescence stem elongation.

A recent study revealed that WUS represses *SEP3* expression by direct binding to its promoter,^47^ implying a possible link between FLP1 and WUS proteins with *SEP3* transcription. It prompted us to conduct a BiFC assay to further confirm the interactions between FLP1 and WUS (Figures 7B–D and S18D–P). In the tobacco transient expression system, consistent with the results described above, FLP1 strongly interacted with WUS in the nucleus forming speckle-like structures (Figure S18P). Although further investigation is required to understand the molecular functions of FLP1, our results raised the possibility that FLP1 promotes flowering through the interaction with developmentally important TFs such as WUS, which directly alters *SEP3* gene expression. Given the TF-deactivating function of *Brachypodium* FPF1 homologs,^19^ FLP1 may attenuate WUS’s transcriptional activities through physical interaction.

### FLP1 may behave like a mobile flowering-promoting signal

FLP1 is a 14-kDa small protein synthesized in the same florigen-producing phloem companion cells active in long-distance transport, including the transportation of FT protein (Figures 2C and G). Previous proteome studies have detected FLP1 protein in various plant tissues distantly from leaves, including floral organs and roots.^48–50^ Similarly, the rice FLP1 homolog, ACE1, is broadly induced within the internodes of elongated stems upon submergence, but it is implicated to function at the intercalary meristem.^17^ Taken together with its flowering-promoting activity at the SAM (Figure 7A), it is plausible that FLP1 may act as a mobile signal for flowering. To explore this possibility, we investigated FLP1’s movement in leaves using the *pSUC2:FLP1- mCelurean-P2A-3xNLS-YFP* line (Figures 7E–H and S19). Since the P2A sequence causes protein cleavage during translation, two different proteins, FLP1-mCerulean and 3xNLS-YFP, are produced simultaneously by the same promoter activity (Figure 7E). In the tobacco transient assay, FLP1-mCelurean did not exhibit a specific cellular localization, whereas 3xNLS-YFP strongly accumulated in the nucleus, confirming the P2A protein cleavage system is working *in planta* (Figure S19A). In the *pSUC2:FLP1-mCelurean-P2A-3xNLS-YFP* line, we could track the movement of FLP1-mCerulean in leaf vascular tissues by comparing the localization patterns of CFP with nuclear-localized three tandemerized YFP likely to stay in the cells where synthesized. This *Arabidopsis* transgenic line showed early flowering (Figure S19B), indicating FLP1- mCerulean retained the flowering-promoting function. Even in the locations with relatively low *SUC2* promoter activity, the FLP1-mCerulean signal appeared more uniform within the phloem (Figures 7F and G), suggesting that FLP1-mCerulean is translocated from the originally expressing companion cells. To further confirm FLP1’s movement, we conducted grafting of *p35S:GFP-FLP1* shoot scions and WT rootstocks (Figures 7I and S20). The GFP-FLP1 protein exhibited no specific cellular localization as FLP1-mCerulean (Figures S20A and B). The GFP- FLP1 protein synthesized in the scion was observed along the phloem of the *Arabidopsis* WT rootstock near the tip (Figure 7J), indicating that GFP-FLP1 was translocated to the root through the vasculature.

Given its potential as a mobile signaling protein, we further studied whether the effect of FLP1 on flowering is transferable to the SAM using Y-shape micrografting (Figure 7K). Here, the hypocotyls of *flp1-1* or *pSUC2:FLP1* #2 scions were inserted into the hypocotyl of *flp1-1* stocks. Once the grafting was successful, on 11 days after grafting (DAG), all the leaves in the stock plants were carefully removed to ensure that only the scion plant controlled phloem streaming. In other words, the removal of the leaves relegated the *flp1-1* stock plant to acting primarily as a sink tissue. We assessed the movement of FLP1 from the scions to the SAM of the *flp1-1* stock based on the *SEP3* gene induction and flowering promotion of the stock. At 14 DAG, the *pSUC2:FLP1* #2 scion significantly increased *SEP3* expression in the SAM of the *flp1-1* stock compared to the *flp1-1* stock grafted with the *flp1-1* scion (Figure 7L). Consistent with the higher expression levels of *SEP3* at the SAM, the *pSUC2:FLP1* scion increased the inflorescence length of the *flp1-1* stock compared to the *flp1-1* stock with the *flp1-1* scion at 18 DAG (Figures 7M and N), indicating the progression of the flowering stage and potentially inflorescence stem elongation induced by FLP1 supplied from the scion. Although our Y-grafting experiments do not eliminate the movement of possible FLP1-induced molecules, the cumulative evidence supports the possibility that FLP1 might be a mobile protein regulating flowering time and stem elongation.

## Discussion

*FT* gene expression is confined to the specific phloem companion cells. Therefore, analyzing specific gene expression profiles in these specialized cells is crucial to understand how plants integrate environmental information and find the timing of the transition to the reproductive growth stage. We employed TRAP-seq translatome analysis that enabled us to isolate mRNAs specifically translated in *FT*-producing cells as well as in other tissue types (Figure 1). An advantage of translatome analysis is a higher correlation with proteome than transcriptome in eukaryote cells.^24^ Our TRAP-seq analysis revealed that uniquely translated mRNA populations in *FT*-producing cells resembled those in *SUC2*-expressing phloem companion cells, compared with those in photosynthetic *RBCS1A*-expressing tissues or in *CER5*-expressing epidermal tissues (Figures 1D–I and S2), indicating we succeeded isolating specialized companion cells expressing *FT*. Highly enriched translating mRNA in *FT*-expressing cells suggest that these cells might be differentiating or newly differentiated phloem companion cells, which are active in metabolism and transportation (Figures 2A–F). These cells uniquely express genes important for FT transport, together with *FT* (Figure 2G), suggesting that FT protein synthesis and transport are coordinated. Our datasets also provided insights into tissue-specific expression of known *FT* transcriptional regulators (Figures S3 and S4). Notably, we observed more tissue-specific translational differences of negative *FT* regulators than of positive regulators. As often regulation of the expression of negative factors is the primary regulatory strategy of plants, *FT* tissue-specific expression may also be controlled by the presence or absence of specific negative regulators in different tissues.

Our TRAP-seq analysis also facilitated the identification of a novel flowering regulator. Using plants grown under modified LD growth conditions that mimic natural light quality, we have identified a previously uncharacterized FLP1 as a possible systemic protein involved in the regulation of both flowering and stem elongation (Figure 7O). FLP1 and FT induce flowering in parallel, but they are under the regulation of CO, suggesting that CO regulates multiple flowering signals simultaneously. Both under the CO regulation, FLP1 and FT form the coherent type 1 feed-forward loop (C1-FFL) with OR gate in natural long days to induce flowering (Figure 4N). The C1-FFL is the most common feed-forward loops that exist in biological systems, and the OR logic C1-FFL ensures continuous output generation (i.e., continuous flowering induction) even having an abrupt loss of input signals (i.e., variable nature of flowering inducing environmental conditions).^51,52^ Having the FT/FLP1 C1-FFL may facilitate plants to induce and maintain flowering status under ever-changing natural environments in spring.

Because *FLP1* is induced in LD+FR but not in LD, it is plausible that factors other than CO are required for *FLP1* gene expression. The promoter of the *FLP1* gene possesses a G-box element, a consensus DNA-binding site of PIF transcription factors (Figure 4L). Our recent study revealed that PIF7 contributes to the morning *FT* gene expression in a phyA-dependent manner under the same LD+FR conditions.^4^ It is interesting to ask whether plants use the same pathway to induce *FLP1* gene expression under natural light conditions.

In addition to its flowering role, FLP1 also possesses a distinct function from FT, which is the promotion of inflorescence stem elongation. The coordination between flowering and inflorescence stem elongation in plants remains poorly understood despite being a common phenomenon.^53^ A system using photoperiod- and R/FR ratio-dependent CO function to regulate the expression of both FLP1 and FT, each with both similar and unique functions, appeals as a convincing mechanism for the orchestration of the complex developmental changes that accompany the floral transition (Figure 7O). FLP1 and FT are synthesized in the same restricted leaf phloem companion cells, possibly similar to FT, FLP1 may be translocated to the SAM to accelerate the floral transition. Our gene expression analyses indicate that the primary target of FLP1 is *SEP3*, which probably contributes to flowering induction through positive feedback regulation among homeotic genes.^9,13^ In addition to the flowering promotion, unlike FT, FLP1 accelerated inflorescence stem elongation through an unknown mechanism. Given the presence of *FLP1* homologs across various plant species^17^ and the consistent role of a few characterized *FLP1* homologs as regulators of flowering, growth, or both,^15–18,38–40,54,55^ these factors may also function as systemic regulators to coordinate the growth and development in meristematic tissues in response to changing environments.

Although FPF1 family proteins, including FLP1, lack any known functional domains, a recent study unveiled that FPF1 homologs of *Brachypodium* physically repress the DNA binding of the FD1,^19^ raised a possibility that other FPF1 family proteins also deactivate TFs. In our Y2H analysis, neither FLP1 nor FPF1 interacted with FD (Data S8). Moreover, *Brachypodium FPF1* homologs genetically depend on functional *FT* to control flowering,^19^ while our study demonstrated that *FLP1* flowering function is independent from *FT* (Figure 4O). Although further confirmation is still required, these results suggest that *Arabidopsis* FD may not be involved in FLP1- or FPF1-dependent flowering regulation, unlike the case of *Brachypodium*.

We found that FLP1 strongly interacts with WUS. Since WUS is a direct negative regulator of *SEP3*, FLP1 might interfere with the transcriptional activity of WUS by attenuating its DNA binding activity. The *WUS* gene is essential for stem cell maintenance. Thus, its mutation causes pleiotropic defects, including the severe disruption of flower formation, preventing us from investigating the genetic interaction between *FLP1* and *WUS* using flowering time measurements. In the future, the FLP1’s role in *SEP3* induction and its possible interaction with WUS should be investigated more in-depth.

## Limitation of the study

Although our experimental evidence supports the hypothesis that FLP1 may act as a mobile signal of flowering, we have not yet directly confirmed its movement to the SAM. We have attempted to generate a variety of different FLP1 fusion constructs; however, based on the flowering and stem elongation phenotypes of the overexpressors, some fusion proteins appeared to lose their functionality. FLP1 apparently easily loses its activity by the protein fusion possibly due to the small molecular size (14 kDa). In addition, observation of the movement of small proteins (attached to larger tagged proteins) to the SAM in *Arabidopsis* often seems challenging due to the narrow path to the SAM as shown in the case of FT.^56^ Even after the FT protein had been recognized as a florigen for a while, its movement to the SAM in *Arabidopsis* had not been experimentally proven until recently.^57^ As the mobility of FT to the SAM was demonstrated through interaction with FD using the BiFC system incorporating superfold fluorescence proteins,^57^ the utilization of interacting TFs might help detect the movement of FLP1 to the SAM in the future. In this study, we demonstrated that FLP1 is active in the SAM to induce flowering (Figure 7A), suggesting that FLP1 proteins produced in the leaf phloem companion cells can be active if they move to the SAM. The conventional *Arabidopsis* micrografting experiment showed that GFP-FLP1 can be transferred through the phloem to the root tip (Figures 7I and J). The Y-shape micrografting experiments demonstrated that the flowering-promoting activity of FLP1 can be transmitted to the grafted *flp1-1* stock plants (Figures 7K–N). At this point, our data still cannot rule out the possibility that FLP1 promotes the production of other mobile signals of flowering, such as other signal proteins, phytohormones, and metabolites. However, if FLP1 mediates flowering induction via other molecules, it is unlikely to be FT since FLP1 induces flowering in an FT-independent manner (Figures 4O and S11). TSF is also unlikely, given the minor effect of FLP1 on *TSF* gene expression (Figure S10). Gibberellin or auxin appears not primary cause either, as FLP1 overexpression did not affect levels of these phytohormones (Figure S15). We have not measured levels of flowering-promoting metabolites such as trehalose-6-phosphate;^58^ however, our RNA-seq experiments suggest that the effect of FLP1 on genes related to sugar metabolism including *TREHALOSE-6-PHOSPHATE SYNTHASE 1* is minor (Figure S14). Although none of these known mobile signals of flowering might not account for the flowering-promoting activity of FLP1, we still need further investigation to examine whether FLP1 itself may act as a mobile signal of flowering and stem elongation. At least, our analysis indicates that *FT*-producing cells co-express *FLP1,* which could induce *SEP3* expression to contribute to floral transition and subsequent fluorescence stem elongation under natural long-day conditions.

## STAR Methods

### Key resource table

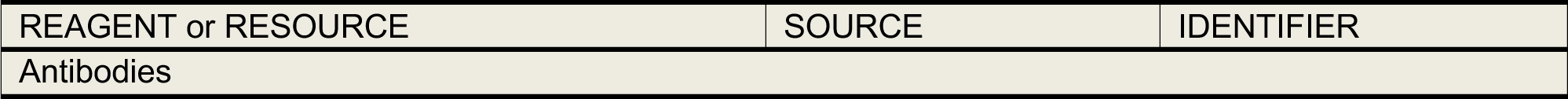
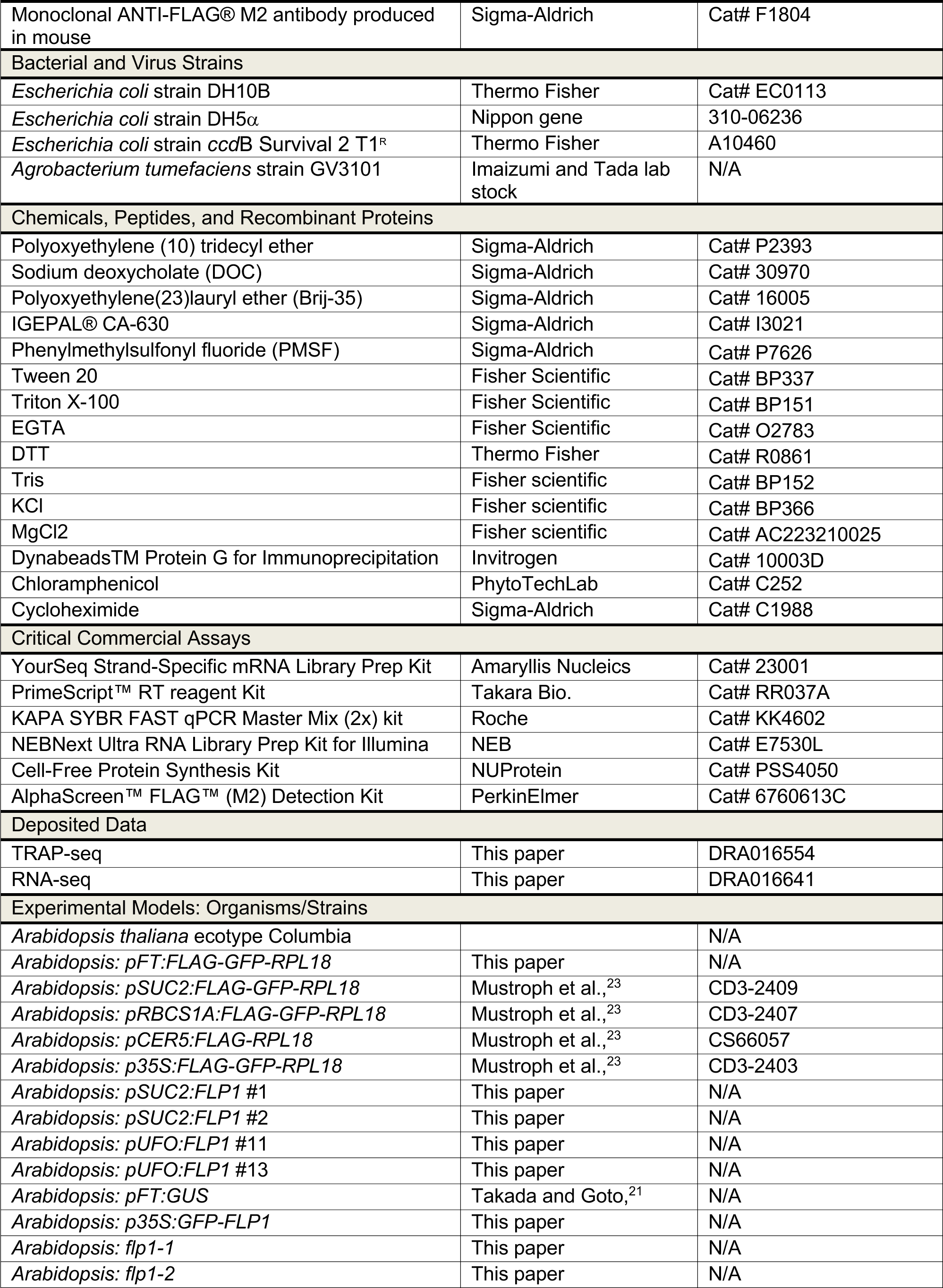
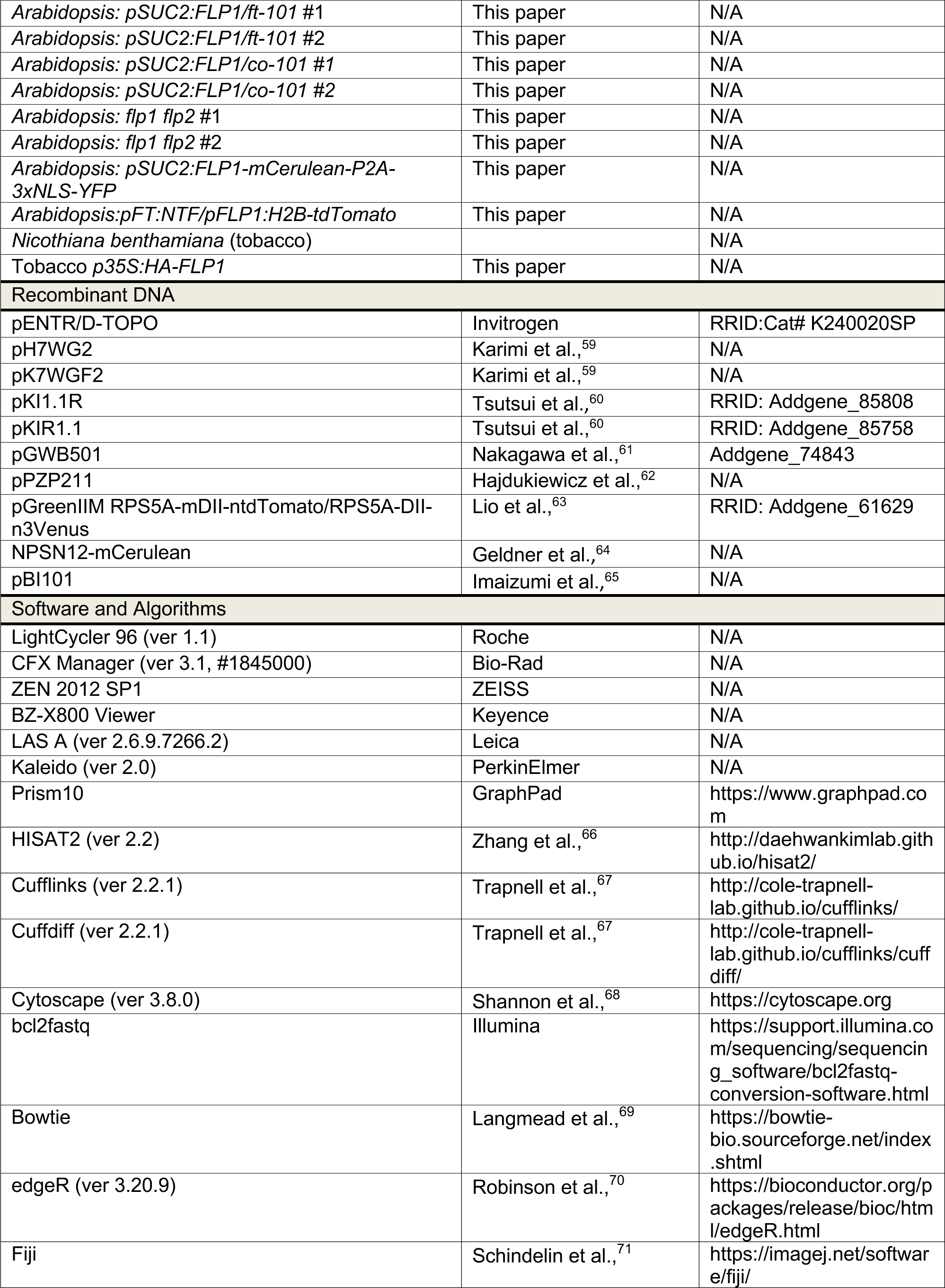
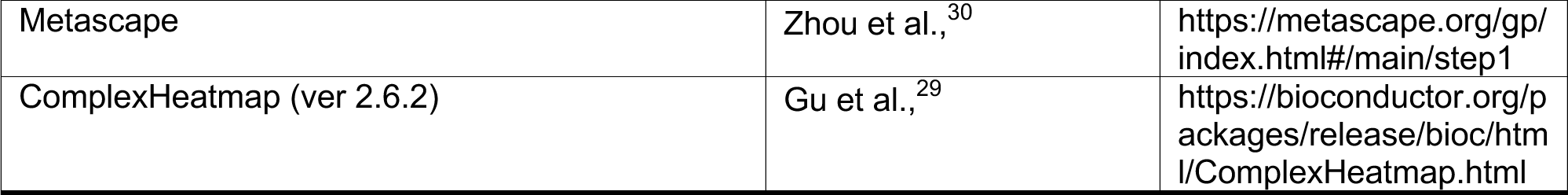

| REAGENT or RESOURCE | SOURCE | IDENTIFIER |
| --- | --- | --- |
| Antibodies |  |  |
| Monoclonal ANTI-FLAG® M2 antibody produced in mouse | Sigma-Aldrich | Cat# F1804 |
| Bacterial and Virus Strains |  |  |
| *Escherichia coli* strain DH10B | Thermo Fisher | Cat# EC0113 |
| *Escherichia coli* strain DH5α | Nippon gene | 310-06236 |
| *Escherichia coli* strain *ccdB* Survival 2 T1<sup>R</sup> | Thermo Fisher | A10460 |
| *Agrobacterium tumefaciens* strain GV3101 | Imaizumi and Tada lab stock | N/A |
| Chemicals, Peptides, and Recombinant Proteins |  |  |
| Polyoxyethylene (10) tridecyl ether | Sigma-Aldrich | Cat# P2393 |
| Sodium deoxycholate (DOC) | Sigma-Aldrich | Cat# 30970 |
| Polyoxyethylene(23)lauryl ether (Brij-35) | Sigma-Aldrich | Cat# 16005 |
| IGEPAL® CA-630 | Sigma-Aldrich | Cat# I3021 |
| Phenylmethylsulfonyl fluoride (PMSF) | Sigma-Aldrich | Cat# P7626 |
| Tween 20 | Fisher Scientific | Cat# BP337 |
| Triton X-100 | Fisher Scientific | Cat# BP151 |
| EGTA | Fisher Scientific | Cat# O2783 |
| DTT | Thermo Fisher | Cat# R0861 |
| Tris | Fisher scientific | Cat# BP152 |
| KCl | Fisher scientific | Cat# BP366 |
| MgCl<sub>2</sub> | Fisher scientific | Cat# AC223210025 |
| Dynabeads™ Protein G for Immunoprecipitation | Invitrogen | Cat# 10003D |
| Chloramphenicol | PhytoTechLab | Cat# C252 |
| Cycloheximide | Sigma-Aldrich | Cat# C1988 |
| Critical Commercial Assays |  |  |
| YourSeq Strand-Specific mRNA Library Prep Kit | Amaryllis Nucleics | Cat# 23001 |
| PrimeScript™ RT reagent Kit | Takara Bio. | Cat# RR037A |
| KAPA SYBR FAST qPCR Master Mix (2x) kit | Roche | Cat# KK4602 |
| NEBNext Ultra RNA Library Prep Kit for Illumina | NEB | Cat# E7530L |
| Cell-Free Protein Synthesis Kit | NUProtein | Cat# PSS4050 |
| AlphaScreen™ FLAG™ (M2) Detection Kit | PerkinElmer | Cat# 6760613C |
| Deposited Data |  |  |
| TRAP-seq | This paper | DRA016554 |
| RNA-seq | This paper | DRA016641 |
| Experimental Models: Organisms/Strains |  |  |
| *Arabidopsis thaliana* ecotype Columbia |  | N/A |
| *Arabidopsis*: *pFT:FLAG-GFP-RPL18* | This paper | N/A |
| *Arabidopsis*: *pSUC2:FLAG-GFP-RPL18* | Mustroph et al.,<sup>23</sup> | CD3-2409 |
| *Arabidopsis*: *pRBCS1A:FLAG-GFP-RPL18* | Mustroph et al.,<sup>23</sup> | CD3-2407 |
| *Arabidopsis*: *pCER5:FLAG-RPL18* | Mustroph et al.,<sup>23</sup> | CS66057 |
| *Arabidopsis*: *p35S:FLAG-GFP-RPL18* | Mustroph et al.,<sup>23</sup> | CD3-2403 |
| *Arabidopsis*: *pSUC2:FLP1 #1* | This paper | N/A |
| *Arabidopsis*: *pSUC2:FLP1 #2* | This paper | N/A |
| *Arabidopsis*: *pUFO:FLP1 #11* | This paper | N/A |
| *Arabidopsis*: *pUFO:FLP1 #13* | This paper | N/A |
| *Arabidopsis*: *pFT:GUS* | Takada and Goto,<sup>21</sup> | N/A |
| *Arabidopsis*: *p35S:GFP-FLP1* | This paper | N/A |
| *Arabidopsis*: *flp1-1* | This paper | N/A |
| *Arabidopsis*: *flp1-2* | This paper | N/A |
| *Arabidopsis*: pSUC2:FLP1/ft-101 #1 | This paper | N/A |
| *Arabidopsis*: pSUC2:FLP1/ft-101 #2 | This paper | N/A |
| *Arabidopsis*: pSUC2:FLP1/co-101 #1 | This paper | N/A |
| *Arabidopsis*: pSUC2:FLP1/co-101 #2 | This paper | N/A |
| *Arabidopsis*: flp1 flp2 #1 | This paper | N/A |
| *Arabidopsis*: flp1 flp2 #2 | This paper | N/A |
| *Arabidopsis*: pSUC2:FLP1-mCerulean-P2A-3xNLS-YFP | This paper | N/A |
| *Arabidopsis*:pFT:NTF/pFLP1:H2B-tdTomato | This paper | N/A |
| *Nicotiana benthamiana* (tobacco) |  | N/A |
| Tobacco p35S:HA-FLP1 | This paper | N/A |
| Recombinant DNA |  |  |
| pENTR/D-TOPO | Invitrogen | RRID:Cat# K240020SP |
| pH7WG2 | Karimi et al.,<sup>59</sup> | N/A |
| pK7WGF2 | Karimi et al.,<sup>59</sup> | N/A |
| pKI1.1R | Tsutsui et al.,<sup>60</sup> | RRID: Addgene_85808 |
| pKIR1.1 | Tsutsui et al.,<sup>60</sup> | RRID: Addgene_85758 |
| pGWB501 | Nakagawa et al.,<sup>61</sup> | Addgene_74843 |
| pPZP211 | Hajdukiewicz et al.,<sup>62</sup> | N/A |
| pGreenIIIM RPS5A-mDII-ntdTomato/RPS5A-DII-n3Venus | Lio et al.,<sup>63</sup> | RRID: Addgene_61629 |
| NPSN12-mCerulean | Geldner et al.,<sup>64</sup> | N/A |
| pBI101 | Imaizumi et al.,<sup>65</sup> | N/A |
| Software and Algorithms |  |  |
| LightCycler 96 (ver 1.1) | Roche | N/A |
| CFX Manager (ver 3.1, #1845000) | Bio-Rad | N/A |
| ZEN 2012 SP1 | ZEISS | N/A |
| BZ-X800 Viewer | Keyence | N/A |
| LAS A (ver 2.6.9.7266.2) | Leica | N/A |
| Kaleido (ver 2.0) | PerkinElmer | N/A |
| Prism10 | GraphPad | https://www.graphpad.com |
| HISAT2 (ver 2.2) | Zhang et al.,<sup>66</sup> | http://daehwankimlab.github.io/hisat2/ |
| Cufflinks (ver 2.2.1) | Trapnell et al.,<sup>67</sup> | http://cole-trapnell-lab.github.io/cufflinks/ |
| Cuffdiff (ver 2.2.1) | Trapnell et al.,<sup>67</sup> | http://cole-trapnell-lab.github.io/cufflinks/cuffdiff/ |
| Cytoscape (ver 3.8.0) | Shannon et al.,<sup>68</sup> | https://cytoscape.org |
| bcl2fastq | Illumina | https://support.illumina.com/sequencing/sequencing_software/bcl2fastq-conversion-software.html |
| Bowtie | Langmead et al.,<sup>69</sup> | https://bowtie-bio.sourceforge.net/index.shtml |
| edgeR (ver 3.20.9) | Robinson et al.,<sup>70</sup> | https://bioconductor.org/packages/release/bioc/html/edgeR.html |
| Fiji | Schindelin et al.,<sup>71</sup> | https://imagej.net/software/fiji/ |
| Metascape | Zhou et al.,<sup>30</sup> | https://metascape.org/gp/index.html#/main/step1 |
| ComplexHeatmap (ver 2.6.2) | Gu et al.,<sup>29</sup> | https://bioconductor.org/packages/release/bioc/html/ComplexHeatmap.html |

### Resource availability

### Lead contact

Further information and requests for resources and reagents should be directed to and will be fulfilled by the lead contact, Takato Imaizumi.

## Materials availability

All data required to support the claims of this paper are included in the main and supplemental information.

All reagents generated in this study are available on request from the lead contact.

## Data and code availability

The TRAP-seq and RNA-seq data have been deposited in the DNA Data Bank of Japan (DDBJ) Sequence Read Archive under accession number, DRA016554 and DRA016641, respectively.

This paper does not report original code.

Any additional information required to reanalyze the data reported in this paper is available from the lead contact upon request.

## Experimental model and study participant details

Plants were grown either on 1x Linsmaier and Skoog (LS) media plates without sucrose containing 0.8% (w/v) agar or Sunshine Mix 4 soils (Sun Gro Horticulture) as described in Song *et al*.^3^ with minor adjustments. Surface sterilized seeds were sown on media plates or soils at a low density to avoid shading effects from neighboring plants. Seeds sown on plates or soils were kept in 4 °C for at least for 2 days for stratification prior to transferring to the incubators.

Plants on plates were grown in plant incubators (Percival Scientific and Nippon Medical & Chemical Instruments) at constant 22 °C under white fluorescent or LED light with a fluence rate of 90-110 µmol m^−2^ sec^−1^. In long-day (LD; 16-hour light and 8-hour dark) conditions without far-red light (FR) LED, the red/far-red (R/FR) ratio was >2.0. In LD+FR and short-day (SD+FR; 8-hour light and 16-hour dark) conditions in which the R/FR ratios were adjusted to approximately 1.0 using dimmable FR LED light (Fluence Bioengineering and Co. Fuji Electric) connecting to the timer switch. To diffuse and dim FR light from the light source, the FR LED light fixture was covered with one-layer white printer paper. The R/FR ratio was frequently checked using spectrophotometers and R/FR sensors (Spectrum Technologies, Apogee Instruments, and Nippon Medical & Chemical Instruments) to make sure that the proper R/FR rates were maintained.

For soil growth, soils were supplemented with a slow-release fertilizer (Osmocote 14-14- 14, Scotts Miracle-Gro) and a pesticide (Bonide, Systemic Granules) and filled in standard flats with inserts (STF-1020-OPEN and STI-0804, T.O. Plastics). For flowering time measurements, plants on soil flats were grown in PGC Flex reach-in models with broad-spectrum white light and FR LED lighting (Conviron) at constant 22 °C. For LD and LD+FR conditions, the fluorescence rate was set to 100 µmol m^-2^ sec^-1^; for SD+FR conditions, 200 µmol m^−2^ sec^−1^. The number of rosette and cauline leaves of soil-grown plants was counted when the length of the bolting stem became about 1 cm. For inflorescence stem length measurements, plants were grown in the same Percival Scientific plant incubators used for sterile growth in plates. For inflorescence stem elongation analysis, the plants were grown under LD+FR conditions. The total length of the bolting stems was recorded 4, 7, and 8 days after the formation of the visible flower bud.

## Method details

### Molecular cloning and plant materials

All *Arabidopsis thaliana* transgenic plants and mutants are Col-0 background. To generate *pSUC2:FLP1* and *pUFO:FLP1* transgenic lines, the full length of *FLP1* cDNA was amplified by the primers (5’- CACCATGTCTGGTGTGTGGGTATTCAACA -3’ and 5’-TACTACATGTCACGGACATGGAAG-3’) and cloned into the pENTR/D-TOPO vector (Invitrogen). Once sequences of *FLP1* cDNA were verified, *FLP1* cDNA was transferred to the binary GATEWAY vectors, pH7SUC2 and pH7UFO, both of which the *35S* promoter in pH7WG2 ^59^ was replaced by 0.9 kb of *SUC2* and 2.6 kb of *UFO* promoters, respectively. The *pSUC2:FLP1* construct was transformed into wild-type (WT) plants possessing *pFT:GUS* reporter gene,^21^ *ft-101,* and *co-101* plants. The *pUFO:FLP1* construct was transformed into WT. To generate *p35S:GFP-FLP1* lines, the *FLP1* cDNA in pENTR/D-TOPO was transformed into the vector pK7WGF2,^59^ which has an N-terminal *GFP* gene. The *p35S:GFP-FLP1* construct was transformed into WT. All transgenic overexpressor lines have single insertions of the constructs and are homozygote when utilized.

The *flp1-1* and *flp1-2* mutant lines were generated by CRISPR-Cas9-mediated gene editing of WT plants using pKI1.1R plasmid.^60^ To make pKI1.1R containing single gRNA targeting the *FLP1* coding region, the annealed primers (5’-ATTGTAACCAGCAGCAGAGGATG-3’ and 5’- AAACCATCCTCTGCTGCTGGTTA-3’) were inserted into pKI1.1R as previously described.^60^ To generate the *flp1 flp2* double mutants, *FLP2* was mutated in the *flp1-2* mutant background using the pKIR1.1 construct containing single gRNA targeting the *FLP2* coding region made by the primer set (5’-ATTGTAAGATGAGACGGCTTCAC-3’ and 5’- AAACGTGAAGCCGTCTCATCTTA-3’). All gene-editing T-DNAs were genetically removed from the genome-edited mutant lines by the T3 stage.

To synthesize the *pFT:NTF* construct, we used pENTR/D-TOPO harboring 5.7 kb upstream of *FT*, nuclear-targeting fusion protein (NTF),^36^ and *FT* genomic and downstream regions. The construct was subsequently transformed into the GATEWAY destination vector pGWB501 ^61^. Due to the technical difficulty in cloning 5.7 kb promoter region within single PCR, several shorter fragments were cloned first and ligated together to generate the 5,722 bp of the *FT* promoter. First, 3,553 to 5,722 bp upstream region containing *Spe*I, *Sac*I, and *Xho*I sites at 3’ end was amplified by PCR using primers (5’- CACCATTTGCTGAACAAAAATCTATTAC-3’ and 5’- <u>CTCGAGGAGCTCACTAGT</u>ATATAAGAGATATGTGTCAATCC-3’, restriction enzyme recognition sequences in the primers are underlined hereafter) and cloned into pENTR/D-TOPO. Subsequently, 1,719 to 3,923 bp upstream of *FT* was amplified using primers (5’- ACTTGGATATGATGTTAAGTATC-3’ and 5’- TATTTTCTACTAATTTTAGTTACACAC-3’), and 1,754 to 3,834 bp upstream region was inserted using *Spe*I and *Sac*I sites. Next, 1 to 1,870 bp upstream region was amplified using primers (ATTAATCTTGTCTGCGACTGCGACC-3’ and 5’- AG<u>CTCGAG</u>CTTTGATCTTGAACAAACAGGTGGTTTC-3’), and 1 to 1,754 bp upstream region was inserted using *Sac*I and *Xho*I sites. To fuse *FT* promoter and *NTF*, *NTF* sequences containing *Xho*I sites at 5’ and 3’ ends were amplified by PCR using primers (5’- AT<u>CTCGAG</u>ATGGATCATTCAGCGAAAAC-3’ and 5’- GA<u>CTCGAG</u>TCAAGATCCACCAGTATCCTC -3’) and inserted into pENTR/D-TOPO carrying 5.7 kb upstream of *FT* promoter using *Xho*I sites. Finally, *FT* genomic fragment (Chr1: 24,331,510–24,335,529) containing *FT* gene body (exons, introns and 3’-UTR) as well as following 1,822 bp of the 3’ sequences, including *FT* regulatory element Block E,^72^ were amplified using primers (5’-<u>GGCGCGCC</u>ATGTCTATAAATATAAGAGA-3’ and 5’- <u>GGCGCGCC</u>TATTAAACTAGCAGTCAAA-3’) and inserted into the *Asc*I site existed in the pENTR/D-TOPO already containing *FT* promoter and NTF genes. The resulting pENTR/D- TOPO construct containing both 5’ and 3’ sequences of *FT* and the *NTF* gene was introduced into pGWB501 binary vector (referred to as *pFT:NTF*). The *pFT:NTF* construct was transformed into WT plants containing *pACT2:BirA* ^36^ for future use for INTACT (isolation of nuclei tagged in specific cell types) approach.

To generate *pFT:FLAG-GFP-RPL18* for TRAP-seq, we amplified the *FLAG-GFP-RPL18* fragment using primers containing a *Sal*I site (5’- ACGC<u>GTCGAC</u>GGTACCTATTTTTACAACAA-3’) and an *Asc*I site (5’-<u>GGCGCGCC</u>CCGGCCGCCGTGCT-3’) and cloned into *Xho*I and *Asc*I sites (note *Sal*I- and *Xho*I-cut overhands are the same) in the same pENTR/D-TOPO used for generating *pFT:NTF,* which already contains the 5.7 kb upstream of *FT* promoter. Then the same 3’ *FT* genomic regions (spanning from *FT* gene body to the Block E sequences) was inserted into the *Asc*I site to generate the entire *pFT:FLAG-GFP-RPL18* construct. The *pFT:FLAG-GFP-RPL18* construct harboring from the *FT* promoter to genomic *FT* sequences was transferred to the pGWB401 binary vector and transformed into WT.

The *pFLP1:H2B-tdTomato* construct was made by swapping the heat-shock promoter (*HS*) of pPZP211 ^62^ *HS:H2B-tdTomato* with 1,853 bp of the *FLP1* promoter. The *FLP1* promoter was amplified by the primers (5’- <u>CCTGCAGG</u>AGAATCTGATGATGTTGAGGCTAGTCG-3’ and 5’- <u>GTCGAC</u>GACTCCGTTTTTGTTGAATACCCACAC-3’) and inserted using *Sbf*I and *Sal*I restriction enzyme sites. This construct was transformed into the already established *pFT:NTF* plants.

To synthesize the *pSUC2:mCerulean-P2A-3xNLS-YFP* construct, we first cloned multiple cloning sites of pRTL2 ^73^ into pENTR/D-TOPO (denoted pENTR-MCS). Next, *3xNLS-YFP* derived from pGreenIIM RPS5A-mDII-ntdTomato/RPS5A-DII-n3Venus ^74^ was cloned into *Bam*HI and *Xba*I sites of the pENTR-MCS. Using NPSN12-mCerulean ^75^ as a template, *mCerulean-P2A* was amplified by PCR using primers (5’- <u>GAGCTC</u>CTTGTACAGCTCGTCCATGCC-3’ and 5’- <u>GGATCC</u>CCCATAGGTCCAGGATTTTCTTCAACATCTCCAGCTTGCTTAAGAAGAGAA AAATTAGTAGCGCCGCTGCCCATATGCTTGTACAGCTCGTCCATGCCGAG-3’), and inserted into *Sac*I and *Bam*HI sites of the pENTR-MCS clone already containing *3xNLS-YFP* cDNA. To generate an in-frame fusion of *FLP1* and *mCerulean* genes, the full length of *FLP1* cDNA, whose stop codon was replaced with *Sac*I site, was amplified using the primers (5’-CACCATGTCTGGTGTGTGGGTATTCAACA-3’ and 5’-GAGCTCCATGTCACGGACATGGAAGA-3’) and cloned into the different pENTR/D-TOPO plasmid. This *FLP1* cDNA without the stop codon was excised by *Not*I and *Sac*I and inserted into *Not*I and *Sac*I sites of the pENTR-MCS containing *mCerulean-P2A-3xNLS-YFP* cDNA. Finally, the pENTR-MCS containing fused *FLP1-mCerulean-P2A-3xNLS* gene was transformed into the pH7SUC2 vector. The *pSUC2:FLP1-mCerulean-P2A-3xNLS-YFP* construct was transformed into WT plants.

To generate the *FLP1:GUS* construct, 1,858 bp of *FLP1* promoter was amplified by the primers (5’-AAAGCTTGAAGCAGAATCTGATGATGTTGAGGCTAGTCGTTATCAC-3’ and 5’- CGGATCCGTTGTAAGGATTCTCCACCAGCCTCATGACTCCG-3’) and inserted into *Hind*III and *Bam*HI sites of pBI101.^65^ The resulting *FLP1:GUS* construct was transformed into WT plants.

### Translating Ribosome Affinity Purification sequencing (TRAP-seq)

All transgenic lines for TRAP-seq, *p35S:FLAG-GFP-RPL18*, *pRBCS1A:FLAG-GFP- RPL18*, *pCER5:FLAG-RPL18*, *pSUC2:FLAG-RPL18*^23^ and *pFT:HF-GFP-RPL18* (Figure 1A) were grown in 1xLS plates as described above. Ten-day-old plants grown under LD conditions were transferred to LD+FR or kept growing in the same LD chamber for additional 4 days.

Whole tissues including roots of 14-day-old plants were quickly harvested and frozen in liquid nitrogen. TRAP RNA was isolated as previously described.^23^ Pulverized tissues were homogenized in approximately five-time volume (w/v) of polysome extraction buffer [0.2 M Tris-HCl pH 9.0, 0.2 M KCl, 0.025 M EGTA, 0.035 M MgCl_2_, 1% (w/v) Brij-35, 1% (v/v) Triton X-100, 1% (v/v) Igepal CA 630, and 1% (v/v) Tween 20, 1% (v/v) Polyoxyethylene (10) tridecyl ether, 1mM DTT, 1mM PMSF, 50 µg/mL cycloheximide, and 50 µg/mL chloramphenicol]. Homogenized samples were centrifuged in 16,000xg for 15 min at 4 °C, subsequently, the supernatant was filtrated with Miracloth. Protein G Dynabeads (Invitrogen) coupled with anti-FLAG M2 antibody (F1804, Sigma-Aldrich) was applied to the filtrated samples transferred to 50 mL falcon tube, followed by the incubation at 4 °C for 2 hours with gentle shaking using a rocking platform. Subsequently, beads were collected with magnets for 4 min at 4 °C. After the supernatant was removed using a pipet, 6 mL of freshly prepared wash buffer (0.2 M Tris-HCl pH 9.0, 0.2 M KCl, 0.025 M EGTA, 0.035 M MgCl2, 1 mM DTT, 1 mM PMSF, 100 µg/mL cycloheximide, and 50 µg/mL chloramphenicol) were added to the beads, mixed by gentle inverting the tube, and incubated at 4 °C for 2 min with gentle shaking on rocking platform. Beads were collected with magnets at 4 °C for 4 min, supernatant was removed carefully. This washing step was repeated two more times with 6 mL and 1 mL of the washing buffer. After 1 mL of the washing buffer was removed, 450 µL of RLT Lysis buffer (Qiagen) with 4.5 µL β-mercaptoethanol was directly added to the beads and incubated for 5 min at room temperature. The rest of the process for TRAP RNA isolation was conducted following by the manufacture protocol of RNeasy Plant Mini Kit (Qiagen) except for skipping the use of shredder spin columns. RNA-seq libraries for 3’-Degital Gene expression were prepared using YourSeq Strand-Specific mRNA Library Prep Kit (Amaryllis Nucleics) following the manufacture protocol. Sequencing was performed on the Illumina Nextseq 550 platform. Approximately 20 to 66 million reads (average 38 million reads) were produced from each sample. Adapter and low-quality sequences were trimmed by Trimmomatic software (version 0.32) (LEADING:20, TRAILING: 20, MINLEN: 36).^76^ By HISAT2 software (version 2.2),^66^ the remaining reads were mapped to *Arabidopsis* genome sequences from The Arabidopsis Information Resource (TAIR, version 10) (http://www.arabidopsis.org/). Mapped reads were lined to 32,398 *Arabidopsis* annotated genes by Cufflinks software (version 2.2.1).^67^ As the expression intensities, the number of mapped reads was normalized by Reads Per Kilobase per Million mapped reads (RPKM). In transgenic plants, expressional differences between LD and LD+FR were inferred by Cuffdiff software (version 2.2.1) in 32,948 genes.^67^ Up- and down-regulated genes were defined as genes with significantly higher and lower RPKM reads (FDR < 0.05), respectively. The interactomes were constructed based on FLOR-ID ^77^ using Cytoscape.^68^

### qRT-PCR

For the time course gene expression analyses, total RNA was extracted using RNeasy Plant Mini kit (Qiagen), and cDNA synthesis and quantitative PCR (qPCR) were performed as previously described.^78^ For other tests, total RNA was extracted from plants grown on plates using NucleoSpin RNA extraction kit (Macherey-Nagel GmbH & Co.). First-strand cDNA was synthesized using 2 μg RNA and PrimeScript RT Master Mix (Takara Bio). qRT-PCR was performed using the first-strand cDNAs diluted 5-fold in water and KAPA SYBR FAST qPCR Master Mix (2x) kit (Roche) and gene-specific primers in a LightCycler 96 (Roche).

### Isopentenyl Pyrophosphate / Dimethylallyl Pyrophosphate Isomerase

(*IPP2*) and *PROTEIN PHOSPHATASE 2A SUBUNIT A3* (*PP2AA3*) were used as the internal control for normalization. For statistical tests, log_2_-transformed relative expression values were used to meet the requirements for homogeneity of variance. qPCR primers used in this study are listed in Table S1.

### GUS assay

*pSUC2:FLP1/pFT:GUS* and *pFT:GUS* plants were grown at low density on 1xLS media in LD+FR for 2 weeks or in SD+FR for either 2 weeks or 3 weeks. *pFLP1:GUS* plants were grown for 2 weeks in LD+FR. Whole plants were collected at ZT4 and immediately incubated in chilled 90% (v/v) acetone for 15 min. The samples were then rinsed twice with sterilized Milli-Q water before being immersed in 50 mM sodium phosphate buffer (pH 7.4) containing 0.5 mM potassium ferrocyanide, 0.5 mM potassium ferricyanide, 0.1% (v/v) Triton X-100, and 0.5 mM 5-Bromo-4-chloro-3-indolyl-β-D-glucuronide (X-Gluc). Immersed samples were then vacuum infiltrated to –0.09 MPa for 40 min. Vacuum pressure was gradually relieved over 10 min before samples were incubated in 37 °C for 12 hours. Samples were removed from the solution and then incubated at room temperature in a series consisting of three 30-min periods immersed in 30% (v/v) EtOH, fixing solution [50% EtOH, 5% (v/v) acetic acid, and 3.7% (v/v) formaldehyde] and 80% EtOH prior to incubation at 4 °C for 16 hours in 95% EtOH solution. Samples were then incubated at room temperature in a series consisting of six 30-min periods immersed in 80% EtOH, 50% EtOH, 30% EtOH, Milli-Q water, 25% (v/v) glycerol and 50% glycerol, and stored at 4 °C before images were captured.

### Tissue clearing and confocal microscopy

For confocal imaging of the *FLP1:H2B-tdTomato/pFT:NTF* and the *pSUC2:FLP1- mCerulean-P2A-3xNLS-YFP*, the first and second true leaves detached from 2-week-old plants were cleared following the modified method of Kurihara *et al*.^79^ Leaves were immersed in 4% (v/v) paraformaldehyde solution in a microtube. Tubes were placed in a chamber with the pressure adjusted to –0.09 MPa for 2 min. After venting not to disrupt samples, the pressure was adjusted again to –0.09 MPa for 1 hour. After pressure was slowly released, solution was removed, and the samples were washed with 1x PBS solution twice with careful pipetting.

Subsequently, ClearSee^80^ was added to microtubes with the samples and the tubes were placed on a pressure chamber adjusted to –0.09 MPa for 2 min. Pressure was released slowly and adjusted to –0.09 MPa again for 1 hour. The samples in ClearSee were placed in a box and placed in the dark at room temperature. We kept replacing ClearSee (at least twice) with a new solution until the sample became transparent. Confocal images were taken using LSM780-DUO-NLO (Zeiss) equipped with 32-channel spectral GaAsP detector and a Plan-Apochromat 63x/1.40 oil immersion objective lens. Cleared leaf tissues of *pSUC2:FLP1-mCerulean-P2A- 3xNLS-YFP* were excited with a 405 nm. Images were processed by spectral unmixing using reference fluorescent spectra of pollen of *LAT52:mTurquoise2* (YMv150) excited with a 405 nm [*LAT52:mTurquoise2* was kindly provided by Dr. Yoko Mizuta (Nagoya University)] and 3xNLS-YFP excited with a 488 nm for FLP1-mCerulean and 3xNLS-YFP, respectively. We used the following setting: GFP channel, Ex. 489 nm, and Em. 490–597 nm; RFP channel, Ex. 560 nm, and Em. 566–690 nm.

### RNA-seq

RNA-seq analysis was conducted using 2-week-old plants grown under LD+FR, LD and SD+FR conditions in biological triplicates as previously described.^81^ Total RNA was extracted using NucleoSpin RNA extraction kit, and cDNA libraries were constructed using NEBNext Poly(A) mRNA Magnetic Isolation Module, NEBNext Ultra RNA Library Prep Kit for Illumina, and NEBNext Multiplex Oligos for Illumina (New England BioLabs) according to the manufacturer’s protocol. The libraries were sequenced using the NextSeq500 sequencer (Illumina). Raw reads containing adapter sequences were trimmed using bcl2fastq (Illumina), and nucleotides with low-quality (QV < 25) were masked by N using the original script. Reads <50 bps were discarded, and the remaining reads were mapped to the cDNA reference sequence using Bowtie with the following parameters: “--all --best –strata.”^69^ Reads were then counted by transcript models. DEGs were selected based on the adjusted *P*-value (FDR < 0.05) calculated using edgeR (version 3.20.9) with default settings.^70^

### Hormone quantification

Contents of plant hormones (IAA, ABA, JA, GA_4_, JA-Ile, SA, tZ, and iP) in 2-week-old LD+FR grown plants in eight biological replicates were quantified by liquid chromatography-mass spectrometry (LC-MS) according to the previous report.^82^ Approximately 100 mg of precisely weighed *Arabidopsis* seedlings were ground in liquid nitrogen by vortexing them with a round-bottom plastic tube and 10 mm in diameter zirconia beads (Biomedical Science), each for 30 sec, repeated three times. Hormones were extracted with 4 mL of the extraction solvent consisting of 80% (v/v) acetonitrile, 19% (v/v) water, and 1% (v/v) acetic acid supplemented with deuterium- or ^13^C-labelled internal standards as described ^83^ for 1 hour at 4 °C. After centrifugation at 3,000xg for 10 min, the supernatant was corrected. The pellet was rinsed with the extraction solvent and centrifuged. Acetonitrile in the merged supernatants was evaporated by a centrifuge concentrator (Genevac miVac Quattro, SP Industries) to remain the water phase. It was applied to an Oasis HLB extraction cartridge (30 mg, Waters Corporation). The cartridge was washed with 1 mL of 1% acetic acid and eluted with 2 mL of 80% acetonitrile containing 1% acetic acid. The eluate was evaporated with a centrifuge concentrator. The resulting water phase was applied to an Oasis MCX cartridge (30 mg, Waters Corporation). After washing with 1 mL of 1% acetic acid, the acidic fraction was eluted from the Oasis MCX cartridge with 2 mL of 80% acetonitrile containing 1% acetic acid. A portion of the acidic fraction was evaporated to dryness and dissolved in 1% acetic acid and subjected to SA analysis. The basic fraction was then eluted from the Oasis MCX cartridge with 2 mL of 50% acetonitrile containing 5% (v/v) ammonia, following to a washing with 1 mL of 5% ammonia aqueous solution. The basic fraction was evaporated to dryness and dissolved in 1% acetic acid and subjected to tZ and iP analysis. The obtained acidic fraction was evaporated to remain water phase and applied to an Oasis WAX cartridge (30 mg, Waters Corporation). After washing with 1 mL of 1% acetic acid and 2 mL of 80% acetonitrile, the acidic hormones, IAA, ABA, JA, JA-Ile and GA_4_, were eluted with 2 mL of 80% acetonitrile containing 1% acetic acid. This fraction was evaporated to dryness and reconstituted in 1% acetic acid and subjected to the analysis. LC-MS analysis was carried out using Agilent 6400 mass spectrometer combined with Agilent 1260 high-performance liquid chromatography system (Agilent Technologies Inc.). The detailed condition of mass spectrometry analysis was described.^83^

### Tobacco stem measurements

The binary *p35S:HA-FLP1* plasmid was introduced in *Agrobacteria* strain GV3101. Transformation was performed using a leaf-disk transformation method based on the protocol with minor adjustments.^84^ Notably, Kanamycin-resistant transgenic plants were transferred from MSIII rooting media to Sunshine Mix #4 soil no longer than 1 week following initiation of roots, grown to maturity and subsequently self-fertilized upon flowering. Multiple homozygous, single-insertion T3 lines derived from separate successfully regenerating calluses were established for analysis of *FLP1* expression and stem growth. Seeds were liquid sterilized and stratified at 4 °C for two days in the dark, then sown onto soil in standard 4 inch pots and watered every two days. Individual plants were cultivated with sufficient space to ensure no direct over-topping of leaves in Percival Scientific plant incubators under LD (R/FR = 2.5) conditions, at 110 µmol m^−2^ sec^−1^, 22 °C, and 75% humidity for 32 days. Total length of stems, and individual internode lengths (> approximately 5 mm) from the node of cotyledon emergence to shoot apex were recorded 14, 18, 21, 25, 28, and 32 days after sowing.

### Yeast two-hybrid transcription factor (TF) library screening

The full-length *FLP1* cDNA in pENTR/D-TOPO was cloned into pDEST_GBKT7 vector^85^ by Gateway reaction and transformed into Y2HGold yeast strain (Takara Bio). The resultant yeast was cultured in synthetic defined medium (–W), transformed with *Arabidopsis* transcription factor library consisting of four 96-well plates, and spotted onto synthetic defined medium plates (–LHW) supplemented with 0.1 mM, 0.25 mM, and 0.5 mM 3-amino-1,2,4- triazole (3-AT; Fujifilm-Wako) as a first screening. The library includes 1,736 transcription factor ^86^ in pDEST_GADT7 vector ^85^ was divided into 384 wells according to their homology.

The second screening was performed for the spots showing yeast growth to determine which transcription factor was exactly positive. Yeast transformation in the first and second screenings was done by the robotic system Freedom evo 100 (Tecan).

### *In vitro* protein interaction assay by AlphaScreen system

For *in vitro* protein–protein interactions, N-terminal FLAG-tagged transcription factors and C-terminal biotinylated FLP1 (FLP1-Biotin) were synthesized using the Cell-Free Protein Synthesis Kit (BioSieg) as described previously.^44^ The primers used for protein synthesis are listed in Table S2. The biotinylated proteins were dialyzed against 1x PBS (137 mM NaCl, 8.1 mM Na_2_HPO_4_, 2.68 mM KCl, and 1.47 mM KH_2_PO_4_) at 4 °C for 24 hours. The synthesized proteins were confirmed by immunoblotting with an anti-FLAG antibody (Fujifilm-Wako) or Streptavidin-HRP (Cell Signaling Technology).

To evaluate the interactions between FLAG-tagged proteins and FLP1-Biotin, the AlphaScreen system [FLAG (M2) Detection Kit, PerkinElmer] was performed following the method of Nomoto *et al.*^45^ A 25 µL reaction buffer [1x Control buffer, 0.01% (w/v) Tween20 (Sigma-Aldrich), 0.1% (w/v) BSA (Sigma-Aldrich), 500 ng anti-FLAG Acceptor beads, 500 ng

Streptavidin-coated Donor beads, produced FLAG-tagged protein and biotinylated protein] were incubated in a 384-well plate at 22 °C for 12 hours. AlphaScreen Units were measured using EnSight Multimode Plate Reader (PerkinElmer).

### Bimolecular fluorescence complementation (BiFC) assay in *Nicotiana benthamiana* leaf epidermal cells

BiFC assay was conducted using the split sfGFP system.^87^ BiFC assay in *N. benthamiana* leaf was conducted as previously described with slight modifications.^78^ *Agrobacterium* carrying binary vectors was cultured in LB media containing 20 µM acetosyringone and antibiotics overnight and subsequently precipitated at 4,000 rpm at room temperature. *Agrobacterium* pellet was resuspended using the inoculation buffer (10 mM MgCl_2_, 10 mM MES-KOH pH5.6, and 100 µM acetosyringone) and OD_600_ was adjusted to 0.3. After incubation for 3 hours at room temperature, *Agrobacterium* was infiltrated into the abaxial side of 3 to 5-week-old tobacco leaves using 1 mL syringe. *Agrobacterium* carrying pMDC99 RPS5A:H2B-tdTomato^88^ and pBIN61 P19 (OD_600_ = 0.2, respectively) were co-inoculated for marking nuclei and suppressing gene silencing. 3 days after inoculation, GFP and RFP signals were observed using fluorescence microscopes KEYENCE BZ-X800 and Leica DMI 3000B.

### Micrografting experiments

For the I-shape micrografting, the shoot of *p35S:GFP-FLP1* was connected through the hypocotyl to the WT rootstock using the micrografting device according to Tsutsui *et al*.^89^ The grafted plants were kept in continuous light and transferred to LD+FR conditions a day before collection. The movement of GFP-FLP1 to the root tip of the WT stock was analyzed 7 days after grafting (DAG). For the Y-shape micrografting, autoclaved nylon membranes Hybond-N+ (GE Healthcare) were placed on the 1% (w/v) agar Murashige–Skoog (MS) media plates (half-strength MS media containing 0.05% (w/v) MES hydrate and 0.5% (w/v) sucrose), and the seeds of *flp1-1* and *pSUC2:FLP1* #2 were sown directly on the membrane with low density. After stratification for 2 days at 4 °C, plates were placed under continuous light for 4 days, and shoots of *flp1-1* and *pSUC2:FLP1* #2 (scion) were cut and inserted into a hypocotyl of *flp1-1* seedlings (stock) on nylon membrane placed on 2% agar 1/2x MS media. At 4 DAG, the adventitious root emerging from the scion was removed; at 6 DAG, grafted plants were transferred to 1% agar MS plates. Successfully grafted plants were kept growing on the vertically oriented plates under LD conditions. At 11 DAG, all leaves in the *flp1* stock were carefully removed. Thanks to this, the scion became the only part where the leaf-initiated phloem streaming occurs, which may have enhanced material transport rates from the scion to the stock. At 14 DAG, the shoot apex tissue containing the SAM from the *flp1-1* stock was dissected individually under the stereomicroscope for gene expression analysis. At 18 DAG, the length of SAM of the *flp1-1* stock was measured as shown in Figure 7M.

### Statistical analysis

Pairwise comparison was conducted with Student’s *t*-test or Welch’s *t*-test under Microsoft Excel environment. The equality of two variances was tested with *F*-test prior to *t*-test. Statistical tests in the flowering time measurements using *ft-101* and *co-101* background transgenic and mutant lines were performed using one-way analysis of variance (ANOVA) followed by Tukey’s multiple comparisons. Phytohormones levels in three different genotypes under LD+FR and SD+FR were compared using two-way ANOVA followed by Tukey’s multiple comparisons. All multiple comparison tests were conducted using GraphPad Prism 10 (GraphPad Software).

## Acknowledgments

We thank Y. Kodama for technical advice for microscopy, J.L. Nemhauser and Y. Mizuta for providing research materials, M. Ashikari, H. Tsuji, K. Nagai, and M. Mizutani for scientific discussion, and M. Ohme-Takagi for critical reading of the manuscript. We also thank T. Niwa, A. Furuta, F. Tobe, and S. Ishikawa for technical assistance. This research was supported by grants from the National Institutes of Health grant (R01GM079712 to J.T.C., C.Q., and T.I.), Japan MEXT KAKENHI grant (JP20H05910 and JP22H04978 to T.I., and JP20H05905 to T.M.), and National Research Foundation of Korea grant (NRF-2021R1A4A1032888 to Y.H.S.).

## Author contributions

Conceptualization: H.T., N.L, and T.I.

Methodology: H.T., N.L., A.K.H., M.Notaguchi

Investigation: H.T., N.L., A.K.H., S.P., M.Notaguchi, K.Y., K.S., Y.K., S.U., B.L.R.D., W.G.A., T.S., T.Matsuura, I.C.M., J.T.C., T.I.

Formal analysis: H.T., N.L., K.S., K.H., J.T.C.

Data curation: K.S., T.S., K.H., J.T.C.

Validation: H.T., N.L., A.K.H., S.P., M.Notaguchi, S.U., W.G.A., T.S., T.Matsuura, T.I.

Visualization: H.T., N.L., T.I.

Funding acquisition: T.Matsushita, Y.H.S., J.T.C., C.Q., T.I.

Resources: S.I., N.M., D.K.

Project administration: H.T., N.L., C.Q., T.I.

Supervision: M.Notaguchi, T.Matsushita, Y.H.S., Y.S., M.Nomoto, Y.T., K.H., C.Q., T.I.

Writing – original draft: H.T., N.L, A.K.H, W.G.A, and T.I.

Writing – review & editing: M.Notaguchi, K.S., Y.K., S.U., S.I., T.S., I.C.M., N.M., D.K., Y.H.S., Y.S., M.Nomoto, Y.T., K.H., J.T.C., and C.Q. reviewed and commented on the manuscript.

## Declaration of interests

Authors declare that they have no competing interests.

## Notes

### Competing Interest Statement

The authors have declared no competing interest.

