## Supplemental information for "Florigen-producing cells express FPF1-LIKE PROTEIN 1 that accelerates flowering and stem growth in long days with sunlight red/far-red ratio in *Arabidopsis*"

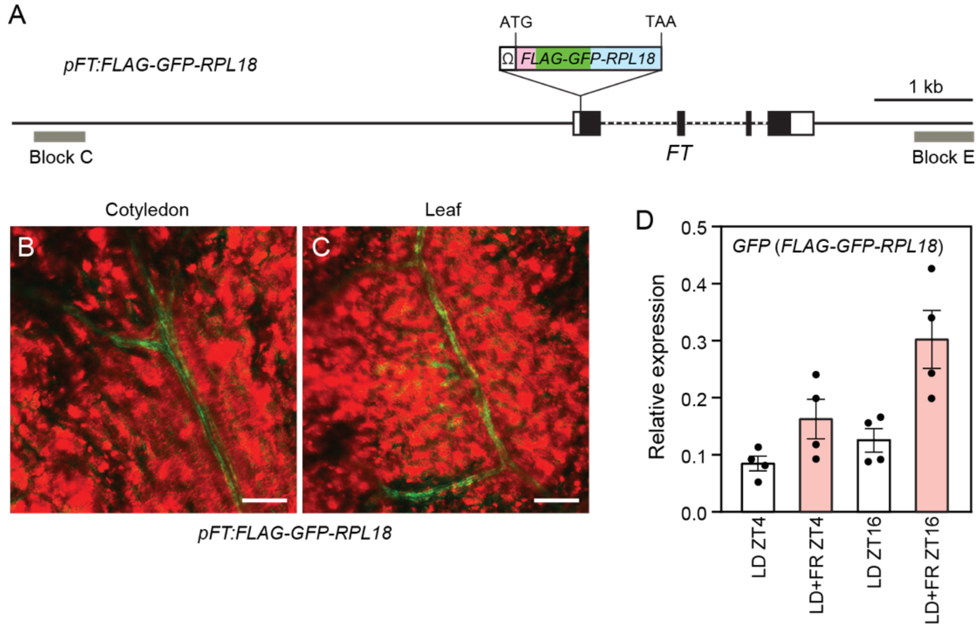


**Figure S1. The transgenic line used in the TRAP-seq analysis in *FT*-producing cells.**
(A) A diagram of the *pFT:FLAG-GFP-RPL18* construct. The *FLAG-GFP-RPL18* gene was inserted between *FT* promoter and gene in the *pFT:FLAG-GFP-RPL18* line. This construct has both Block C and Block E, upstream and downstream essential enhancer elements for *FT* transcription, respectively. White boxes indicate 5'- and 3'-UTRs, and black boxes are exons. Scale bar,1 kb.

(B and C) Vasculature-specific expression of FLAG-GFP-RPL18 protein in cotyledons (B) and true leaves (C) of *pFT:FLAG-GFP-RPL18* plants grown for 2 weeks in LD+FR. The green color is GFP fluorescence, while the red color is bright field. Scale bars, 100 µm.

(D) The expression levels of *FLAG-GFP-RPL18* transcripts in 2-week-old *pFT:FLAG-GFP-RPL18* plants grown under LD and LD+FR conditions. Samples were harvested in the morning (ZT4) and the evening (ZT16). qRT-PCR primers for *GFP* sequences (Table. S1) were used to detect the relative expression levels of *FLAG-GFP-RPL18* transcripts. Each dot indicates a biological replicate ($n$ = 4).

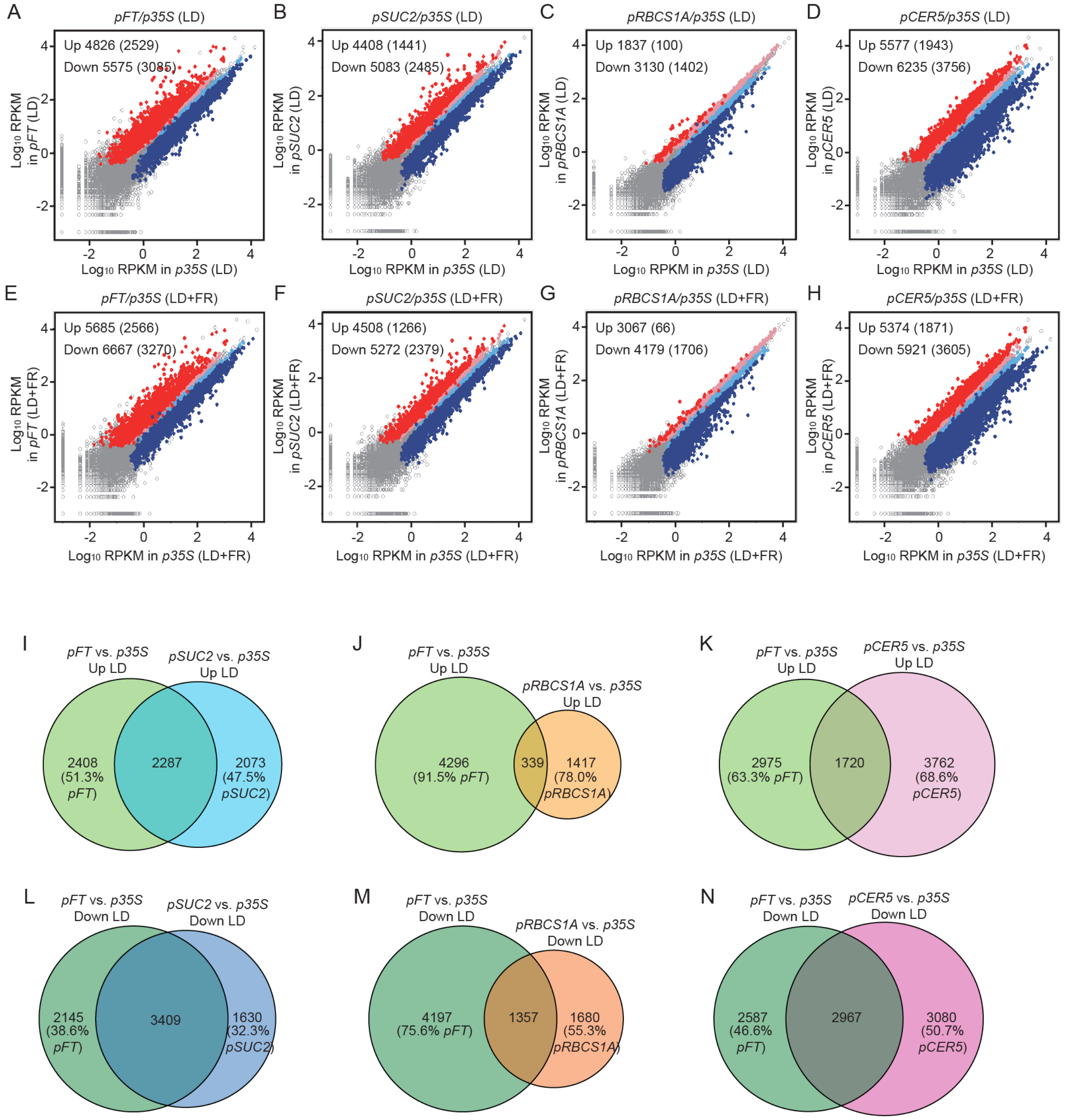


**Figure S2. Tissue/cell-specific differentially translated genes in our TRAP-seq analysis.** (A–H) Comparison between the translatome datasets of *p35S:FLAG-GFP-RPL18* and other tissue-specific line datasets: *pFT:FLAG-GFP-RPL18* (A and E), *pSUC2:FLAG-GFP-RPL18* (B and F), *pRBCS1A:FLAG-GFP-RPL18* (C and G), *pCER5:FLAG-RPL18* (D and H) grown under LD (A–D) and LD+FR (E–H) conditions for 2 weeks. X and Y axes are $\log_{10}$-transformed RPKM values. Red and pink, and blue and light blue colors of dots indicate significantly up-regulated and down-regulated genes in comparison with *p35S* (False Discovery Rate: FDR <

0.05). Red and blue indicate that differences are 2-fold or more; pink and light blue are less than 2-fold differences. The number of genes with FDR < 0.05, and FDR < 0.05 and 2- or more fold changes in parenthesis were shown in the left corner of the plots.

(I–N) Quantitative Venn diagrams showing overlaps between translatome datasets of *pFT:FLAG-GFP-RPL18* and other tissue-specific lines under LD conditions. Up-regulated genes (I–K) and down-regulated genes (L–N) in comparison with *p35S:FLAG-GFP-RPL18*. The percentages of the translated genes in each unique portion of the diagrams are shown.

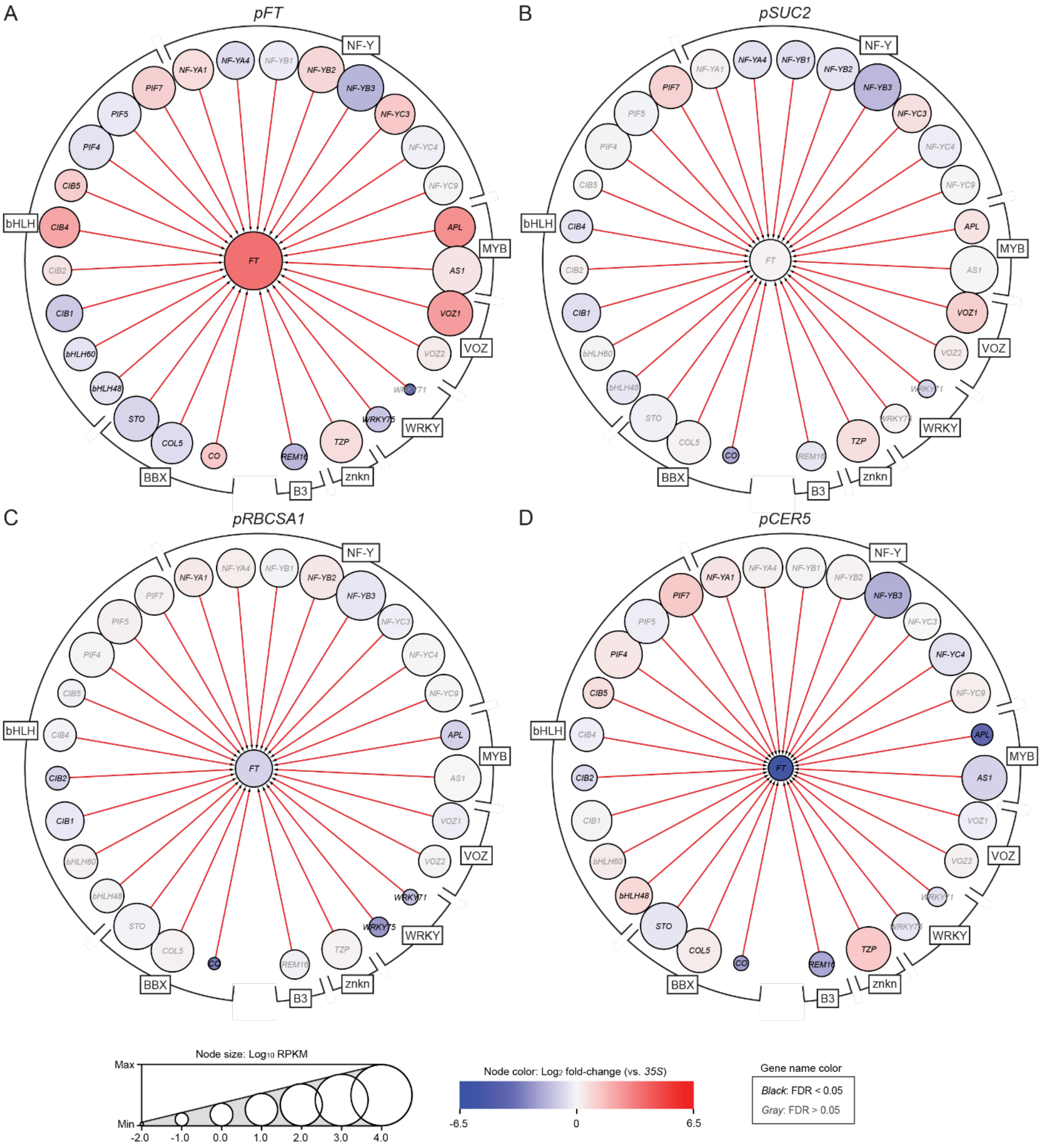


**Figure S3. Tissue/cell-specific translational levels of *FT* and *FT* positive regulator genes under LD+FR conditions.**
Interactome diagrams between *FT* and *FT* positive regulator genes with translated mRNA levels of each gene obtained from our tissue/cell-specific TRAP-seq datasets were visualized using Cytoscape. Edges connecting nodes indicate interactions between *FT* gene and *FT* positive regulators. The node circle size and its color indicate the $\log_{10}$-transformed RPKM values in each TRAP-seq dataset and the $\log_2$ fold-change against the values in the *p35S:FLAG-GFP-RPL18* data, respectively. The black color of the gene name indicates a significant difference from

*p35S:FLAG-GFP-RPL18* (FDR < 0.05), while gray indicates not significant. The TRAP-seq data were derived from the plants grown in LD+FR. *pFT:FLAG-GFP-RPL18* (A), *pSUC2:FLAG-GFP-RPL18* (B), *pRBCS1A:FLAG-GFP-RPL18* (C), and *pCER5:FLAG-RPL18* (D). Names of TF gene families are indicated.

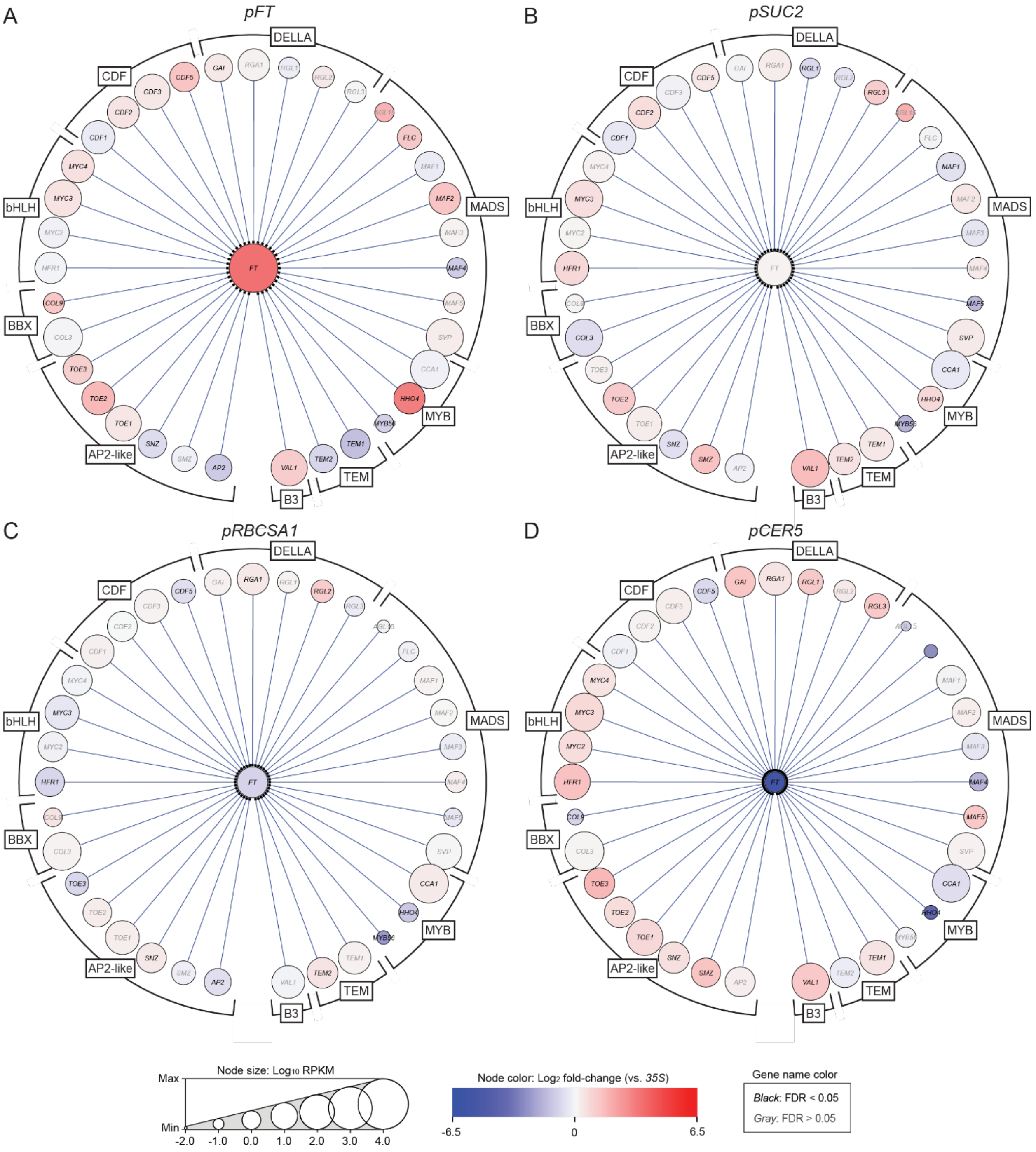


**Figure S4. Tissue/cell-specific translational levels of *FT* and *FT* repressor genes under LD+FR conditions.**

Interactome diagrams between *FT* and *FT* repressor genes with translated mRNA levels of each gene obtained from our tissue/cell-specific TRAP-seq datasets were visualized using Cytoscape. Edges connecting nodes indicate interactions between *FT* gene and *FT* negative regulators. The node circle size and its color indicate the $log_{10}$-transformed RPKM values in each TRAP-seq dataset and the $log_2$ fold-change against the values in the *p35S:FLAG-GFP-RPL18* data, respectively. The black color of the gene name indicates a significant difference from *p35S:FLAG-GFP-RPL18* (FDR < 0.05), while gray indicates not significant. The TRAP-seq data were derived from the plants grown in LD+FR. *pFT:FLAG-GFP-RPL18* (A), *pSUC2:FLAG-GFP-RPL18* (B), *pRBCS1A:FLAG-GFP-RPL18* (C), and *pCER5:FLAG-RPL18* (D). Names of TF gene families are indicated.

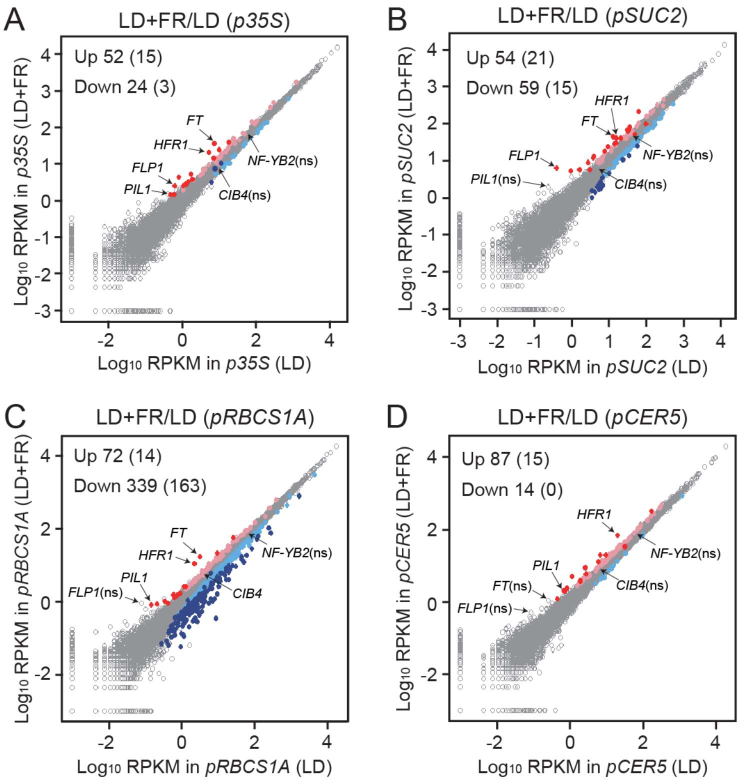


**Figure S5. Comparison between tissue/cell-specific translatome datasets under LD and LD+FR conditions.**

X and Y axes are $\log_{10}$-transformed RPKM values under LD and LD+FR, respectively. Red and pink, and blue and light blue colors of dots indicate significantly up-regulated and down-regulated genes under LD+FR (FDR < 0.05). Red and blue are 2-fold or more; pink and light blue are less than 2-fold changes. The number of genes with FDR < 0.05, and FDR < 0.05 and 2- or more fold changes in parenthesis were shown on the left corner of the plots. *p35S:FLAG-GFP-RPL18* (A), *pSUC2:FLAG-GFP-RPL18* (B), *pRBCS1A:FLAG-GFP-RPL18* (C), and *pCER5:FLAG-RPL18* (D).

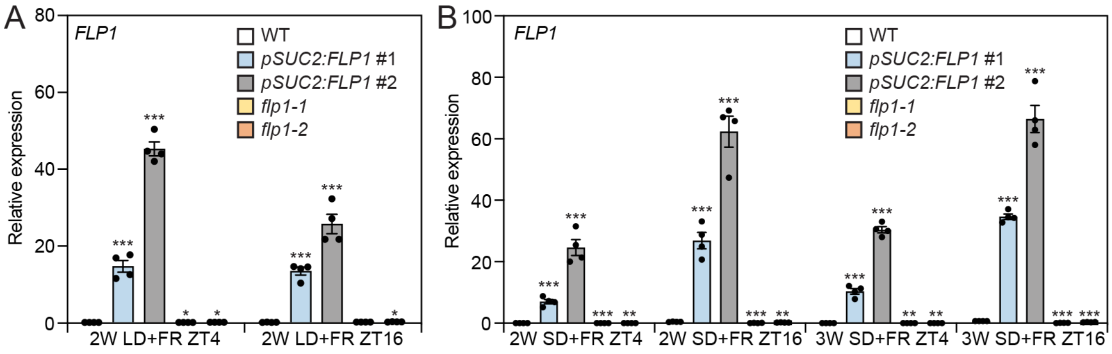


**Figure S6. *FLP1* gene expression levels in WT, *pSUC2:FLP1,* and *flp1* mutants.**
Plants were grown under LD+FR (A) and SD+FR (B) conditions for 2 or 3 weeks and harvested at ZT4 and ZT16. The results represent the means ± SEM. Each dot indicates a biological replicate ($n$ = 4). Asterisks denote significant differences from WT (*$P$<0.05; **$P$<0.01; ***$P$<0.001, $t$-test).

WT

```
              gRNA          PAM
WT 100 GGTGGTAACCAGCAGCAGAGGATGAGGAGGAAAATTCTCGTCCATCTTCCAAGCAGCGAGGTTGTGTCTT
       ||||||||||||||||||||||||||||||||||||||||||||||||||||||||||||||||||||||
WT 100 GGTGGTAACCAGCAGCAGAGGATGAGGAGGAAAATTCTCGTCCATCTTCCAAGCAGCGAGGTTGTGTCTT
```

Translation: MSGVWVFNKNGVMRLVENPYNQSAGDSSESSSSGGNQQQRMRRKILVHLPSSEVVSSYGSLEKILKNLGW
ERYYSGDNTDHLLQFHKRTSIDLISLPRDFSKFNSIHMYDIVVKNPNVFHVRDM*

*flp1-1*

```
flp1-1 100 GGTGGTAACCAGCAGCAGAGAGGTTGTGTCTTCGTACGGAAATGAGGAGGAAAATTCTCGTCCATCTTCCAAGCAGCGAGGTTGTGTCTT
           ||||||||||||||||||||                     |||||||||||||||||||||||||||||||||||||||||||||||||
    WT 100 GGTGGTAACCAGCAGCAGAGG--------------------ATGAGGAGGAAAATTCTCGTCCATCTTCCAAGCAGCGAGGTTGTGTCTT
```

**AGGTTGTGTCTTCGTACGGAA** insertion and **G** deletion > frame shift and early termination

Translation: MSGVWVFNKNGVMRLVENPYNQSAGDSSESSSSGGNQQQRGCVFVRK*
(Wrong amino acids highlighted in green)

*flp1-2*

```
flp1-2 100 GGTGGTAACCAGCAGCAGAGG----------------------------------------TTGTGTCTT
           |||||||||||||||||||||                                        |||||||||
    WT 100 GGTGGTAACCAGCAGCAGAGGATGAGGAGGAAAATTCTCGTCCATCTTCCAAGCAGCGAGGTTGTGTCTT
```

**ATGAGGAGGAAAATTCTCGTCCATCTTCCAAGCAGCGAGG (40 bp)** deletion > frame shift and early termination

Translation: MSGVWVFNKNGVMRLVENPYNQSAGDSSESSSSGGNQQQRLCLRTDHLRRS*
(Wrong amino acids highlighted in green)

**Figure S7. The mutation patterns of *flp1-1* and *flp1-2* mutants.**
Sequences of guide RNA and PAM were highlighted with blue and red letters, respectively.

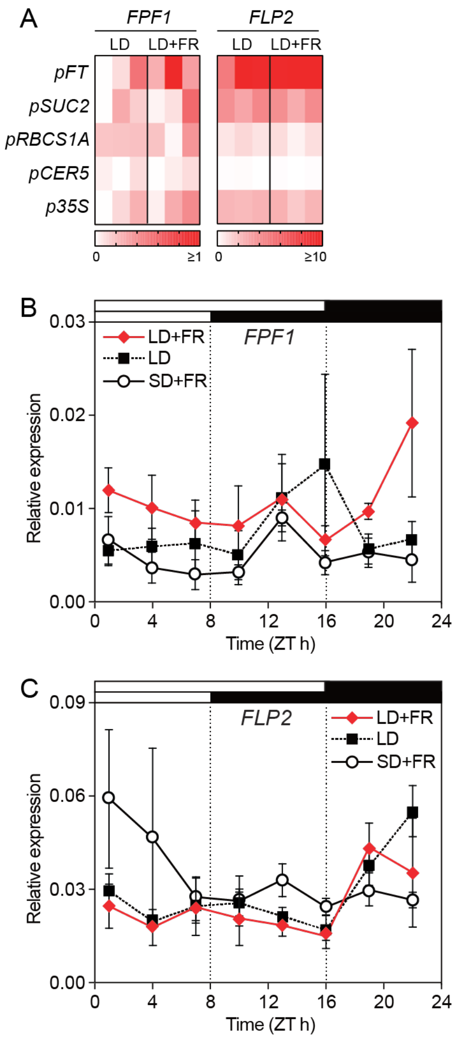


**Figure S8. Gene expression patterns of *FPF1* and *FLP2* under different photoperiod conditions.**

(A) Tissue/cell-specific translational levels of *FPF1* and *FLP2* in TRAP-seq. Letters on the left indicate the tissue/cell-specific TRAP lines used. The growth conditions are displayed on the top. The colors of squares indicate RPKM values in each biological replicate ($n = 3$).

(B and C) Time course analysis of *FPF1* (B) and *FLP2* (C) expression levels in WT plants grown under LD+FR, LD, and SD+FR conditions. The results represent the means ± SEM. ($n = 3$ biologically independent samples). White and black bars on top indicate time with light and dark.

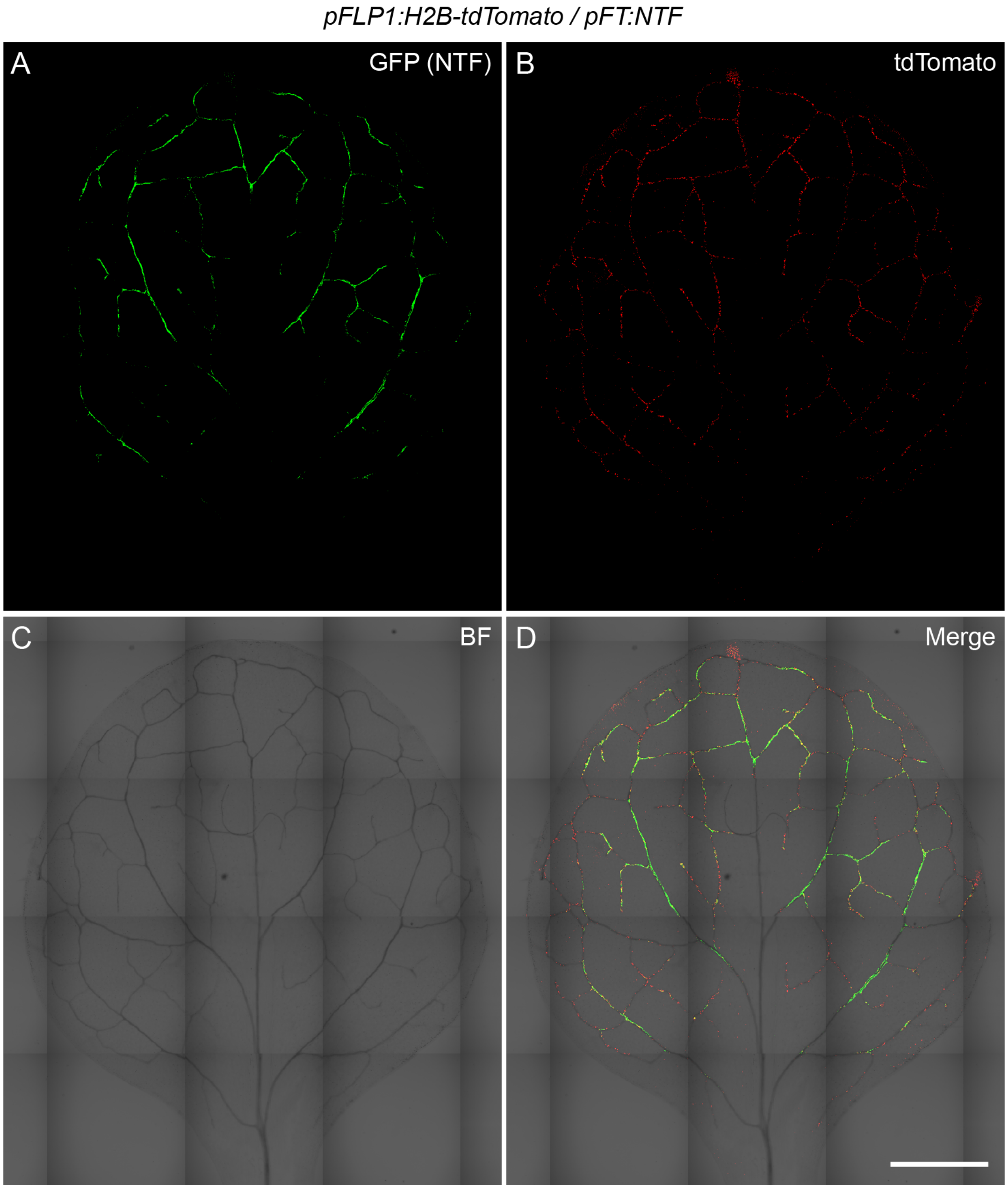


**Figure S9. Overlap between *FT* and *FLP1* promoter activities in the entire true leaf.** *pFT:NTF* (A), *pFLP1:H2B-tdTomato* (B), bright field (BF) (C), and merge of all images (D). Note, while tdTomato signals from the Histone H2B-tdTomato were exclusively in the nuclei, GFP signals from NTF were frequently observed in both nuclei and the cytosol due to weaker specificity in nuclear localization. Plants were grown for 2 weeks in LD+FR. Scale bar, 1 mm.

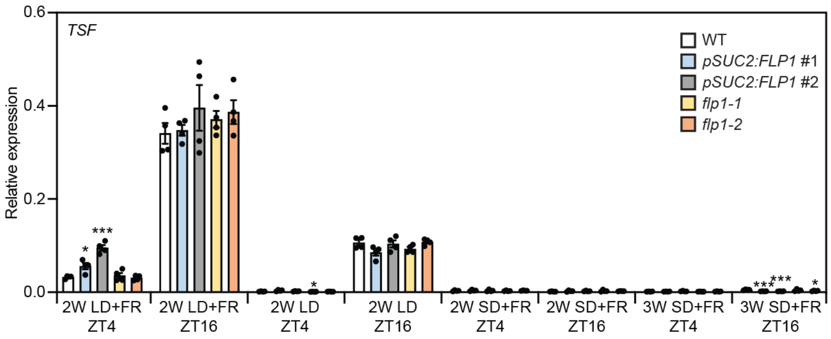


**Figure S10. *TSF* expression in WT, *pSUC2:FLP1,* and *flp1* mutants.**
*TSF* gene expression levels were analyzed in the plants grown under LD+FR, LD, and SD+FR for 2 or 3 weeks and harvested at ZT4 and ZT16. The results represent the means ± SEM. Each dot indicates a biological replicate ($n$ = 4). Asterisks denote significant differences from WT (**P*<0.05; ***P*<0.01; ****P*<0.001, *t*-test).

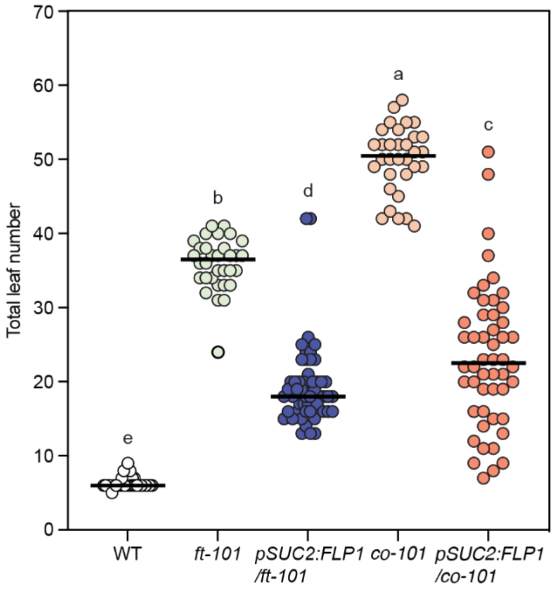


**Figure S11. Flowering time of *pSUC2:FLP1* individual plants in *ft-101* and *co-101* mutant backgrounds at the T1 generation.**
T1 plants were grown on hygromycin selection plates (the plates containing the same growth media except for hygromycin were used for growing WT, *ft-101*, and *co-101* plants) and transferred to soils when grown for 10 days on the plates in LD. Note that WT plants transferred from plates usually flower early than WT plants grown on soil from the beginning. Each dot indicates individual non-transformant and T1 transformants ($n \geq 32$). Bars indicate means, and different letters indicate a statistically significant difference ($P$<0.05, Tukey's test).

WT

```
                                PAM          gRNA
WT 100  GTCTTGGTTTACTTGCCGACCGGTGAAGCCGTCTCATCTTACTCGTCGCT
        ||||||||||||||||||||||||||||||||||||||||||||||||||
WT 100  GTCTTGGTTTACTTGCCGACCGGTGAAGCCGTCTCATCTTACTCGTCGCT
```

Translation: MSGVWVFNNGVIRLVENPNQSGGVSTQSHGRRNVLVYLPTGEAVSSYSSLEQILRSLGWERYFSGDSDLIQYHKRSSIDLISLPRDF SKFNSVYMYDIVVKNPNSFHVRDFN*

*flp1-2 flp2-1*

```
flp1-2flp2-1 100  GTCTTGGTTTACTTGCCGACCGGTGAAAGCCGTCTCATCTTACTCGTCGCT
                  |||||||||||||||||||||||||||  |||||||||||||||||||||||
          WT 100  GTCTTGGTTTACTTGCCGACCGGTGAA-GCCGTCTCATCTTACTCGTCGCT
```

**A** insertion > frame shift and early termination

Translation: MSGVWVFNNGVIRLVENPNQSGGVSTQSHGRRNVLVYLPTGESRLILLVARTNPKEPRVGKILQWRLRSHPVPQTLLHRPHLLTKRL LQVQLRLHVRHRCQEP*

(Wrong amino acids highlighted in green)

*flp1-2 flp2-2*

```
flp1-2flp2-2 100  GTCTTGGTTTACTTGCCGACCGGTGACTGTGTGGACCAAATGAGACCGGTGAATGAGCCGTCTCATCTTACTCGTCGCT
                  ||||||||||||||||||||||||||                             ||||||||||||||||||||||||
          WT 100  GTCTTGGTTTACTTGCCGACCGGTGA-----------------------------AGCCGTCTCATCTTACTCGTCGCT
```

**CTGTGTGGACCAAATGAGACCGGTGAATG** insertion > frame shift and early termination

Translation: MSGVWVFNNGVIRLVENPNQSGGVSTQSHGRRNVLVYLPTGDCVDQMRPVNEPSHLTRRSNKS*

(Wrong amino acids highlighted in green)

**Figure S12. The mutation patterns of the *FLP2* gene in the *flp1-2 flp2-1* and *flp1-2 flp2-2* double mutants generated by genome editing.**

Sequences of guide RNA and PAM in *FLP2* were highlighted with blue and red letters, respectively. The *FLP2* gene was mutated in the *flp1-2* background to generate the *flp1 flp2* double mutants.

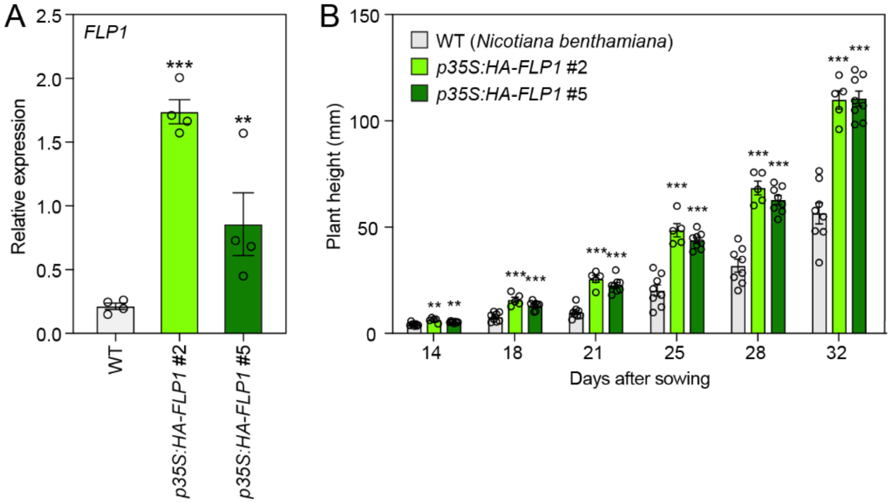


**Figure S13. The effect of overexpression of *Arabidopsis FLP1* in tobacco on the stem growth.**
(A) *FLP1* expression levels in 2-week-old WT (*N. benthamiana*) and *FLP*1 overexpressing tobacco *p35S:HA-FLP1* plants. The results represent the means ± SEM. Each dot indicates a biological replicate ($n$ = 4).

(B) The effect of *FLP1* overexpression on tobacco stem elongation. The results represent the means ± SEM of independent biological replicates. Each dot indicates a biological replicate ($n$ = 5–8). Asterisks denote significant differences from WT (** $P$<0.01, ***$P$<0.001, $t$-test).

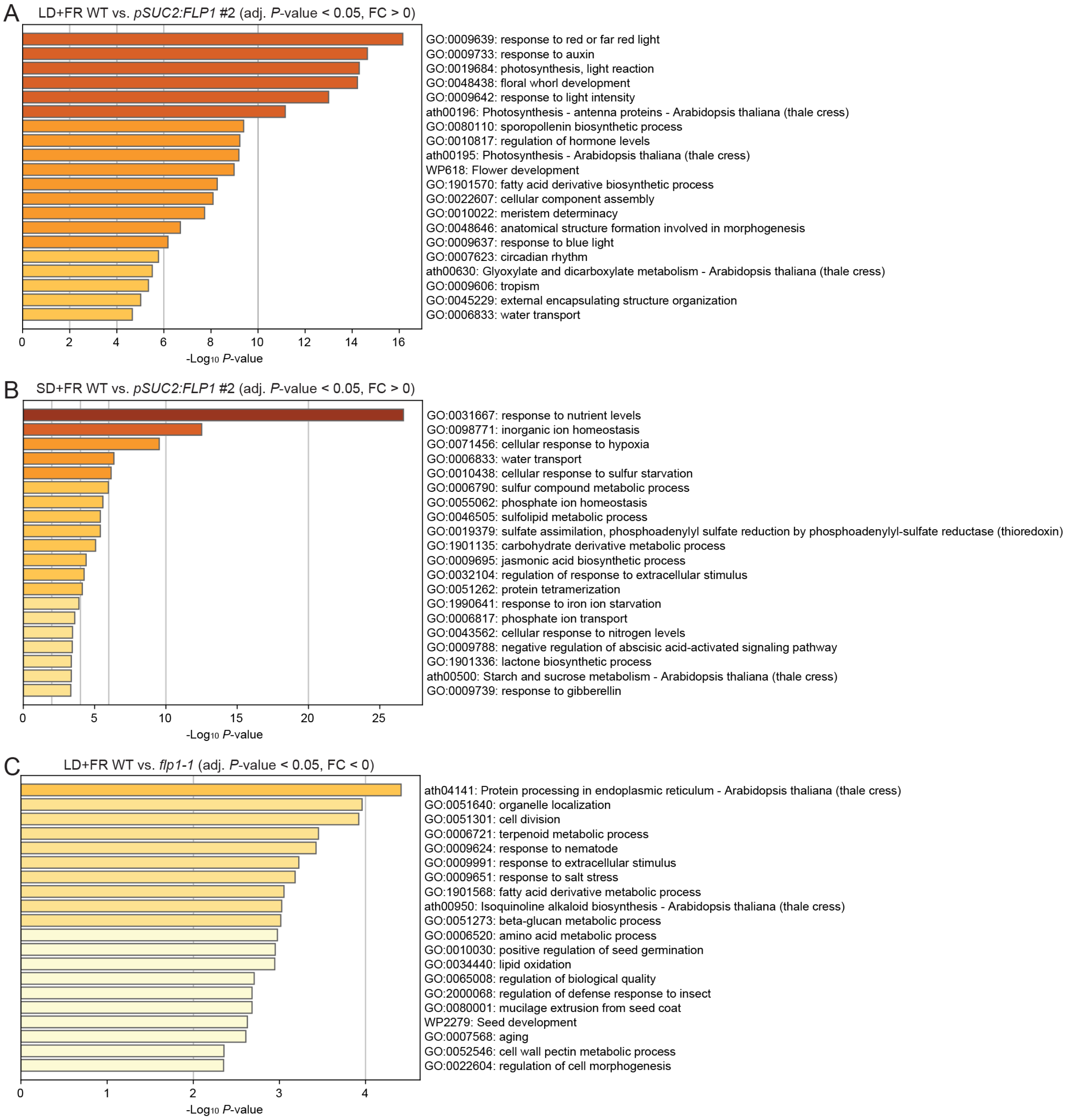


**Figure S14. GO terms enriched in up-regulated genes in *pSUC2:FLP1* #2 and down-regulated genes in *flp1-1* line compared to the genes expressed in WT plants.** Significantly up-regulated genes in *pSUC2:FLP1* #2 in LD+FR (A) and SD+FR (B) and down-regulated genes in *flp1-1* in LD+FR (C) (FDR < 0.05) were used for Metascape enrichment analysis. The numbers of the genes that show the differential expression in each condition are listed in Figure 6A.

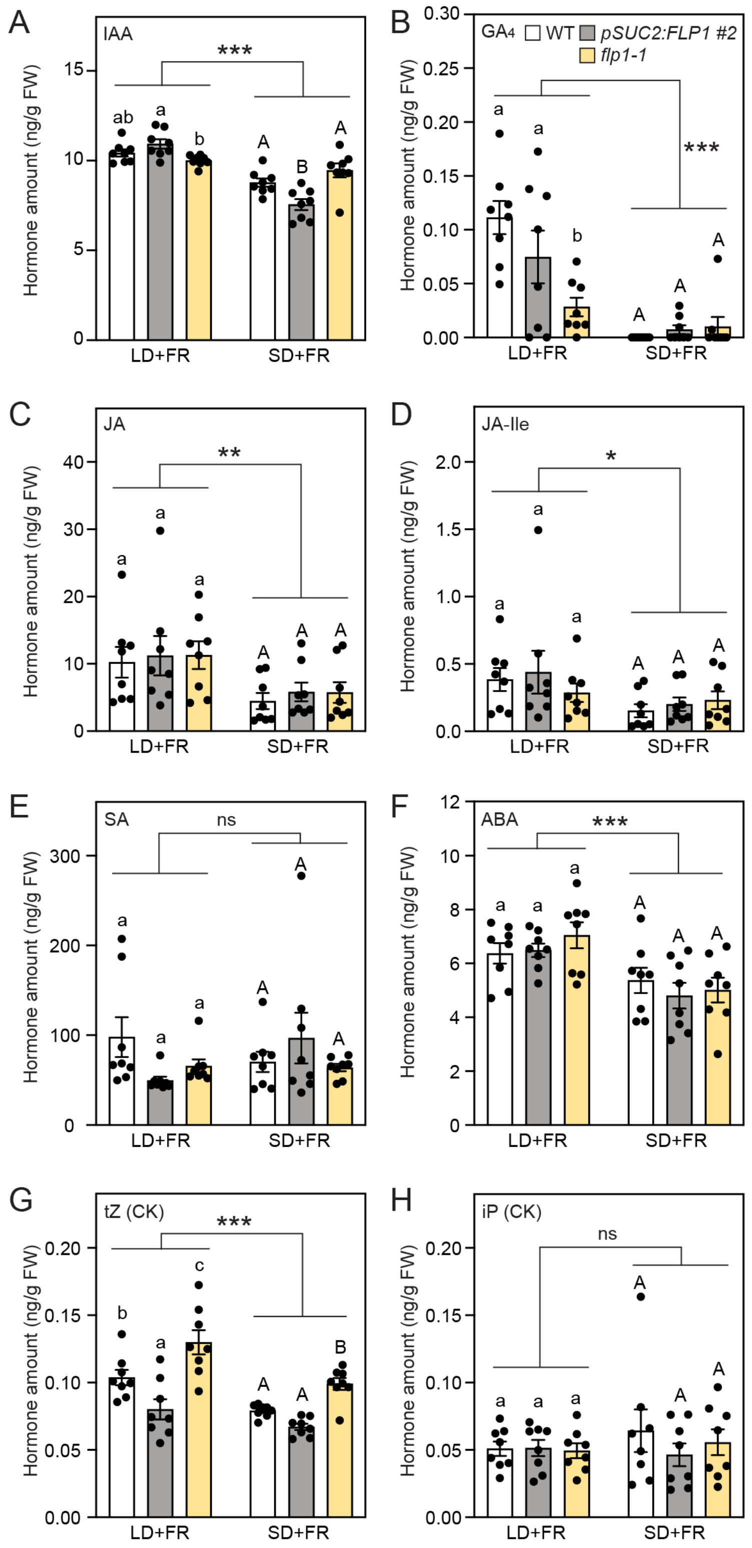


**Figure S15. The effects of *FLP1* expression levels on plant phytohormone contents.** WT, *pSUC2:FLP1* #2, and *flp1-1* plants were grown for 2 weeks in LD+FR and SD+FR.

IAA, indole-3-acetic acid (A); $GA_4$, gibberellin $A_4$ (B); JA, jasmonic acid (C); JA-Ile, jasmonic acid-isoleucine (D); SA, salicylic acid (E); ABA, abscisic acid (F); tZ (CK), trans-zeatin, cytokinin (G); and iP (CK), isopentenyladenine, cytokinin (H). The results represent the means ± SEM. Each dot indicates a biological replicate (*n* = 8). Different lowercase and capital letters indicate statistically significant differences in LD+FR and SD+FR conditions, respectively ($P<0.05$, Tukey's test). Asterisks show significant effects of day length on hormone contents (*$P<0.05$; **$P<0.01$; ***$P<0.001$, two-way ANOVA).

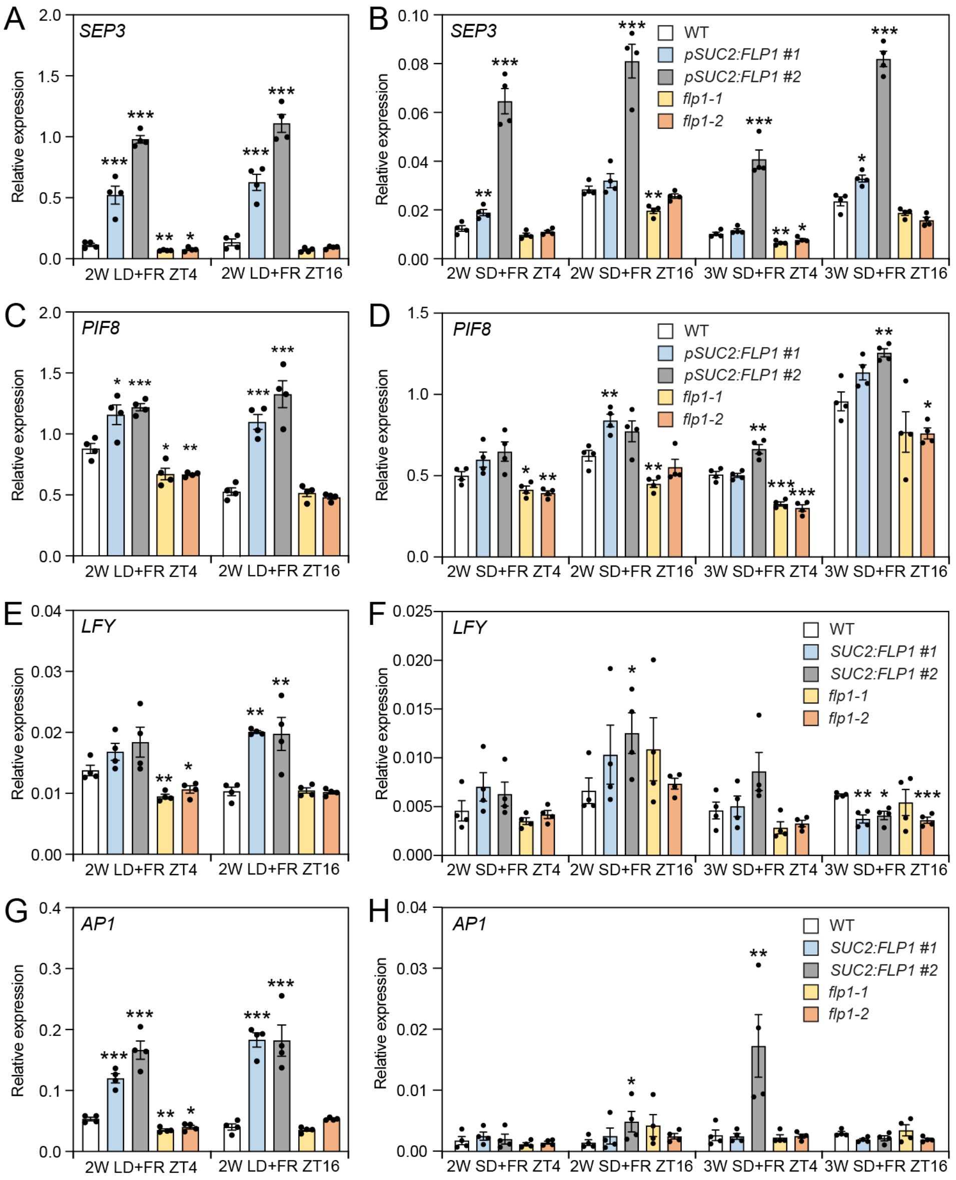


**Figure S16. The effect of *FLP1* levels on the gene expression at 2 to 3-week-old.** *SEP3* (A and B), *PIF8* (C and D), *LFY* (E and F), and *AP1* (G and H) were analyzed using plants were grown under LD+FR or SD+FR conditions for 2 or 3 weeks and harvested at ZT4 and ZT16. The results represent the means ± SEM. Each dot indicates a biological replicate ($n = 4$). Asterisks denote significant differences from WT (*$P$<0.05; **$P$<0.01; ***$P$<0.001, $t$-test).

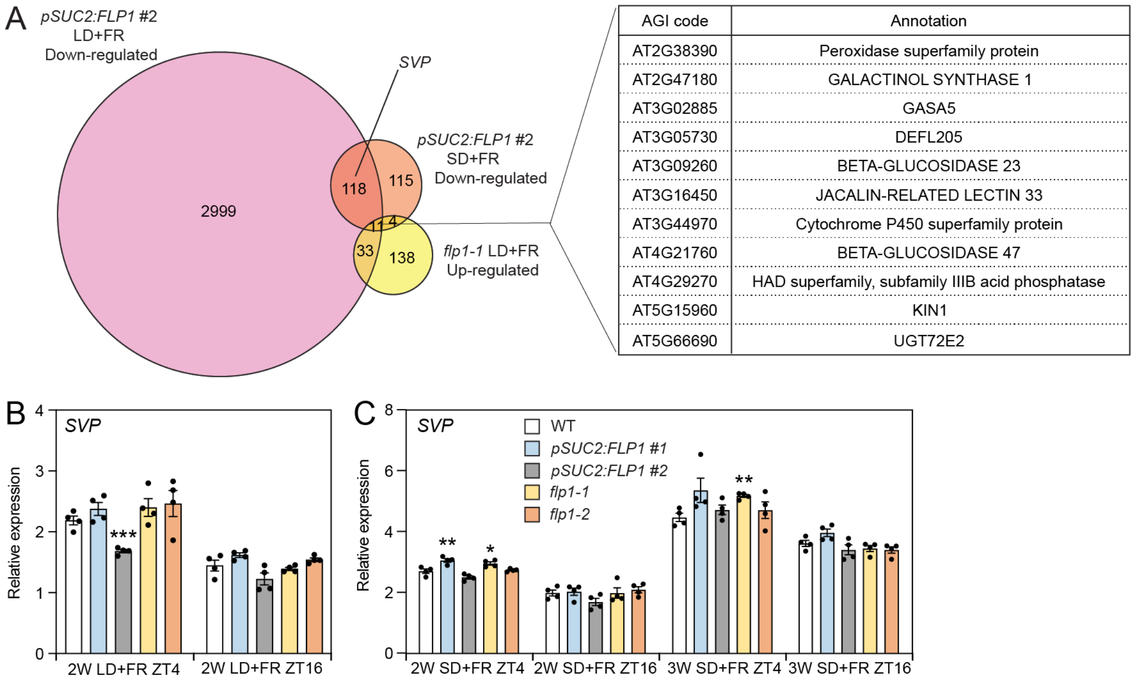


**Figure S17. Genes downregulated by FLP1.**
(A) A Venn diagram consisting of down-regulated genes in 2-week-old *pSUC2:FLP1* #2 grown under LD+FR and SD+FR, and up-regulated genes in *flp1-1* mutant grown under LD+FR conditions. The table on the right lists 11 genes affected by *FLP1* under all three conditions. The category where *SVP* is included is indicated.

(B and C) *SVP* expression levels in 2-week-old plants grown under LD+FR (B) and 2- and 3-week-old plants under SD+FR (C) conditions. The results represent the means ± SEM. Each dot indicates a biological replicate ($n$ = 4). Asterisks denote significant differences from the parental line (**P*<0.05; ***P*<0.01; ****P*<0.001, *t*-test).

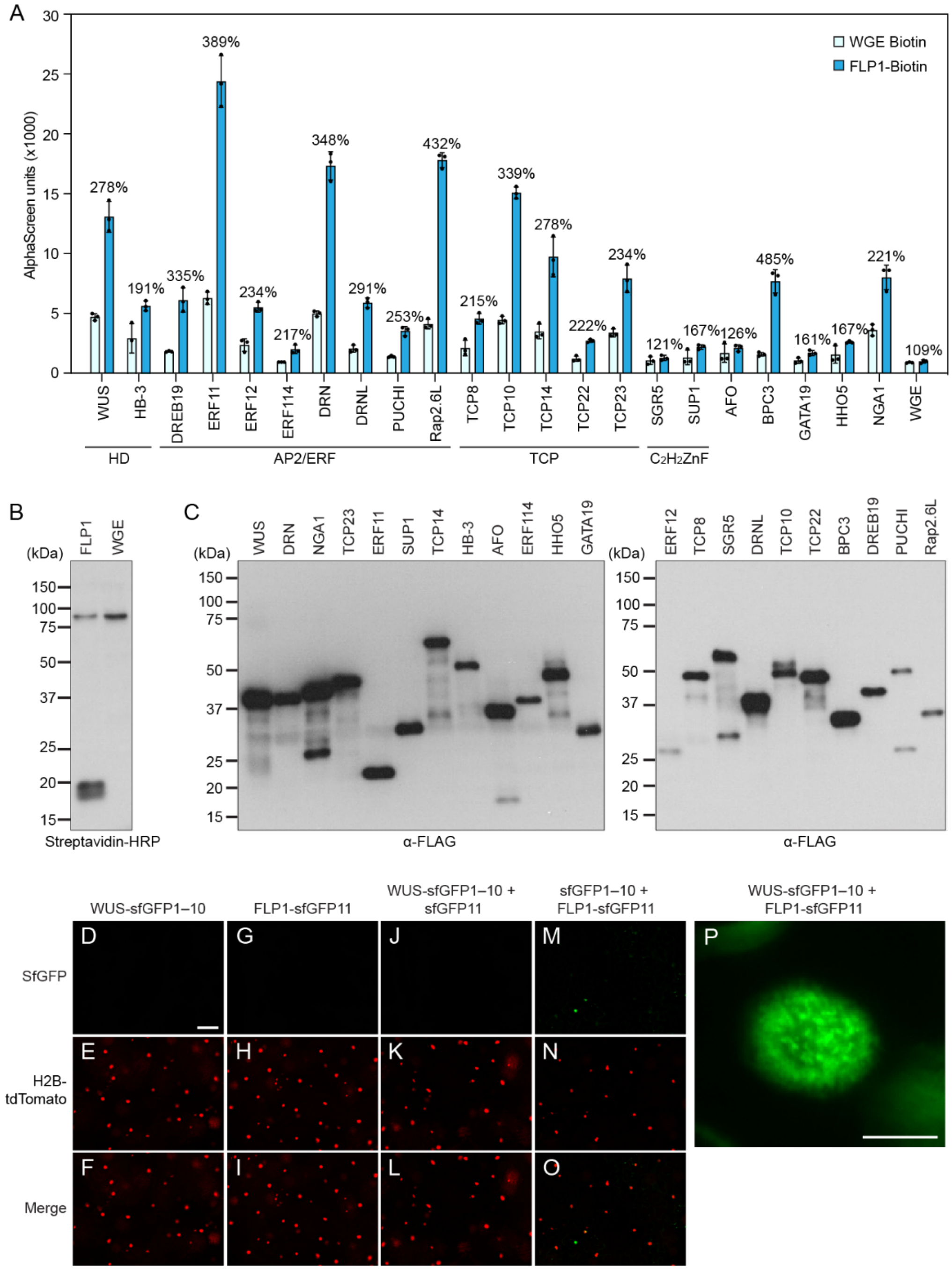


**Figure S18. AlphaScreen and BiFC assay for the protein-protein interaction confirmation between FLP1 and the 22 transcription factors selected from yeast two-hybrid screening results.**

(A) *In vitro* interaction between FLP1 and transcription factors tested using AlphaScreen. Wheat germ extract (WGE) without protein induction was used as a negative control. The results represent the means of three technical AlphaScreen assay replicates ± SD. The percentage indicates a relative difference from the WGE Biotin values.

(B and C) C-terminal biotin-tagged FLP1 (FLP1-Biotin) and N-terminal FLAG-tagged transcription factors were synthesized using the wheat germ cell-free protein synthesis system. The success of protein synthesis of FLP1 (B) and transcription factors (C) was confirmed by western blotting using streptavidin-HRP and anti-FLAG antibodies, respectively.

(D–O) Negative controls for *N. benthamiana* tobacco BiFC assay for FLP1 and WUS interaction shown in Figure 7B–D. WUS-sfGFP1–10 alone (D–F), FLP1-sfGFP11 alone (G–I), and the WUS-sfGFP1–10 and sfGFP11 combination (J–L) did not emit any GFP signals. The combination of sfGFP1–10 and FLP1-sfGFP11 (M–O) sometimes showed patchy and non-nuclear localizing GFP-positive aggregates. H2B-tdTomato marks the positions of nuclei. Scale bar in (D) for all panels D–O, 100 μm.

(P) The speckle-like structure formed by FLP1-sfGFP11 and WUS-sfGFP1–10 in the tobacco nucleus. Scale bar, 10 μm.

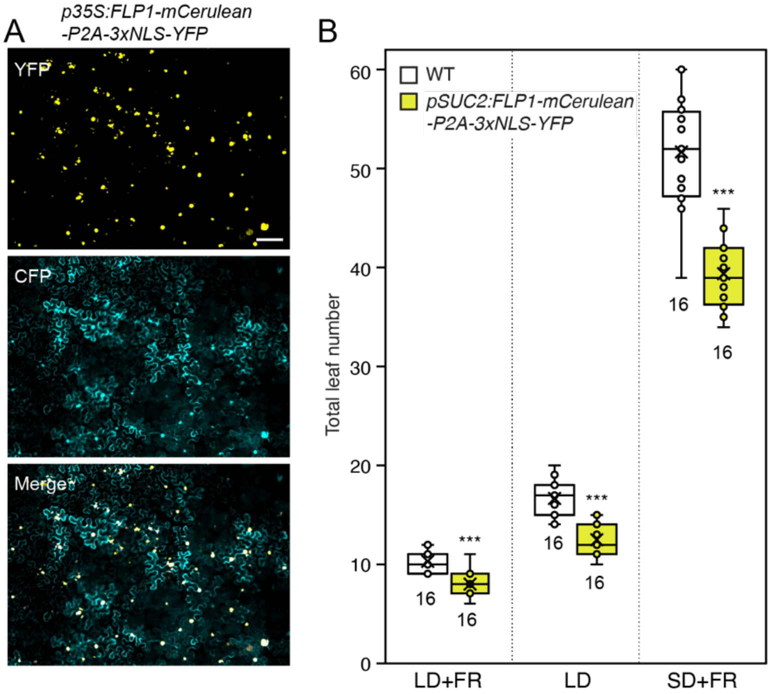


**Figure S19. Characteristics of *FLP1-mCerulean-P2A-3xNLS-YFP*.**
(A) Transient expression of *p35S:FLP1-mCerulean-P2A-3xNLS-YFP* in *N. benthamiana* tobacco epidermal cells. YFP signals were exclusively nuclear localized, while CFP signals were found in both cytosol and nucleus, indicating ribosomal skipping at the *P2A* sequences generated FLP1-mCerulean-P2A and 3xNLS-YFP separately. Scale bar, 100 µm.

(B) The early flowering phenotype of *pSUC2:FLP1-mCerulean-P2A-3xNLS-YFP* line. The bottom and top lines of the box indicate the first and third quantiles, respectively. The bottom and top lines of the whiskers denote minimum and maximum values. Circles indicate inner and outlier points. The bar and the X mark inside the box indicate median and mean values, respectively. The numbers below the box indicate sample sizes. Asterisks denote significant differences from WT (****P*<0.001, *t*-test).
.

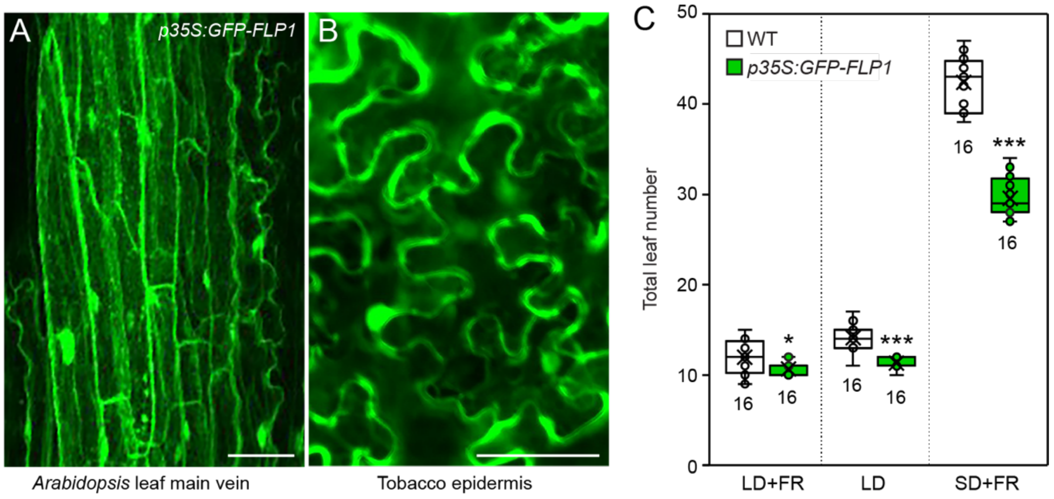


**Figure S20. Characteristics of the *p35S:GFP-FLP1* plants used in grafting experiments.** (A and B) Subcellular localization of GFP-FLP1 in the main leaf vein of the *Arabidopsis p35S:GFP-FLP1* transgenic line (A) and in *N. benthamiana* tobacco epidermal cells transfected with the *p35S:GFP-FLP1* construct using an *Agrobacterium*-mediated transient assay procedure (B). FLP1 protein exists in cytosol and nucleus. Scale bars, 50 µm.

(C) The early flowering phenotype of the *p35S:GFP-FLP1* line. The bottom and top lines of the box indicate the first and third quantiles, respectively. The bottom and top lines of the whiskers denote minimum and maximum values. Circles indicate inner and outlier points. The bar and the X mark inside the box indicate median and mean values, respectively. The numbers below the box indicate sample sizes. Asterisks denote significant differences from WT (*$P$<0.05; ***$P$<0.001, $t$-test).

**Table S1.** qRT-PCR primers used in this study.

| Locus | AGI code | Forward (5'→3') | Reverse (5'→3') |
|---|---|---|---|
| *PP2AA3* | AT1G13320 | GCGGTTGTGGAGAACATGATACG | GAACCAAACACAATTCGTTGCTG |
| *IPP2* | AT3G02780 | GTATGAGTTGCTTCTCCAGCAAAG | GAGGATGGCTGCAACAAGTGT |
| *FT* | AT1G65480 | CTGGAACAACCTTTGGCAAT | TACACTGTTTGCCTGCCAAG |
| *TSF* | AT4G20370 | CTCGGGAATTCATCGTATTG | CCTCTGGCAGTTGAAGTAAG |
| *FLP1* | AT4G31380 | TCCAGTTCCACAAGAGAACTTCG | TCACGGACATGGAAGACGTTAGG |
| *FPF1* | AT5G24860 | CAGGCGTGTGGGTCTTCAA | CTTCTCCTGTCGGTAAATAGACCATCA |
| *FLP2* | AT5G10625 | GAGTGATACGTCTAGTGGAGAA | GAGCGACGAGTAAGATGAGA |
| *SEP3* | AT1G24260 | TATGACGCCTTACAGAGAACC | ATACCCATCAGCTAACCTTAGTC |
| *LFY* | AT5G61850 | ACGCCGTCATTTGCTACTCT | CTTTCTCCGTCTCTGCTGCT |
| *AP1* | AT1G69120 | AGTGGGATCAGCAGAACCAAGGCC | TTGCAGTTGTAAACGGGTTCAAGAGTCAG |
| *SVP* | AT2G22540 | AAGAGAACGAGCGACTTGG | ATACGGTAAGCCGAGCCTAA |
| *RBCS1A* | AT1G67090 | GGCCTCCGATTGGAAAGAAGAAG | GGTGTTGTCGAATCCGATGATCCTA |
| *SUC2* | AT1G22710 | GTGGGAGGTGGACCATTCGACG | CCGGAGGCGGTGAAGGCAAC |
| *GC1* | AT1G22690 | TCGTCCAAGAATCAATTGTGGGC | GTGTTGCCGGAGGTTCCCGG |
| *LHCB2.1* | AT2G05100 | TTGGTGTATCCGGTGGTGGCC | GTCCGTACCAGATGCTTTGAGGAGTAGA |
| *AHA3* | AT5G57350 | GGCTCATGCACAAAGGACTTTACACG | GCGATCTCAGCTCGTCTCTTGGC |
| *Sultr2.1* | AT5G10180 | GGTGTTGAGCTAGTGATCGTTAACCCG | CCCGTAACACAACTGGTCCTTTGA |
| *CER5* | AT1G51500 | AGGAATATCGCTCGAGATGG | TGTCTCCCGAATCCTTTGAG |
| *ML1* | AT4G21750 | CTACTCACAGTTGCGTTTCAGATAC | CCTTCCGAAAACATCGATTAGGCTC |
| *GFP* | None | TGAGCAAGGGCGAGGAGCTG | TGCAGATGAACTTCAGGGTCAGCT |

**Table S2.** Primers used for *in vitro* protein synthesis for the AlphaScreen assay.

| | |
|---|---|
| 1st-FLP1-CF | CACAAAACATTTCCCTACATACAACTTTCAACTTCCTATTATGTCTGGTGTGTGGGTATTC |
| 1st-FLP1-CR | AGTACCTCCCTGCTGGAGACCCATGTCACGGACATGG |
| 1st-WUS-NF | CCAGCAGGGAGGTACTATGGAGCCGCCACAGCATC |
| 1st-ESR1-NF | CCAGCAGGGAGGTACTATGGAAAAAGCCTTGAGAAAC |
| 1st-NGA1-NF | CCAGCAGGGAGGTACTATGATGACAGATTTATCTC |
| 1st-TCP23-NF | CCAGCAGGGAGGTACTATGGAGTCCCACAACAAC |
| 1st-ERF11-NF | CCAGCAGGGAGGTACTATGGCACCGACAGTTAAAAC |
| 1st-SUP-NF | CCAGCAGGGAGGTACTATGGAGAGATCAAACAGC |
| 1st-TCP14-NF | CCAGCAGGGAGGTACTATGCAAAAGCCAACATCAAG |
| 1st-HB-3-NF | CCAGCAGGGAGGTACTATGGCTTCTTCGAATAGAC |
| 1st-AFO-NF | CCAGCAGGGAGGTACTATGTCTATGTCGTCTATG |
| 1st-ERF114-NF | CCAGCAGGGAGGTACTATGTATGGGAAGAGGCCTTTTG |
| 1st-HHO5-NF | CCAGCAGGGAGGTACTATGGTTCAAACAGAAACC |
| 1st-GATA19-NF | CCAGCAGGGAGGTACTATGGGTTTCTCAATGTTC |
| 1st-ERF12-NF | CCAGCAGGGAGGTACTATGGCGTCAACGACGTGTG |
| 1st-TCP8-NF | CCAGCAGGGAGGTACTATGGATCTCTCCGACATC |
| 1st-SGR5-NF | CCAGCAGGGAGGTACTATGAGAACAGATCAAGTG |
| 1st-DRNL-NF | CCAGCAGGGAGGTACTATGGAAGAAGCAATCATG |
| 1st-TCP10-NF | CCAGCAGGGAGGTACTATGGGACTTAAAGGATATAG |
| 1st-TCP22-NF | CCAGCAGGGAGGTACTATGAATCAGAATTCCTCTG |
| 1st-BPC3-NF | CCAGCAGGGAGGTACTATGGAAGAAGATGGATTG |
| 1st-DREB19-NF | CCAGCAGGGAGGTACTATGGAAAAGGAAGATAAC |
| 1st-PUCHI-NF | CCAGCAGGGAGGTACTATGTCAACCTCCAAAACC |
| 1st-Rap2.6L-NF | CCAGCAGGGAGGTACTATGGTCTCCGCTCTCAGC |
| 1st-pENTR/D-TOPO_Uni-NR | CCTTATGGCCGGATCCAAGAGCTCTTTTTTTTTTCTTTGTACAAGAAAGCTGGG |
| 2nd-pENTR/D-TOPO_Uni-NR1 | CCCTCGAAGGATCAGGCCCTTATGGCCGGATCCAA |
| 2nd-pENTR/D-TOPO_Uni-NR2 | GGCCCCCCCTCGAAGG |

**Data S1. (separate file)**
Normalized readcounts of three biological replicates of tissue/cell-specific TRAP-seq samples and its comparison against *p35S:FLAG-GFP-RPL18*.

**Data S2. (separate file)**
List of genes in the clusters in heatmap.

**Data S3. (separate file)**
The effects of red/far-red light adjustment on tissue/cell-specific translatome profiles.

**Data S4. (separate file)**
Genome-wide comparison between WT and *pSUC2:FLP1* #2 grown under LD+FR.

**Data S5. (separate file)**
Genome-wide comparison between WT and *pSUC2:FLP1* #2 grown under SD+FR.

**Data S6. (separate file)**
Genome-wide comparison between WT and *flp1-1* grown under LD+FR.

**Data S7. (separate file)**
Genome-wide comparison between *ft-101* and *pSUC2:FLP1/ft-101* #2 grown under LD+FR.

**Data S8. (separate file)**
List of transcription factors identified as interactors of both FLP1 and FPF1 in yeast two-hybrid assay.
